# Distinct tissue-niche localization and function of synovial tissue myeloid DC subsets in health, and in active and remission Rheumatoid Arthritis

**DOI:** 10.1101/2024.07.17.600758

**Authors:** Lucy MacDonald, Aziza Elmesmari, Domenico Somma, Jack Frew, Clara Di Mario, Roopa Madhu, Audrey Paoletti, Theodoros Simakou, Olympia M. Hardy, Barbara Tolusso, Denise Campobasso, Simone Perniola, Marco Gessi, Maria Rita Gigante, Luca Petricca, Dario Bruno, Lavinia Agra Coletto, Roberta Benvenuto, John D. Isaacs, Andrew Filby, David McDonald, Jasmine P. X. Sim, Nigel Jamieson, Kevin Wei, Maria Antonietta D’Agostino, Neal L. Millar, Simon Milling, Charles McSharry, Elisa Gremese, Karen Affleck, Kenneth F. Baker, Iain B. McInnes, Thomas D. Otto, Ilya Korsunsky, Stefano Alivernini, Mariola Kurowska-Stolarska

**Affiliations:** Research into Inflammatory Arthritis Centre Versus Arthritis (RACE), Glasgow, Birmingham, Newcastle, and Oxford, United Kingdom; School of Infection & Immunity, University of Glasgow, United Kingdom; Immunology Research Core Facility, Gemelli Science and Technology Park, Fondazione Policlinico Universitario A. Gemelli IRCCS, Rome, Italy; Division of Rheumatology, Inflammation, and Immunity, Brigham and Women’s Hospital and Harvard Medical School, Boston, MA 02115, USA; Division of Genetics, Department of Medicine, Brigham and Women’s Hospital, Boston, MA 02115, USA; Broad Institute of MIT and Harvard, Cambridge, MA 02141, USA; Division of Clinical Immunology, Fondazione Policlinico Universitario A. Gemelli IRCCS, Rome, Italy; Institute of Pathology, Fondazione Policlinico Universitario A. Gemelli IRCCS, Rome, Italy; Division of Rheumatology, Fondazione Policlinico Universitario A. Gemelli IRCCS, Rome, Italy; Translational and Clinical Research Institute, Newcastle University, Newcastle upon Tyne, United Kingdom; Musculoskeletal Unit, Newcastle-upon-Tyne Hospitals, Newcastle upon Tyne, United Kingdom; Flow Cytometry Core Facility, Newcastle University, Newcastle upon Tyne, United Kingdom; School of Cancer Sciences, University of Glasgow, United Kingdom; Respiratory and Immunology Research Unit, GSK, Stevenage, United Kingdom

**Keywords:** immune-tolerance, arthritis, disease remission, dendritic cells, synovial tissue

## Abstract

Current rheumatoid arthritis (RA) treatments do not restore immune tolerance. Investigating dendritic cell (DC) populations in human synovial tissue (ST) may reveal pathways to re-instate tolerance in RA. With single-cell and spatial-transcriptomics of synovial tissue biopsies, validated by micro co-culture systems, we identified condition and niche-specific myeloid DC clusters with distinct differentiation trajectories and functions. Healthy synovium contains a unique tolerogenic AXL^pos^ DC2 cluster in the superficial sublining layer. In active RA, a macrophage-rich lining-layer niche becomes populated with inflammatory DC3 clusters that specifically activate memory CCL5^pos^ TEM and CCL5^pos^CXCL13^pos^ TPH, promoting synovitis. In the sublining lymphoid niche, CCR7^pos^ DC2 mReg specifically interact with naïve-T-cells, potentially driving the local expansion of new effector T-cells. Sustained remission sees the resolution of these niches but lacks the recovery of tolerogenic AXL^pos^ DC2, indicating latent potential for disease flare. A human RA disease-flare model showed that the activation of blood predecessor of ST-DC3 clusters precedes the onset of inflammation in joints. Therapeutic strategies targeting pathogenic ST-DC3 clusters, or reinstating tolerogenic AXL^pos^ DC2, may restore immune homeostasis in RA.

**In brief:** Deconstruction of human RA synovium, using single-cell spatial transcriptomics and micro-culture systems, reveals distinct neighbourhoods within the synovial architecture across health, and RA patients with active disease or sustained remission. Discrete niches are identified that contain distinct myeloid DC clusters that differ in frequency, differentiation trajectories, and effector functions.

**Highlights:**

- Human RA synovium exhibits condition and niche specific myeloid DC clusters that vary in their tissue differentiation trajectories and functions.
- ST-CD14^pos^ DC3 (iDC3) support inflammatory CCL5^pos^ TEM and CCL5^pos^ TPH cell activation in the hyperplastic lining layer.
- ST-CCR7^pos^ DC2 (mReg), driven by MIR155, interact with naïve-T-cells in sublining lymphoid niches.
- A specific inflammatory signature of blood predecessors of ST-DC3s predict flare in RA.

**Graphical abstract:** 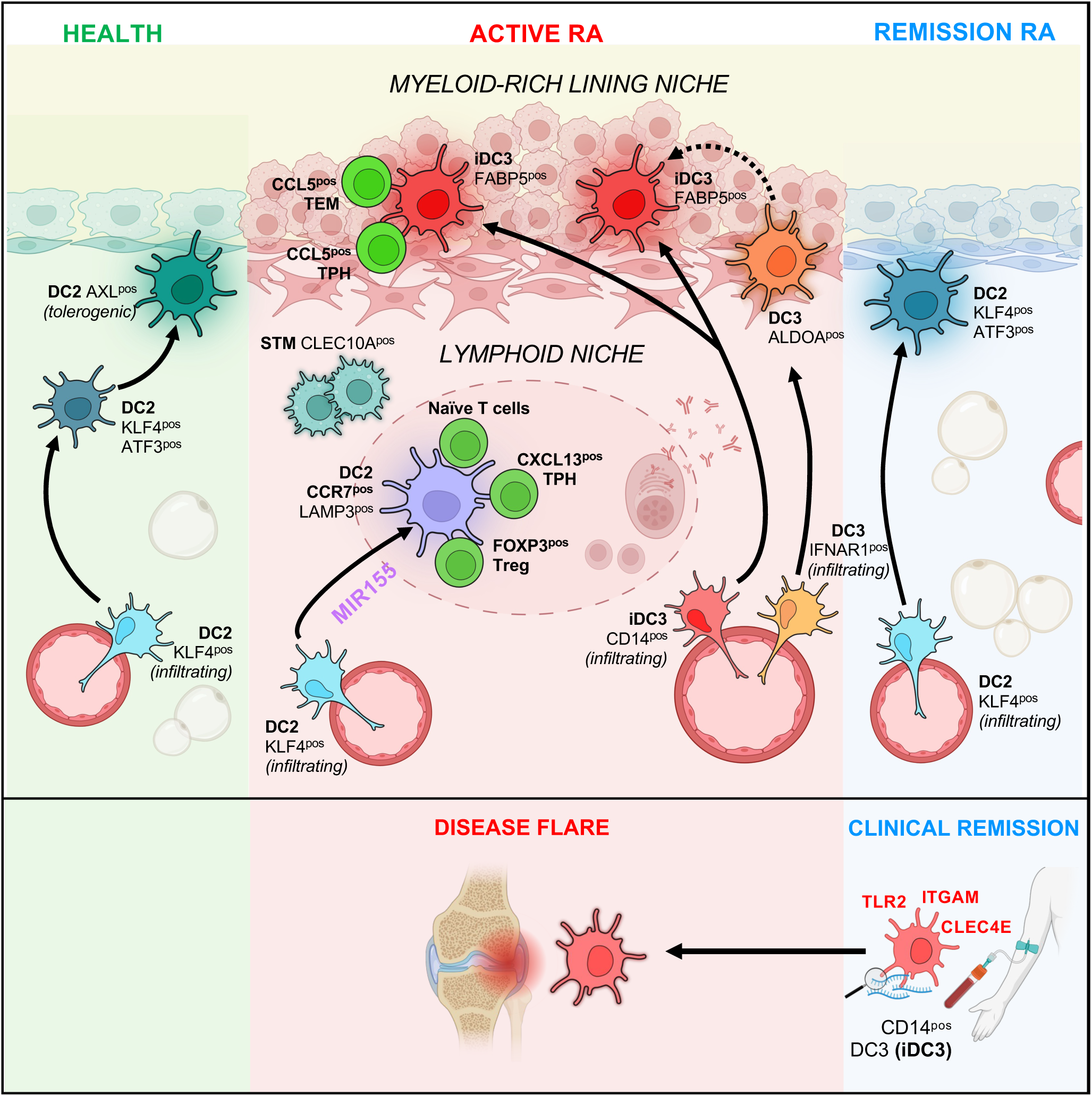

## Introduction

In health, dendritic cells (DCs) direct a protective immune response against pathogens while maintaining tolerance to self-antigens, thus minimising autoimmunity. Self-tolerance is lost in rheumatoid arthritis (RA), associated with the development of citrullinated-peptide specific autoreactive T-cells. These include Th1 effector memory cells (TEM), which drive inflammation^1^, as well as peripheral helper cells (CXCL13^pos^CXCR5^neg^TPH)^2^ and follicular helper T-cells (CXCL13^pos^CXCR5^pos^TFH) ^3^ that help B-cells produce pathogenic anti-modified protein antibodies (AMPA)^4^ in synovial tissue and draining lymph nodes, respectively. AMPA immune-complexes are involved in triggering inflammatory innate^5^ and stromal^6^ synovial cell activation leading to chronic joint pathology and systemic inflammation^7^.

Advances in targeted therapies^8^ have transformed the management of RA. However, 40% of RA patients still do not achieve lasting disease remission, nor is a renewed state of immunological homeostasis achieved^8–10^. Ten to twenty % of RA patients can achieve drug-free remission offering proof-of-concept that pathogenic innate-inflammatory and adaptive responses can be endogenously restrained^9,11^. However, even these patients still exhibit AMPA, demonstrating persistence of autoimmunity and absence of *immunological cure*^12^.

DCs can reset the adaptive immune response and reinstate immune tolerance^13^. Clinical trials with infused autologous tolerogenic DCs generated *ex-vivo* in autoimmune diseases, including diabetes^14^, multiple sclerosis^15,16^, and RA^17,18^ provide promising evidence that DCs have the potential to reinstate peripheral tolerance^14,16,18^. DCs function in peripheral tissues and tissue-draining lymph nodes, forming a dynamic pool sustained by a constant influx of DC precursors from peripheral blood (PB)^13^. Thus, identifying DC phenotypes and their dynamic flow between blood and tissues will inform their roles in the development of autoimmune diseases.

PB DC populations include plasmacytoid DCs (pDCs) and conventional DCs (cDCs). Recent studies utilizing single-cell omics have uncovered heterogeneity within human and mouse conventional DCs in PB and tissues^19–22^. In addition to lymphoid DC1 (CD141^pos^CLEC9A^pos^), two subsets of myeloid DC2 (CLEC10A^pos^) have been identified that differ in their bone marrow progenitors and in their dominant transcription factors, and are now called DC2 and DC3^19–21,23–26^, respectively. DC3 develop from monocyte-DC progenitors (MDP), independently from the common DC progenitor (CDP) restricted DC1/DC2 lineage^23,26^. PB DC2 are CD1c^high^CLEC10A^pos^CD163^neg^CD14^neg^ with a substantial proportion of cells expressing CD5 and BTLA, while PB DC3 are CD5^neg^BTLA^neg^CD1c^low^ and contain different phenotypic clusters that differ in the combination of CD163 and CD14 expression^20,21^. DC3 characterized by high expression of CD163 and CD14 are the most inflammatory, with the greatest potential to activate Th2 and Th17 in vitro^20,27^, to support tumour cytotoxic CD8^pos^CD103^pos^CD69^pos^ tissue resident memory T-cells^21^, and to drive CD4^pos^ immunity in experimental viral respiratory infections^24^. This cluster of DC3 is expanded in PB and kidney tissue in Systemic Lupus Erythematosus (SLE)^20,27^ and will be called, in this manuscript, the inflammatory DC3 cluster (iDC3).

Functional DC atlases of tissue *e.g.*, cancer^21,28,29^, inflamed skin^30^, uveitis^31^ and SLE^20,32^ provide insight into the pathogenic role of discrete DC subsets and their tissue phenotypic states. For example, studies first in cancer^33^ and then other pathologies^29^ led to the fine description of CCR7 and LAMP3 expressing mature DCs enriched in immunoregulatory molecules (mReg DC) as a molecular state (phenotype) acquired by both DC1 and DC2 subsets in tissue, that is associated with the induction of an immunogenic, regulatory and migratory gene program^33,34^ depending on the surrounding environment.

Indirect evidence supports the role of synovial tissue DCs in RA pathogenesis. DC populations reported in synovial fluid/tissue of patients with active RA include plasmacytoid CD123^pos^ DC^35,36^ and conventional CD141^pos^ DC1^37^ and CD1c^pos^ DC2^38–41^, with increased expression of co-stimulatory molecules compared to PB, indicative of maturation. Increased CD1c^pos^ DC2 cells in the T-cell areas of synovial tissue-draining lymph nodes in early RA provides indirect evidence for the potential priming of naïve-T-cells by DCs carrying antigens derived from synovial tissue^42^.

Currently, there is no atlas of the phenotypic and functional diversity of synovial DCs that regulate synovial tissue tolerance or its breach in health, disease, or remission. Herein, we sought to delineate the DC subsets and their phenotypic and functional heterogeneity (clusters) in different neighbourhoods (niches) within human synovial tissue. Using single-cell RNA sequencing (scRNAseq), cellular indexing of transcriptomes and protein epitopes sequencing (CITEseq), and spatial transcriptomics, together with micro-coculture of synovial tissue DC clusters with autologous T-cells, we identified the critical niches that determine DC subset/T-cell subset interactions in RA. These studies establish hierarchies of interaction for naïve-T-cells, TEM, and TPH maturation driven by geographically discrete DC subsets in the pathogenesis of RA, setting the stage for novel therapeutic interventions.

## Results

### Synovial tissue contains distinct myeloid DC subsets and their phenotypic clusters

Our previous single-cell omic studies on synovial tissue macrophages (STMs) and tissue infiltrating monocyes^43^ revealed that, although healthy synovium has relatively low cellularity, it contains a population of myeloid CD1c^pos^CLEC10A^pos^DCs. Using immunofluorescent staining on synovial tissue biopsies, we confirmed the presence of CLEC10A positive (marker of DC2 and DC3), CD68 negative (macrophage marker) DCs in healthy synovium and localized these cells in the superficial sublining layer below the protective lining layer of TREM2^pos^ STMs^43,44^. In active RA, the distorted and hyperplastic lining layer contained an increased number of CLEC10A^pos^CD68^neg^ DCs compared to healthy tissue. In remission, the structure of the lining layer and the number of DCs returned to a state resembling that of healthy individuals (**Figure 1A**).

**Figure 1.**
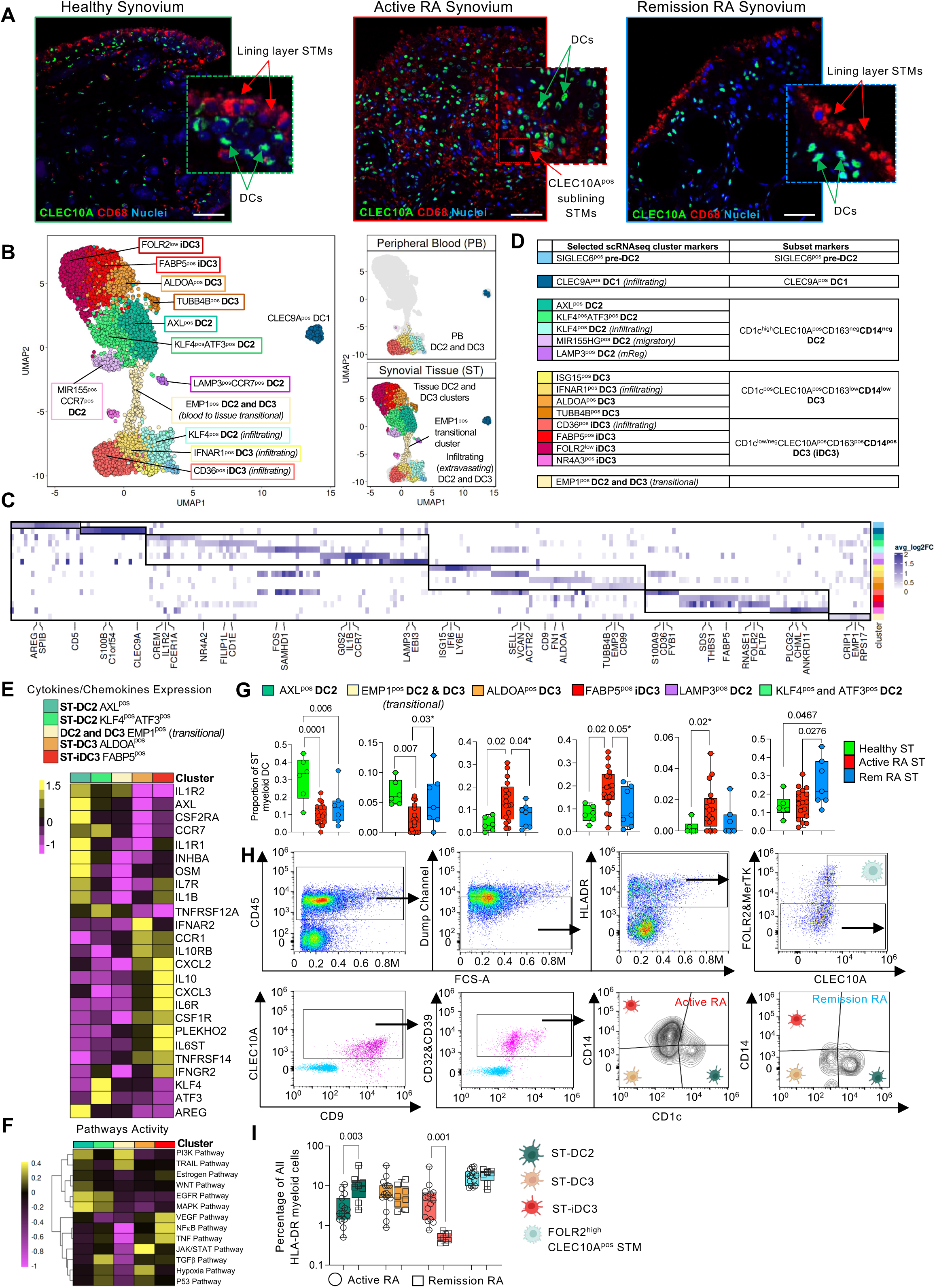
Single-cell omics identified phenotypically distinct clusters of synovial tissue myeloid DC subsets that differ in frequency between health, active RA, and RA in sustained disease remission. **(A)** Representative confocal microscopy images showing IF staining for myeloid DC markers, CLEC10A (green), CD68 (red) and nuclei DAPI (blue) in synovial tissues (ST). The inserts show CLEC10A^pos^CD68^neg^ DCs and CD68^pos^CLEC10A^neg^ lining layer STMs in healthy and RA in remission tissues, and CLEC10A^pos^CD68^neg^ DCs and CD68^pos^CLEC10A^pos^ sublining STMs in active RA tissue at 40x magnification. Images are representative of synovial tissue from healthy donors (n=5), active RA (n=6), and remission RA (n=5). Scale bars = 50μm. Minimum and Maximum display values for CLEC10A were set to 60 and 225 respectively using QuPath (Version 0.4.2). DC; dendritic cells, STM; synovial tissue macrophages. **(B)** UMAP visualization of integrated CITE-seq (n=7) and scRNAseq (n=35) data of myeloid DCs (n=7869) from synovial tissue (6510 cells) and blood (1359 cells). Each cell is represented by an individual point and is coloured by cluster identity based on trajectory analysis from blood precursors (Figure S4). Synovial tissue was obtained from healthy donors (n=7), active RA (n=18), remission RA (n=9), and blood DCs from matched active RA (n=3) and from healthy donors (n=5). **(C)** Heatmap visualizing the average log fold change (logFC) of the top 5 marker genes of ST-DC clusters. Differentially expressed (DE) genes were determined using MAST and were considered significant if expressed in more than 40% of cells in the appropriate cluster with adjusted p-value <0.05 after Bonferroni correction for multiple comparisons. Average log-fold change ≥0.25. The total number of DE genes for each cluster is provided on the right of the heatmap. **(D)** Table summarizing scRNAseq markers of ST-DC subsets and their phenotypic clusters. **(E)** Average-expression heatmap visualising scaled expression of DE genes from the KEGG_cytokine/cytokine receptor pathway and selected cluster markers in ST myeloid DC clusters that differ between joint conditions (Expressed in >25% of cells per cluster, with log-fold change >0.5, and p<0.05 in MAST with Bonferroni correction for multiple comparison) **(F)** Heatmap visualising pathway activity across selected ST-DC clusters as in E. Pathway activity based on top 500 genes per pathway from PROGENy database. **(G)** Proportion of ST DC2, DC3, and iDC3 clusters which differ between healthy controls, active RA, and remission RA based on ST scRNAseq analysis. Data is presented as boxplot with a median and inter-quartile range. One-way ANOVA with Tukey corrections for multiple comparisons, or Two-sided Mann-Whitney (marked with *) if two groups were compared were used. The exact p-values are provided on the graphs. **(H-I)** Multiparameter FACS phenotyping of synovial tissue myeloid DC clusters guided by CITEseq/scRNAseq deconvolution (Figure S12-13). (H) Representative gating strategy for synovial tissue myeloid DC clusters from active RA and RA in remission. First, live cells were gated, followed by CD45-positive cells. In the next step, lineage-positive cells expressing CD3 (T cells), CD19 & CD20 (B cells), CD15 (neutrophils), CD117 (mast cells), and CD56 (NK & NKT cells) were excluded (Dump Channel). Subsequently, cells expressing high levels of HLA-DR were gated. This was followed by gating cells negative for the synovial tissue macrophage markers FOLR2 & MerTK. Cells expressing FOLR2 and CLEC10A are CLEC10A^pos^ STMs. In the FOLR2 & MerTK negative gate, myeloid DCs were gated based on the expression of CLEC10A. The SPP1^pos^ STMs were excluded by the expression of CD9 and lack of expression of CLEC10A. To further exclude any remaining non-DC contamination from myeloid DCs, ST DC2/DC3/iDC3 were confirmed by high expression of CD32 & CD39. The subsequent combination of CD1c and CD14 expression distinguished ST DC2, DC3, and iDC3 in active and remission. (I) Proportion of synovial tissue myeloid DC clusters in total synovial tissue myeloid cells in active (n=15) and remission RA (n=8) presented as boxplot with a median and inter-quartile range. Two-sided Mann-Whitney was used to compare active with remission RA. The exact p-values are provided on the graphs.

To identify DC subsets and their functional tissue phenotypes (clusters) in healthy and in discrete disease states, we performed scRNAseq (with or without CITEseq) on all tissue cells or CD45^pos^ cells from synovial tissue biopsies of healthy donors (n=7), active RA (n=18), and RA in sustained disease remission (n=9). For the purpose of guiding the appropriate annotation of synovial tissue DC subsets/clusters, this was integrated with scRNAseq from matched blood of active RA (n=3) and healthy donors (n=5) (Figure S1A-D), where myeloid DC populations are well-annotated^20,23^ (Figure S2A-B). Briefly, we subsetted PB myeloid cells (all DC and monocyte populations, n=19,784 cells) and integrated them with synovial tissue myeloid cells (broadly annotated as tissue monocytes, macrophages, and dendritic cells, n=50,687) using Harmony (Figure S2C-E). This revealed synovial tissue cells with transcriptomic profiles that cluster together with blood conventional DCs (DC1 or DC2/DC3s) (Figure S2D), suggesting that these are early tissue infiltrating DCs. To identify all synovial tissue myeloid DCs, including those that acquire specific molecular states due to longer residency and activation in tissue, we also retained any other cell cluster that expresses high levels of HLADR and CD11c proteins (highly expressed by conventional DCs), as well as CLEC10A mRNA (marker of myeloid DCs) (Figure S3A-D). This included three additional tissue clusters. The first, based on (i) top myeloid DC2/3 markers (*CLEC10A^pos^, FCER1A^pos^, HLADPB1^pos^, CD1c^pos^)* and (ii) clustering with tissue infiltrating blood DC2/DC3 we temporarily called CLEC10A^pos^NR4A3^pos^CXCR4^pos^ tissue DCs (Figure S3E-G). We found that the other two clusters expressing CLEC10A in synovial tissue are subsets of synovial tissue macrophages: FOLR2^high^LYVE1^pos^, which are perivascular macrophages^43,45^ and FOLR2^high^CLEC10A^pos^ STMs (Figure S3C-D), exhibiting the strongest antigen-presenting cell transcriptomic profile among all STM subsets^43^. We retained the latter as well as the blood/tissue monocyte clusters together with tissue CLEC10A^pos^NR4A3^pos^CXCR4^pos^ DCs and infiltrating DCs in the pool of cells of interest, to exclude contamination of ST-DCs with any monocyte/macrophage cells (Figure S3E-F). Re-clustering of these populations clearly separated transcriptomic profiles of tissue CLEC10A^pos^NR4A3^pos^CXCR4^pos^ DCs, infiltrating myeloid DC clusters and DC1 from monocytes and FOLR2^pos^ macrophages (Figure S3G-I). Subsequent unbiased re-clustering only synovial tissue DCs led to the identification of a small population of DC1 and 14 myeloid DC clusters across healthy, active RA and RA in remission synovium (**Figure 1B-D**, Figure S3J-K and Figure S4). While DC1 were reported to be abundant in RA synovial fluid^37^, they constitute a very small population in synovial tissue and were therefore not the focus of further analysis.

To aid annotation of myeloid DC clusters and to determine which cluster differentiates from which infiltrating PB DC subsets (DC2, DC3 and its iDC3 phenotype), we next performed cell trajectory analysis on those 14 clusters (Figure S4). RNA velocity, inferring differentiation direction from the increased ratio of unspliced to spliced RNA counts (Figure S4A-C), supported by PAGA cell transition confidence evaluation (Figure S4D-F), identified 5, 5, and 4 distinct tissue phenotyping clusters maturing from PB DC2, DC3, and iDC3, respectively (**Figure 1B-D**). All these clusters transcriptionally differ from tissue monocytes and macrophages (Figure S5 and Supplementary material_xls) e.g., by a higher expression of the genes from antigen presentation pathways (Figure S6), including *HLADPB1* found in previous DC single cell studies^20^. Synovial tissues also contained a small population of SIGLEC6^pos^ pre-DC2 (**Figure 1B**). In summary, human synovial tissue contains a rich population of myeloid DCs.

### Healthy synovial tissues exhibit a unique myeloid DC cluster composition that is not restored in remission RA

The 14 myeloid phenotypic clusters of DC2, DC3, and iDC3 **(Figure 1B-D)** exhibited distinctive transcriptomes, characterized by differential expression of between 67 to 955 genes **(Figure 1C and Supplementary material_xls).** Identified clusters of the ST-DC2 subset express high levels of CD1c, lack of CD163 and CD14 and include (i) KLF4^pos^, (ii) KLF4^pos^ATF3^pos^, (iii) AXL^pos^, and two CCR7-positive clusters: (iv) MIR155^pos^ and (v) LAMP3^pos^. Identified clusters of the ST-DC3 subset express CD163, and CD1c at a lower level and include (i) IFNAR1^pos^, (ii) ALDOA^pos^, (iii) TUBB4B^pos^ and (iv) ISG15^pos^. The ST-iDC3 express high levels of CD14 as compared to other DC3s and includes (i) CD36^pos^, (ii) FABP5^pos^, (iii) NR4A3^pos^, and (iv) FOLR2^low^ clusters **(Figure 1D and Figure S7).** Synovial tissue KLF4^pos^ DC2, IFNAR1^pos^ DC3, and CD36^pos^ iDC3 represent an early tissue-infiltrating state of peripheral blood DC2, DC3, and iDC3, respectively, based on their transcriptomic profiles, clustering together with their peripheral blood predecessors **(Figure 1B).** However, they exhibited distinct integrin expression patterns. While PB clusters expressed components of leukocyte-cell-adhesion integrins facilitating cell extravasation from blood, such as VLA4 (*ITGA4 and ITGB1*), their tissue-infiltrating phenotypes downregulated these integrins and showed upregulation of integrins facilitating interactions with the extracellular matrix, such as collagens, fibronectin, and tenascin-C (e.g., *ITGA10, ITGA11, ITGA5, and ITGB8*), thus facilitating post-extravasation migration and maturation within the tissue^46^ **(Figure S8).** In addition, the matched blood and synovial tissue approach enabled the identification of the EMP1^pos^ cluster, which might represent a subsequent transitional molecular state between infiltrating DC2 or DC3 and tissue-niche specific DC phenotypic clusters **(Figure 1B and Figures S4).** This state, in contrast to other clusters, lacks the expression of cytokines and cytokine receptors **(Figure 1E and Figure S8A)** but instead is enriched in the expression of genes encoding EMPs (epithelial membrane proteins) **(Figure 1C and Figure S8C)** and show activation signature of their downstream PI3K pathway **(Figure 1F)**, which are crucial for tumour cell invasiveness and tissue metastasis^47^.

Next, we deconvoluted the frequency of identified tissue myeloid DC phenotypic clusters in different disease states and found substantial differences in AXL^pos^ DC2, EMP1^pos^ transitional DC2 and DC3, ALDOA^pos^ DC3, FABP5^pos^ iDC3, LAMP3^pos^ DC2 and KLF4^pos^ DC2 clusters between health, active RA, and RA in sustained remission **(Figure 1G).** In health, AXL^pos^ DC2 dominate, constituting approximately 40% of all dendritic cells, while all other clusters each constitute less than 5-10% **(Figure 1G),** and represent majority of DCs located under protective lining layer TREM2^pos^ STMs **(Figure S9A).** The AXL^pos^ DC2 cluster is significantly decreased in active RA, especially in difficult-to-treat-RA **(Figure S9B)** and is not reinstated in sustained remission. It exhibits regulatory features as compared to other clusters that includes molecules important for local tissue homeostasis, such AXL, an immune checkpoint that limits adaptive immune response^41,48,49^, amphiregulin (AREG), which is critical for tissue repair ^50^ and *INHBA,* which is known for promoting an immunosuppressive environment^51^. It also exhibits molecular signature reflecting activation of the tolerogenic WNT pathway^52^ **(Figure 1E-F and Figure S10A).** Cell trajectory analysis **(Figure S4D)** showed that this AXL^pos^DC2 cluster likely develops from tissue infiltrating KLF4^pos^ DC2 through an intermediate ST KLF4^pos^ATF3^pos^ phenotype.

Taken together, healthy synovial tissue possesses a unique tissue resident AXL^pos^ DC2 cluster, which differentiates from infiltrating KLF4^pos^ DC2 and exhibits tolerogenic features likely crucial for maintaining local immune tolerance.

### Synovial tissue from active RA is enriched in iDC3 phenotypic clusters that resolve in remission RA

In active RA synovium, we observed an increase in ALDOA^pos^ DC3 and FABP5^pos^ iDC3 phenotypic clusters compared to healthy tissue. Cell trajectory analysis indicated that they likely develop from infiltrating CD14^pos^ DC3 (iDC3) **(Figure S4E).** They together constitute more than 35% of myeloid DC and both shared transcriptomic profiles suggesting activation **(Figure 1E)**. Notably, we observed increased expression of *C15orf48* **(Figure S10E-F)**, a metabolic switch in complex 1 of the respiratory chain underlying pro-inflammatory activation of myeloid cells. We showed previously that mutation in humans leading to permanent replacement of NDUFA4 in complex 1 of the respiratory chain with C15orf48 results in spontaneous production of pro-inflammatory mediators by myeloid cells, including IL-12, IL-6 and TNF^53^. In addition, ALDOA^pos^ DC3 and FABP5^pos^ iDC3 showed cluster-specific activation pathways. ST ALDOA^pos^ DC3s were distinguished from other DC clusters by expression of type I IFN receptor *(IFNAR2*) **(Figure 1E)** and activation of the JAK/STAT pathway **(Figure 1F)** ^25^. ST FABP5^pos^ iDC3 were unique for high expression of the high-affinity IL-6 receptor (*IL6R and IL6ST*) and TNF receptor (*TNFRSF1B*), as well as activation of the TNF and NF-κB pathways **(Figure 1E-F).** We confirmed the presence of both DC3 and CD14^pos^ iDC3 phenotypes in a recently published synovial tissue scRNAseq data set of active RA^54^ **(Figure S11A).**

In the remission synovial tissue, we observed a reduction in ALDOA^pos^ DC3 and FABP5^pos^ iDC3 compared to active RA. However, we noted a lack of restoration of the AXL^pos^ DC2 cluster that characterizes healthy tissue. Instead, we observed an increase in the proportion of its intermediate stages, namely infiltrating KLF4^pos^ and KLF4^pos^ATF3^pos^ DC2 clusters (**Figure 1G**). Transcriptomics of both suggest they progress toward regulatory function in remission tissue (**Figure 1E-F** and Figure S10B-C). For example, the infiltrating KLF4^pos^ cluster expresses *DUSP1* (Figure S10B), a key activator of IL-10^55^, while the ATF3^pos^ cluster expresses *VSIG4*, a PD-L1-like inhibitory molecule that inhibits effector T-cells^56^ (Figure S10C). In addition, the ATF3^pos^ DC2 cluster shows activation of the tolerogenic TGF-beta pathway (**Figure 1F**) and high expression of *ATF3* (**Figure 1E**), which are known for suppressing cytokine gene expression^57^. However, a lack of restoration of AXL^pos^ phenotype potentially suggests an impaired tolerogenic state in remission synovium compared to health.

We next established a specific flow cytometry gating strategy for overall synovial tissue DC2, DC3/iDC3 subsets. We found that canonical PB myeloid DC subset markers, such as CD5 or BTLA for DC2, have limited expression in synovial tissue DCs (Figure S3H and Figure S11A). Thus, our ST-DC subsets specific gating was guided by surface protein expression from CITE-seq data of digested synovial tissue and previously identified markers of STM^43^ (**Figure 1H** and Figure S12). We confirmed the accuracy of the CITE-seq guided gating strategy using single-cell ST-DC index SORT-seq (Figure S13A-C). Briefly, after excluding the majority of tissue resident STMs based on their high FOLR2 and MerTK expression, the synovial tissue myeloid DC populations were distinguished from other ST myeloid cells, such as tissue-infiltrating monocytes and inflammatory STMs, by their expression of CLEC10A, CD32, CD39, and CD9 (details in Methods and Figure S12). The synovial tissue myeloid DC populations of DC2, DC3, and its iDC3 phenotype could then be further distinguished by the expression of CD1c and CD14 (**Figure 1H**). Thus, flow cytometry analysis of additional synovial biopsies from patients with active RA (n=15) and RA in remission (n=8) confirmed that remission tissues predominantly contain ST-DC2s (**Figure 1I**). Active RA is significantly enriched in cells of the CD14^pos^ iDC3 phenotype of DC3 as compared to remission. Other DC3s are present in both active and remission tissue, with a trend towards a higher contribution to the DC pool in active RA, consistent with our scRNAseq data. Similar to our previous study^43^, the relative contribution of high HLADR expressing FOLR2^pos^CLEC10A^pos^ STM macrophages to ST myeloid cells was comparable between active and remission tissues.

In summary, these data identified distinctive distribution of myeloid DC subsets and their specific phenotypic clusters in human synovium associated with homeostasis, and with inflammation and remission in RA.

### The LAMP3^pos^CCR7^pos^ DC2 (mReg) cluster is enriched in active RA synovium

While iDC3 clusters dominate the pool of ST myeloid DCs in active RA, the synovial DC2 population is still present and shows significant changes in cluster composition compared to healthy individuals and those in disease remission. The AXL-positive DC2 cluster is reduced, while the CCR7-expressing DC2s are increased compared to healthy individuals and those in remission from RA **(Figure 1G).** We identified two synovial tissue CCR7*-*positive DC2 clusters (**Figure 2A).** Their high CCR7 expression suggests the potential for migration into draining lymph nodes (DLN), where their distinct high expression of the co-stimulatory molecule *CD40* would enable them to activate naïve-T-cells. They also share high expression of NF-κB pathway genes (e.g., *REL, TRAF1, NFKB1*), suggesting an activated state **(Figure 2A-B).** The two CCR7^pos^ clusters were distinguished by their levels of *MIR155* and *LAMP3* expression. One cluster expresses high levels of *MIR155*, while the other exhibits lower levels of *MIR155* and high levels of *LAMP3* **(Figure 2A).** MIR155 amplifies myeloid cell activation by inhibiting negative regulators of TLR/cytokine receptor signalling^58^. Consistent with this, high expression of pro-inflammatory mediators such as *IL1*β and multiple chemokines were observed in the CCR7^pos^MIR155^pos^ DC2 cluster **(Figure 2B)**. In contrast, the LAMP3^pos^ cluster exhibit *IL12*, *IL-15* and *IL-32,* together with increased expression of regulatory genes such as *EBI3* (encoding a subunit of regulatory cytokines IL-27 and IL-35) and the immunomodulatory enzyme *IDO1*, the product of which is key for the differentiation and activation of T-regs **(Figure 2B),** thus resembling the recently identified mReg DC phenotype^33^. To investigate this, we integrated our synovial tissue single-cell data set with a myeloid cell (MNP) single-cell RNA compendium (MNPVerse)^29^ **(Figure 2C and Figure S11B-D).** We detected a specific cluster that corresponds to mReg DC from MNPVerse and ST CCR7^pos^LAMP3^pos^ DC2 **(Figure S11B-D).** Among all MNPVerse myeloid cells, the ST CCR7^pos^LAMP3^pos^ DC2 signature scored highest with a MNPVerse mReg signature **(Figure 2C),** confirming that ST CCR7^pos^LAMP3^pos^ DCs are indeed mReg. mReg are characterized by a unique immunogenic, regulatory, and migratory gene programme^33,59^. We found that these gene programs were expressed higher in the LAMP3^pos^ compared to the MIR155^pos^ cluster. Additionally, while the immunogenic program was increased in the LAMP3^pos^ DC2 cluster in both active and remission RA compared to health, the expression of some immunoregulatory genes such as immune checkpoints (PDL-1 [CD274], PDL-2 [PDCD1LG2], and CD200) was increased in cells from remission tissue compared to those with active RA **(Figure 2D).** This suggests a distinct immunogenic versus regulatory program in active and remission RA, respectively.

**Figure 2.**
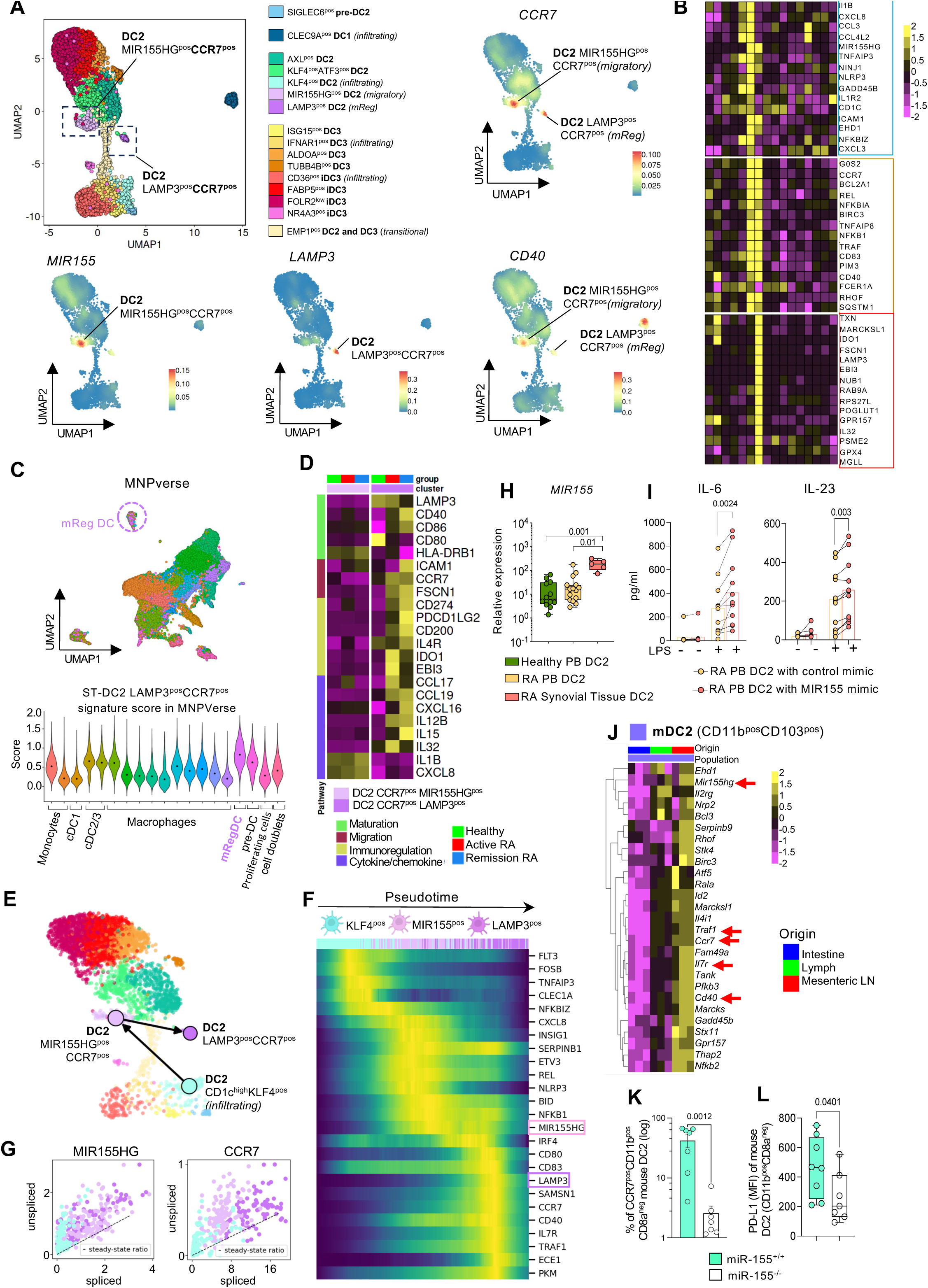
Synovial tissue mReg DCs (LAMP3^pos^CCR7^pos^ DC2) differentiate from activated MIR155^pos^CCR7^pos^ intermediates, and this process is driven by MIR155. **(A)** UMAP visualization of integrated CITE-seq (n=7) and scRNAseq (n=35) data of myeloid DCs from synovial tissue of healthy, active RA and RA in disease remission as in Figure 1. Lines indicate two most-highly CCR7 expressing clusters MIR155^pos^ and LAMP3^pos^. Density plots illustrate expression of CCR7, MIR155, LAMP3 and CD40 in UMAPs of integrated PB/ST myeloid DC data set as in Figure 1. **(B)** Heatmap visualizing scaled top 15 unique marker genes of DC2 MIR155^pos^CCR7^pos^ (blue box) and DC2 LAMP3^pos^CCR7^pos^ (red box) clusters, and the top 15 shared marker genes of these clusters (orange box) as compared to any other ST-DC cluster. Selection criteria: expressed >40% of cells in a cluster with log-fold change >0.25 and p<0.05 MAST with Bonferroni correction for multiple comparison. **(C)** Gene-set module score of ST DC2 LAMP3^pos^CCR7^pos^ clusters (computed from their unique differentially expressed genes, Methods) plotted across MNPVerse dataset clusters (Mulder et al, Immunity 2021). **(D)** Heatmap showing distinct expression of genes linked to mReg pathways identified by Maier et al. Nature 2020 in ST MIR155^pos^CCR7^pos^ and LAMP3^pos^CCR7^pos^ clusters between different joint conditions. **(E)** Single-cell RNA Velocity-directed PAGA revealed the differentiation trajectory of ST-DC2 LAMP3^pos^CCR7^pos^ from tissue infiltrating DC2 through the intermediate MIR155^pos^CCR7^pos^ ST-DC2 stage. Analysis based on single-cell omics data from ST of active RA (n=17). **(F)** Pseudotime analysis of data from panel E shows candidate genes responsible for the maturation trajectory of ST-DC2 LAMP3^pos^CCR7^pos^ from MIR155^pos^CCR7^pos^ ST-DC2. Cells are coloured by cluster identity and by pseudotime. **(G)** The ratio of newly transcribed (unspliced) pre-mRNA to spliced mRNA of MIR155 and CCR7 indicates increasing transcription activity from KLF4^pos^ DC2 towards LAMP3^pos^ DC2. **(H)** Relative expression of MIR155 as compared to housekeeping microRNA (qPCR) in total CD1c^high^ DC2 population isolated from PB of Healthy donors (n=12), PB of RA patients with active disease (n=16), and from synovial tissues of RA patients with active disease (n=5). Data are presented as boxplot with a median and inter-quartile range. One-way ANOVA with Tukey corrections for multiple comparisons. The exact p-values are shown on graph. **(I)** Blood CD1c^high^ DC2 were FACS-sorted from active RA patients (n=11) and transfected with miR-155 or control mimic (20 nM). These were unstimulated or stimulated with LPS (100 ng/mL), and in vitro synthesis of IL-6 and IL-23 were quantified after 48h by ELISA. Data are presented as paired dot-plot with mean bars; each dot represents one patient. A two-sided, paired t-test was used, and p-values are shown on the graph. **(J)** Heatmap shows log-normalized expression values of ST-DC2 LAMP3^pos^CCR7^pos^ ^cluster^ markers significantly upregulated in bulk RNAseq of the mouse CD11b^pos^CD103^pos^ myeloid DC2 cluster in murine mesenteric LN and Lymph versus intestine tissue (n=3 mice) with p-value ≤ 0.05 (DESeq2 with Bonferroni correction). To collect migratory DC in lymph, the thoracic lymph duct was cannulated, after prior removal of the mLN, as described in (Mayer et al, 2020). Red arrows point to the genes of interest (mReg gene score). **(K)** The percentage of CCR7 expressing mouse DC2 (CD11b^pos^CD8a^neg^) was evaluated in mesenteric lymph nodes (mLN) of WT (miR-155+/+, n=7) and miR-155 deficient mice (miR-155-/-, n=7). Data are presented as scatter dot plot with mean and SEM. A two-sided Mann-Whitney test was used, and exact p-values are provided on the graph. **(L)** The MFI of PD-L1 expression by DC2 (CD11b^pos^CD8a^neg^) in mesenteric lymph nodes (mLN) of WT (miR-155+/+, n=8) and miR-155 deficient mice (miR-155-/-, n=7). Data is presented as boxplot with a median and inter-quartile range. Two-sided Mann-Whitney, the exact p-values are provided on the graphs.

To explore the relationship between MIR155^pos^ and mReg clusters in synovial tissue, we performed cell trajectory analysis. RNA velocity **(Figure S4E),** aided by PAGA cell transition confidence evaluation **(Figure 2E)**, indicated that the ST CCR7^pos^MIR155^pos^ cluster may constitute an intermediate stage of mReg DC2 differentiation from tissue infiltrating KLF4^pos^ DC2s.

Next, to explore molecular mechanisms underlying the differentiation from infiltrating KLF4^pos^ DC2 to mReg phenotype, we identified genes **(Figure 2F-G)** with high ratio of unspliced transcripts and differentially expressed (DE) along the pseudotime of cell progression through the trajectory identified in **Figure 2E**. This highlighted MIR155 as (i) a potential driver of the highly proinflammatory state of the MIR155^pos^ cluster and (ii) a candidate regulator of the migratory program of LAMP3^pos^ mReg. MIR155 expression is increased in the RA ST-DC2 cells as compared to PB counterparts **(Figure 2H)**, suggesting its primary role in the local tissue maturation of infiltrating DC2 into an inflammatory MIR155^pos^ state. *In vitro* overexpression of MIR155 in PB DC2 from active RA significantly increased their production of IL-6 and IL-23 **(Figure 2I),** suggesting that MIR155 might govern the stimulatory functions of the CCR7^pos^MIR155^pos^ intermediate cluster in the synovium of patients with active RA.

Access to lymph and lymph nodes DCs in humans is challenging. To, as closely as possible, validate the migratory properties of the mReg and MIR155^pos^ clusters, we took advantage of a well-characterised mouse gut lymphatic cannulation model. We FACS-sorted mouse DC2 (CD11b^pos^CD8a^neg^), after exclusion other DCs and myeloid cells from intestinal tissue, matched lymph collected by cannula and from mesenteric LNs, as described previously^60^ and estimated the presence of mReg/MIR155^pos^ signatures across these different compartments. The mReg/MIR155^pos^ cluster markers, including high expression levels of *Ccr7, mir155*, *Cd40,* NF-κB pathway (*traf1*), were significantly increased in lymph, and particularly in DLN, as compared to the tissue, confirming the LN migratory properties of these clusters **(Figure 2J).**

Along the mReg differentiation trajectory, high MIR155 expression appears before CCR7 expression and other markers of the mReg phenotype **(Figure 2F-G).** To explore the specific role of MIR155 in the mReg migratory program, we investigated CCR7^pos^ DC2 cells in the mesenteric LN of WT and MIR155-deficient mice. We found a significant decrease (∼80%) in the proportion of CCR7^pos^ DC2 in MIR155-deficient mice as compared to WT littermates **(Figure 2K)**. These MIR155-deficient DC2 cells also had significantly less expression of another marker of mReg, PD-L1 **(Figure 2L)**, confirming a potential key role of MIR155 in the regulation of mReg phenotype.

Taken together, these data suggest that active RA synovium contains an immunogenic mReg cluster, which differentiates from infiltrating DC2 through the highly activated CCR7^pos^MIR155^pos^ intermediate stage and likely migrates to draining lymph nodes to initiate a peripheral autoimmune response.

### Different myeloid DC subsets localize in distinct synovial tissue niches

To elucidate the function of distinct myeloid DC subsets/clusters in tissue, we next mapped their localization in synovial tissue niches and evaluated their interactions with specific T-cell clusters. We performed single-cell spatial transcriptomics (using a 980-gene panel, CosMx) on synovial tissue biopsies from patients with active, treatment-naïve RA (n=3) and RA in disease remission (n=3), spanning 36 and 16 fields of view (FOVs) respectively, across histologically defined lining and sublining layers **(Figure S14-16).** To accurately identify cells within active and remission tissue, we firstly performed initial image segmentation with Mesmer^61^ to estimate cellular boundaries followed by refinement based on transcript densities with Baysor^62^. We then performed reference annotation of single-cell spatial data by integration with our single-cell sequencing data for those tissues **(Figure 3A-B)**. The marker gene correlation between reference scRNAseq myeloid clusters and those from the integration of spatial and scRNAseq data, and a confusion matrix demonstrating the contribution of scRNAseq-identified cell clusters within the new integrated scRNAseq/spatial data clustering, confirmed our ability to confidently map ST-DC down to the subset level (DC1, DC2, DC3 and its iDC3 phenotype) and to the CCR7^pos^DC2 (MIR155/mReg) cluster level **(Figure S14B-C).**

**Figure 3.**
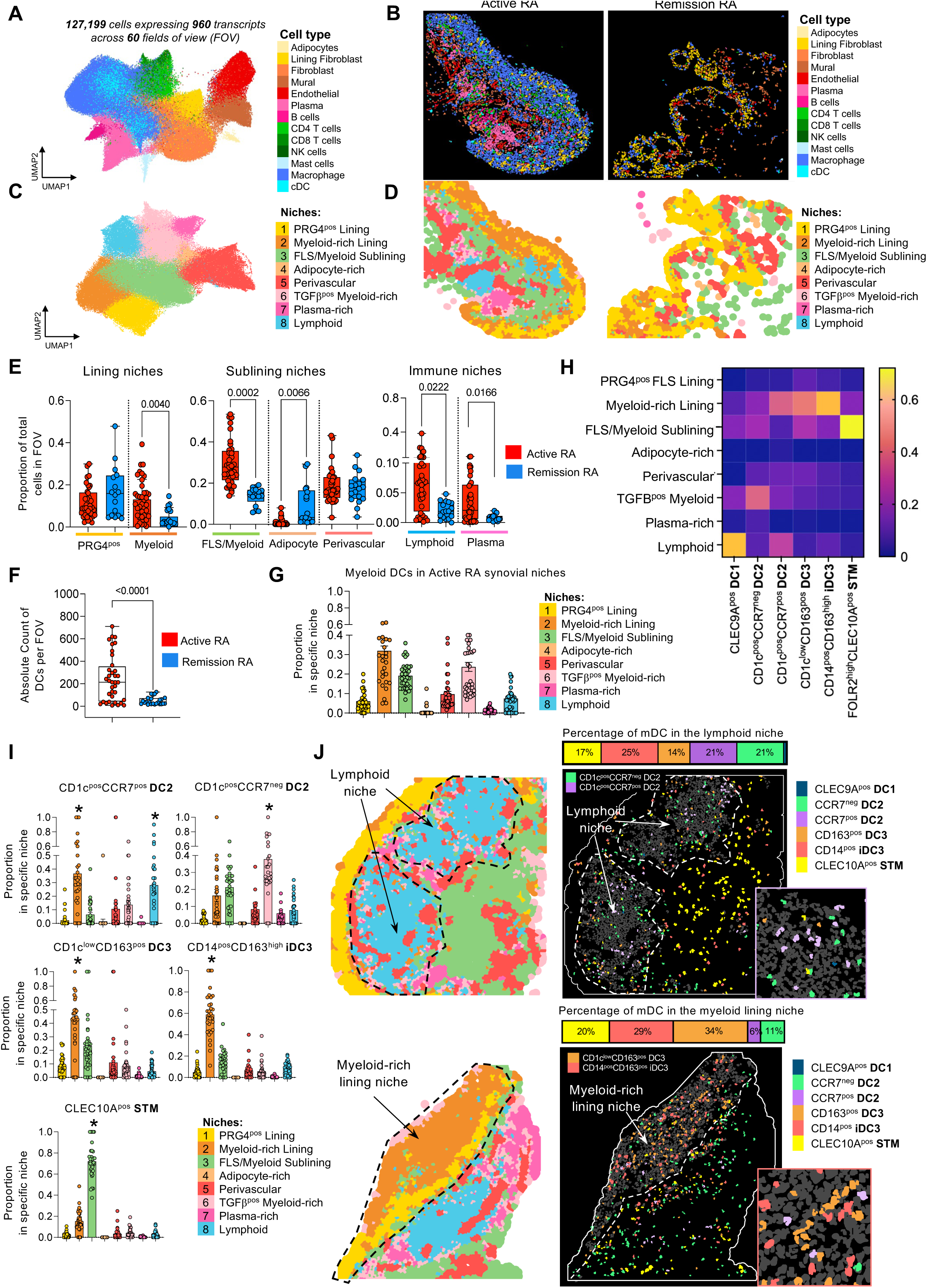
Different myeloid DC clusters localize in distinct synovial tissue niches. **(A)** UMAP visualization of clustering of coarse cell types identified from single-cell spatial transcriptomic data (CosMx, 960 gene panel) of 127,199 cells from n=69 fields-of-view (FOV) from synovial tissue of active RA (n=3) and RA in remission (n=3). **(B)** Illustration of tissue single cell segmentation and coarse cell type annotation of representative FOVs from active RA and remission RA synovium. **(C)** UMAP visualization of Louvain clustering of tiles from Voronoi tessellation of the tissue (details Methods). Clusters represent regions (niches) from tissue segmentation that are annotated based on coarse cell type composition and differential gene expression More details in Figure S14-15. **(D)** Illustration of niche annotation of representative FOVs from active RA and remission RA synovium. **(E)** Boxplots with a median and inter-quartile range show the proportion of total cells in each FOV per identified niche in active RA (n=36 FOVs, n=3 patients) and in remission RA (n=16 FOVs, n=3 patients). Two-sided Mann-Whitney, the exact p-values are provided on the graphs. **(F)** Absolute number of myeloid DC clusters among all spatially mapped cells in active RA and Remission FOVs (n=36 in active and n=16 in remission tissues). Each dot represents one FOV. Data is presented as boxplot with a median and inter-quartile range. Two-sided Mann-Whitney, the exact p-values are provided on the graphs. **(G)** Distribution of myeloid DC clusters, DC1 and CLEC10A^pos^ STMs in different niches of active RA synovium. Each dot represents one FOV (n=36 active RA). Data are presented as the proportion of myeloid DCs per niche (Bar plots with SEM). **(H)** Heatmap summarizing the proportion of each myeloid DC cluster, DC1 and CLEC10A^pos^ STM per niche. **(I)** Proportion of cells of selected myeloid DC clusters per niche (Bar plots with SEM). * p<0.01 compared to any other niche (One-way ANOVA with Dunnett’s corrections for multiple comparisons). **(J)** Illustration of distinct distribution of ST DC2 and DC3 and iDC3 clusters in lymphoid and lining myeloid-rich niches. The stack bar on the side shows the proportion of all myeloid DC clusters in those niches.

Next, to pinpoint myeloid DC localization, we sought to identify regions of synovial tissue characterised by distinct cellular neighbourhoods. Using neighbourhood analysis (Voronoi tessellation & Louvain clustering, details in Methods), we identified 8 distinct synovial tissue neighbourhoods (niches) (**Figure 3C-D**). Each contained a distinct distribution of cell types and exhibited a unique transcriptional signature indicative of their local cell composition/activation (Figure S15-16)^63^. These comprise two lining layer niches: (i) ***PRG4^pos^ niche***, dominated by PRG4-expressing lining layer fibroblasts (FLS), and (ii) the ***myeloid-rich lining***, dominated by macrophages. Three structural sublining niches were identified and include (i) enriched in interstitial macrophages and FLS (***FLS/Myeloid sublining niche***), (ii) adipocytes (***adipocyte-rich niche***), and (iii) ***perivascular niche*** enriched in endothelial cells and pericytes. Three immune niches were identified: (i) enriched in T and B cells (***lymphoid niche***), that represent previously described in RA synovium ectopic germinal centres^7,64^, (ii) plasma cells (***plasma-rich niche***), and (iii) a niche enriched in regulatory TGFβ-expressing myeloid cells (***TGF***β***-positive myeloid-rich***). Active RA contains a larger myeloid-rich lining niche, lymphoid niche, and plasma-rich niche compared to remission. Conversely, remission exhibits a larger adipocyte-rich niche compared to active RA **(Figure 3E).**

Next, we investigated the distribution of different DC subsets within niches. We mapped 8887 DCs in the tissue of active RA, with a median of approximately 213 DCs per field of view (FOV). Due to the relative low cellularity of remission FOVs, we were only able to map 698 DCs across all remission FOVs, with a median number of 24 per FOV **(Figure 3F).** Therefore, in further analysis, we focused solely on the distribution of DC subsets within active RA niches. We found that DCs are particularly enriched in myeloid-rich lining, TGFβ^pos^ myeloid-rich sublining, myeloid/FLS sublining, and lymphoid niches **(Figure 3G)** and show subtype-specific distribution **(Figure 3H).** Thus, we found that CCR7^pos^ DC2 (mReg/MIR155) were significantly enriched in the lymphoid niche, while the remaining DC2 (CCR7-negative) were in the TGFβ^pos^ myeloid-rich sublining niche. DC3 and its iDC3 phenotype were significantly enriched in the myeloid-rich lining niche compared to any other tissue niche. **(Figure 3I-J).** In summary, ST-DC subsets exhibit tissue niche-specific distribution, suggesting unique local functions in regulating adaptive immunity.

### Distinct T-cell subtypes in tissue niches

To infer the function of distinct DC subsets in active RA, we first needed to map specific CD4^pos^ T-cell synovial clusters. To achieve this, we first deconvoluted synovial tissue T-cell clusters in RA across different disease stages, spanning from naive-to-treatment RA to difficult-to-treat RA, and RA in disease remission, using single-cell omics **(Figure 4A and Figure S17).** Integration of these data with a dataset from matched blood aided in cluster deconvolution and allocation of naive versus memory status to the T-cells. Among T-cells, we identified twelve distinct CD4^pos^ T-cell clusters: two naïve-T-cell clusters and ten memory T cells, including two central memory (TCM) clusters, six T-effector memory (TEM) clusters, Tregs, and a proliferating T-cell cluster. As expected, synovial tissue is enriched in memory cells with a small contribution of naïve-T-cells, constituting less than 5% of the synovial tissue CD4^pos^ T-cell pool. We integrated the synovial tissue part of this dataset with single-cell spatial transcriptomic data as in Figure 3 to accurately identify T-cell clusters within the tissue. We confidently mapped majority of the T-cell clusters identified by scRNAseq (**Figure S14D-E)**. As expected, active RA synovium is enriched in T-cells compared to remission. We annotated and mapped 6153 T-cells in active RA synovial tissues with a median of 137 cells per FOV compared to 294 T-cells in remission tissue with a median of 12 T-cells per FOV **(Figure 4B).**

**Figure 4:**
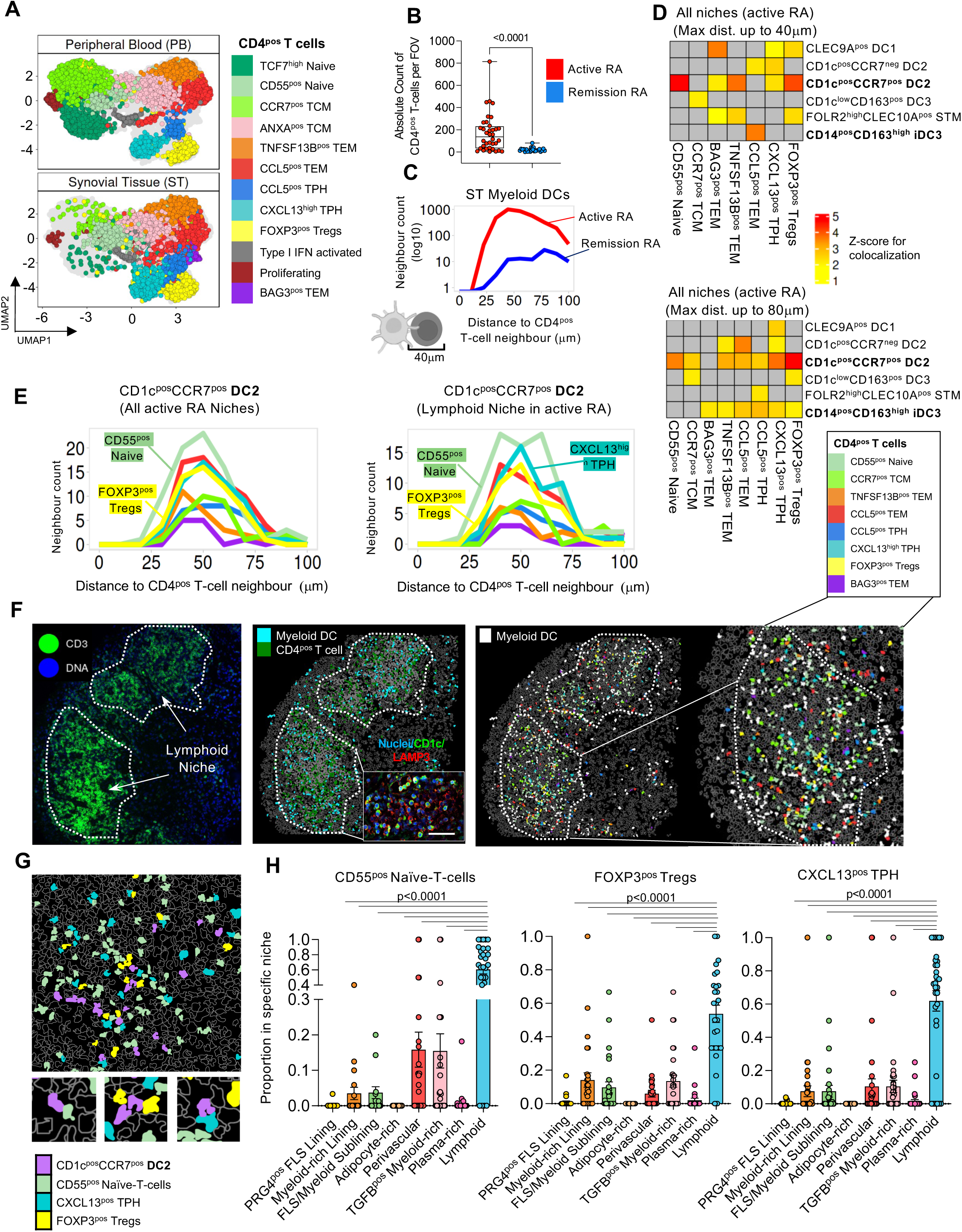
Distinct functions of synovial tissue myeloid dendritic cell clusters. **(A)** Split UMAP visualization of integrated synovial tissue (ST) and blood scRNAseq data from healthy individuals, patients with active RA and RA in remission identifying twelve CD4^pos^ T-cell clusters based on top marker genes (Details in Figure S17). Blood samples: active RA (n=3); synovial tissue samples: healthy (n=3), active RA (n=13), and RA in remission (n=5). **(B)** Absolute number of synovial tissue CD4^pos^ T-cells in all spatially mapped cells per FOV in active RA (n=36 FOVs) and RA in remission (n=16 FOVs) tissues. Data is presented as a box plot with median and interquartile range. **(C)** Neighbourhood analysis (see Methods) shows a higher number of direct (40mm) and proximal (80mm) interactions of myeloid dendritic cells (DC) with CD4^pos^ T-cells in active RA compared to remission RA tissues. Number of FOVs as in (B). **(D)** Neighbourhood analysis of interactions between specific synovial tissue myeloid DC clusters, DC1 and CLEC10A^pos^ STMs and CD4^pos^ T-cells in active RA tissue. The direct (40mm) and proximal (80mm) interactions with CD4^pos^ T-cells that have significant Z-scores are labelled with yellow-red colours, while insignificant ones are marked with grey (n=36 FOVs in active RA). **(E)** Histograms illustrating the distances between CD1c^pos^CCR7^pos^ DC2 cells to different CD4^pos^ T-cell clusters in all niches or specifically in the lymphoid niche. **(F)** Representative images of the lymphoid niche from active RA tissue showing anti-CD3 staining (green), followed by deconvolution of myeloid DC and T-cell interactions, with specific T-cell clusters identified by spatial transcriptomics (CosMx). Representative confocal microscopy images of IF staining for CD1c (green), LAMP3 (red) and nuclei DAPI (blue) showing mReg in the lymphoid niche of synovium. This Image is representative of synovial tissue from active RA (n=6). Scale bar = 20 μm. **(G)** Representative image showing interactions of CD1c^pos^CCR7^pos^DC2 cells with FOXP3^pos^ T-regs, CD55^pos^PDE4B^pos^ naïve-T-cells, and CXCL13^pos^MAF^pos^ TPH cells in synovial tissue based on spatial transcriptomic deconvolution. **(H)** Proportion of T-cell clusters that showed interactions with CD1c^pos^CCR7^pos^ DC2 cells in the lymphoid niche. Data is presented as a bar plot with SEM. One-way ANOVA with Dunnett’s corrections for multiple comparisons. The exact p-values are shown on the graph.

Prior to investigating ST-DC subset interactions with CD4^pos^ T-cell clusters in the tissue, we sought to better understand the functional biology of ST CD4^pos^T-cells that we mapped in active RA synovium. Our unique ST scRNA-seq dataset, capturing T-cell phenotypes across different disease stages added deeper resolution to previously identified ST T-cell clusters in RA^54,65^ (Figure S17). Mining our data with the KEGG cytokine pathway allocated T-cell clusters with functional phenotypes determined by dominant pathogenic cytokine (Figure S17F and Figure S18A). Thus, we identified an additional phenotype of TPH^66^ that exhibits higher levels of TPH activation markers such as PD-1, LAG3, PRDM1, and unique expression of CCL5 (Figure S17F), a potential key chemokine initiating joint flares^2,67^, in addition to the hallmark TPH cytokine CXCL13. The TPH phenotype characterized by CCL5 expression dominates in early naïve-to-treatment RA, while the classical CXCL13-positive TPHs dominate in patients with persistent treatment-resistant synovitis (Figure S17E), suggesting potential evolution from acute CCL5-positive to classical TPH phenotype during disease progression. Both TPHs show an increase in CXCL13 in active RA compared to their counterparts that persist in remission, indicating enhanced activation (Figure S18E-F). Another two large clusters of effector memory CD4^pos^ T-cells in the active RA synovium include: (i) TEM expressing *TNFSF13B*, encoding BAFF (RORA^pos^TNFSF13B^pos^ cluster), which is a key survival factor for B cells, controlling B-cell maturation into plasma cells^68^. (ii) The second TEM cluster is distinguished by high expression of CCL5 (Figure S17F), GZMA (Figure S17C), a trigger of inflammatory mediators from tissue fibroblasts^69^, and IFN-γ, a driver of co-stimulatory molecules expression (CCL5^pos^GZMA^pos^ cluster) (Figure S17F). This cluster is also characterized by high expression of *PD1* and *CD69* (Figure S17F), altogether suggesting a highly activated pro-inflammatory Th1 state.

### Integrating DC subset and T-cell subset biology in RA synovium

We next sought to infer the T-cell stimulatory functions of distinct ST-DC subsets by identifying which T-cell phenotypes (clusters), identified above, they interact with in tissue, using neighbourhood analysis. Briefly, nearest neighbours were identified for each cell, and we determined whether these interactions were significant using a permutation approach; shuffling the position of all cells around the index cell type to reveal the likelihood that those interactions are not expected by chance. Initially, we assessed overall DC/T-cell interactions in the synovium, revealing a higher number of direct interactions (up to distance of 40 μm, the diameter of a cell) between myeloid ST-DCs and CD4^pos^ T-cells in active RA compared to remission RA tissues **(Figure 4C),** thus validating our analytical approach.

In active RA, we explored the interactions between specific ST-DC subsets and specific CD4^pos^ T-cell clusters **(Figure 4D and Figure S16C).** We examined these interactions at two maximal distances: 40 µm to capture the initial DC-driven T-cell phenotypes and 80 µm to capture potential daughter T-cell phenotypes resulting from this interaction. We examined the interactions of CCR7^pos^ DC2 (encompassing mReg and its MIR155^pos^ intermediate stage), CCR7^neg^ DC2, DC3, iDC3, DC1 and CLEC10A^pos^ STMs as non-DC comparator cells with high HLADR expression. This analysis revealed that each one exhibits a distinct pattern of interactions with T-cell clusters, implying varied T-cell stimulatory functions. Overall, mReg and iDC3 showed the highest statistically significant associations with T-cells, potentially indicating the most robust T-cell stimulatory functions.

### mReg DC2 interactions with T-cell subsets in the synovial tissue lymphoid niche

Only mReg/MIR155^pos^ showed statistically significant interactions with naïve-T-cells, and the most significant interactions with Tregs among all DC subsets. Moreover, they exhibited the strongest interactions with BAFF-producing T-cells (TNFSF13B^pos^) and with CXCL13^pos^CCL5^neg^ TPHs, suggesting a unique role in the ectopic germinal centre response in the synovium **(Figure 4D).** Histograms showing the number of interactions of CCR7^pos^ DC2 (mReg/MIR155^pos^) with distinct T-cell clusters at progressing distances, confirms the strongest interactions with naïve-T-cells, Tregs, and classical CXCL13^pos^ TPH in the lymphoid niche that represents ectopic germinal centre **(Figure 4E).** Representative staining for mReg marker LAMP3 and spatial transcriptomic images of the lymphoid niche illustrate high frequency of mReg in this niche and their close interactions with naïve-T-cells, Tregs, and classical CXCL13^pos^ TPH **(Figure 4F-H).**

In summary, CCR7^pos^ (mReg/MIR155^pos^) DCs, but not other DC subsets, uniquely interact with naïve-T-cells in the lymphoid niche. Although these T-cells constitute less than 5% of the total T-cell population in the synovium, they are entirely located in this niche. The highly mature state of mReg/MIR155^pos^ DCs in this geographical location is likely responsible for driving naïve-T-cells toward effector pathogenic function, for example CXCL13^pos^ TPH cells in situ. Additionally, the regulatory gene program of mReg likely underlies its strong interactions with Tregs as part of an immune response-induced negative feedback mechanism.

### ST-iDC3 activate CCL5^pos^ TEM and CCL5^pos^ TPH cells in synovial tissue myeloid-rich hyperplastic lining layer

The myeloid-rich hyperplastic lining layer is a unique feature of active RA synovium and a dominant niche for DC3 and its iDC3 phenotype (**Figure 3I-J).** Although at lower frequency than lymphoid niches, the myeloid-rich lining layer niche contains T-cells including highly activated CCL5^pos^ TPH and CCL5^pos^ TEM of Th1 phenotype (expressing *INF*γ*, PD1, and CD69*) **(Figure 5A-B).** Neighbourhood analysis of spatial transcriptomic data across all niches indicated that iDC3 preferentially interact directly (at a distance of 40 μm) with CCL5^pos^ TEM cells, compared to any other DC cluster **(Figure 4D).** Such interactions extend to CCL5^pos^ TPH cells and, to a lesser extent, to other T-cell clusters when examined over a wider 80 μm area, specifically in the myeloid-rich lining layer **(Figure 4D and Figure 5C-D)**. Independent ligand-receptor analysis of our synovial scRNA-seq datasets (CellPhoneBD) confirmed statistically significant cellular interactions between ST-iDC3 and both CCL5^pos^ TEM and CCL5^pos^ TPH **(Figure 5E)** and identified the potential modes of communication between iDC3 and these T-cell subsets **(Figure S19M).** The interactions of other DC3 phenotypes are limited to central memory T-cells **(Figure 4D),** suggesting that these DC3 may maintain the reservoir of these long-lived cells in synovial tissue, while its iDC3 phenotype might play a prominent role in the activation of pathogenic CCL5^pos^ TEM and CCL5^pos^ TPH clusters.

**Figure 5:**
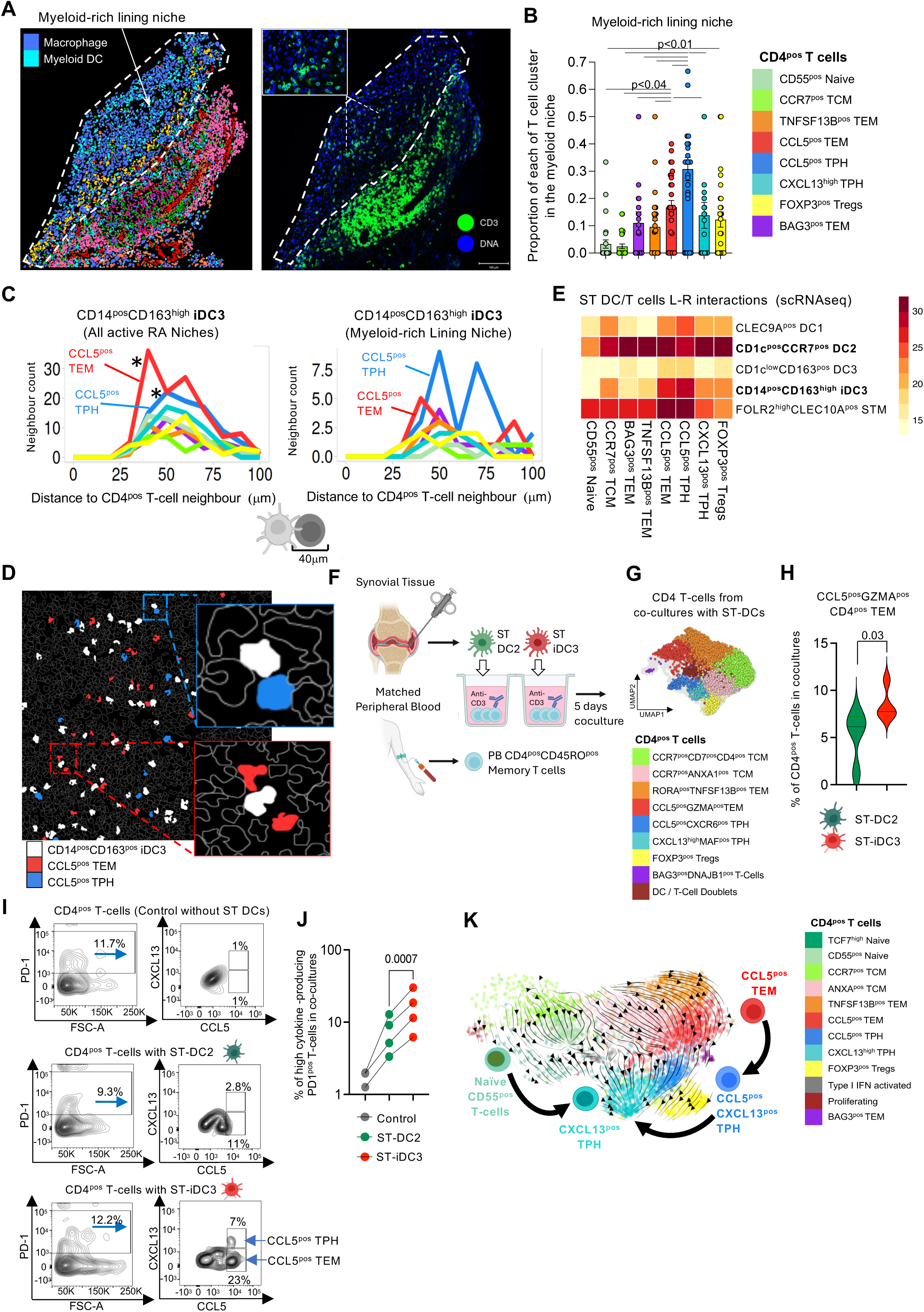
Synovial tissue iDC3 support CCL5^pos^TEM/CCL5^pos^TPH activation in the myeloid-rich hyperplastic lining layer. **(A)** Representative image of the myeloid-rich lining niche from active RA tissue identified by spatial transcriptomics (CosMx) followed by the image of IF staining with anti-CD3 (green) in this niche. **(B)** Distribution of CD4^pos^ T-cell clusters in the myeloid-rich lining layer niche. Each dot represents one FOV (n=36 active RA). Data are presented as the proportion of specific T-cell clusters per this niche (Bar plots with SEM). One-way ANOVA with Dunnett’s corrections for multiple comparisons. The p-values are shown on the graph. **(C)** Histograms illustrating the distances between ST-iDC3 to different CD4^pos^ T-cell clusters in all niches or specifically in the myeloid-rich lining niche (Neighbour analysis of active RA FOVs as in Figure 4) Star represents interactions with significant z-score as depicted in Figure 4D. **(D)** Representative images showing interactions of ST-iDC3 with CCL5^pos^GZMA^pos^ TEM and CCL5^pos^CXCR6^pos^ TPH in synovial tissue based on spatial transcriptomic deconvolution. **(E)** Heatmap showing the absolute number of predicted cellular interactions between T-cell and DC subsets found to be significant (p<0.05) based on scRNAseq data from Figure 1B using CellPhoneBD. **(F)** DC2 and iDC3 were sorted from synovial tissue biopsies of patients with active RA and co-cultured with autologous blood memory CD4^pos^ T-cells for 5 days at 1 to 5 ratio (details in Methods). **(G)** T-cell phenotypes from the co-cultures with ST-DCs were deconvoluted by scRNAseq and integrated with synovial tissue T-cell scRNASeq data to give them an appropriate identity. Details on integration in Figure S19F. **(H)** Violin plot, displaying median and inter-quartile range, depict the percentage of CCL5^pos^GZMA^pos^ TEM in co-cultures with synovial biopsy derived patient-matched ST-DC2 and ST-iDC3. Data from 4 active RA synovial biopsies where both DC subsets were present. T-test, p-value is provided on the graph. The number of successfully sequenced co-culture T-cells ranged from 101 to 1698 (Supplementary material in xls). More details in Figure S19 D-J. **(I)** An example of surface PD-1 and intracellular CCL5 and CXCL13 protein staining in CD4^pos^ memory PB T-cells co-cultured with autologous ST-DC2 or ST-iDC3 (sorted from synovial biopsies according to the CITE-seq gating strategy in Figure S13) in the presence of anti-CD3 antibody (0.25mg/ml) and IL15 (20ng/ml) or in the in the presence of anti-CD3 antibody (0.25mg/ml) and IL15 (20ng/ml) only. **(J)** Dot plot summarizing induction of cytokine production by PD1^pos^ T-cells co-cultured with patient-matched ST-DC2 and DC3/iDC3 (n=4 RA). Paired T-test, p-value is provided on the graph. **(K)** Single-cell trajectory of synovial tissue CD4^pos^ T-cells with RNA Velocity analysis visualized on UMAP. The direction of arrows infers the path of cell trajectory based on spliced versus unspliced RNA counts and suggests differentiation path from (i) CCL5^pos^ TEM to CCL5^pos^ TPH and to CXCL13^pos^ TPH, and (ii) from naïve-T-cells to CXCL13^pos^ TPH.

To test whether in active RA synovium, ST-iDC3 drives the activation of CCL5^pos^ T cells we established micro co-culture systems of synovial biopsy-sorted DCs with autologous memory peripheral blood CD4^pos^ T-cells **(Figures 5F**, **Figure S13 and S19A-L).** We chose memory T-cells because they are the most relevant to the emergence of clinical synovitis. The breach of immune tolerance, which results in the generation of circulating memory pathogenic CD4^pos^ T-cells, occurs in RA patients years before clinical symptoms appear. These cells then localize in the joints due to an unknown yet trigger, contributing to the emergence of synovitis^7^. We included anti-CD3 antibody stimulation to mimic antigen-induced TCR engagement and assessed T-cell activation using scRNAseq to accommodate the low number of T-cells in co-cultures **(Figure 5G).** Briefly, after eliminating STMs and other lineage-positive cells from the synovial biopsy digest, HLADR^pos^CD11c^pos^ cells were gated. Subsequent gating of CLEC10A-positive cells included all ST-DC2 and ST-DC3 and excluded all infiltrating monocytes and other cells not eliminated in lineage-positive gate (as deconvoluted in **Figures S12, Figure S13A-D,** and confirmed in published dataset in **Figures S11A)**. We compared the T-cell-stimulatory potential of patient-matched biopsy-sorted ST-iDC3 (CLEC10A^pos^CD1c^low/neg^) to ST-DC2 (CLEC10A^pos^CD1c^high^). After 5 days of autologous memory T-cells co-culture with ST-iDC3 or ST-DC2, their phenotypes were deconvoluted with an immune-gene panel (399 genes). To accurately annotate the phenotypes of T-cells that emerged from the co-cultures, we integrated their scRNA-seq data with a reference synovial tissue memory CD4^pos^ T-cells scRNAseq dataset **(Figure S19D-F).** A Local Inverse Simpson’s Index (LISI) score of 1.82, where 1 indicates no mixing and 2 indicates optimal mixing of in vitro and in vivo T-cells, suggests a good degree of integration between co-cultured and tissue T-cells **(Figure S19F).** This integration enabled the identification of two TCM clusters, Tregs, and five TEM clusters: TNFSF1B^pos^ TEM, CCL5^pos^ TEM, CCL5^pos^CXCL13^pos^ TPH, CXCL13^pos^ TPH and a minor BAG3^pos^ cluster in the ST-DC/T-cell co-cultures, similar to synovial tissue. Comparison of the frequency of these different T-cell clusters between distinct co-culture conditions revealed a significant increase in the relative frequency of CCL5^pos^ TEM in co-cultures with ST-iDC3 compared to those with patient’s biopsy matched ST-DC2 **(Figure 5H and Figure S19G).** This increase was accompanied by enhanced expression of the activation marker ICOS in CCL5^pos^ TEM cells **(Figure S19H),** while the expression of the proliferation marker *PCNA* and *TCF7* was comparable across different co-culture conditions **(Figure S19I-J).** These findings indicate that ST-iDC3s might be more effective in activating CCL5^pos^ TEM cells compared to ST-DC2, consistent with iDC3/CCL5^pos^ TEM colocalization in situ deconvoluted by spatial transcriptomics.

In synovial biopsies that yielded a sufficiently robust number of ST-DCs, we extended our findings to the protein level using a refined co-culture system. We employed a rigorous gating strategy to sort specific ST-DC subsets based on CITE-seq analysis, which we confirmed with ST-DC index plate sort sequencing **(Figure 1H**, **Figure S12, and Figure S13B-C).** PB Memory CD4^pos^ T-cells were co-cultured with patient-matched ST-DC2 or ST-iDC3 in the presence of anti-CD3 antibody (to mimic antigen stimulation) and IL-15 to provide an additional survival signal for memory T-cells. A control group included T-cells treated with anti-CD3 stimulation and IL-15 only. Intracellular cytokine staining of T-cells was used as a phenotypic readout for TEM (CCL5 & IFNγ) and TPH (CCL5 and CXCL13) activation at the end of the co-cultures. We compared the functions of ST-iDC3 with those of ST-DC2 derived from the same patients’ biopsies and to T-cells incubated with anti-CD3 antibody and IL-15 only. Both ST-DC2 and ST-DC3 significantly enhanced anti-CD3-driven T-cell activation, as evidenced by intracellular cytokine staining. However, ST-iDC3 were notably more effective than matched ST-DC2 in inducing CCL5, CXCL13, and IFNγ production by PD1^pos^ memory T-cells **(Figure 5I-J and Figure S19K).** Additionally, CCL5-producing T-cells co-cultured with ST-iDC3 showed higher expression of the activation marker CD69 compared to those co-cultured with ST-DC2 **(Figure S19L),** reminiscent of the CCL5^pos^ TEM/TPH phenotypes found in tissue (**Figure 17E)**. Taken together, these findings confirmed our tissue spatial transcriptomics and initial co-cultures, indicating that ST-iDC3 can activate memory CCL5^pos^ TEM and CCL5^pos^ TPH.

To dissect the relationship between these two CCL5^pos^ T-cell clusters driven by ST iDC3, we conducted RNA velocity-based cell trajectory analysis on all ST CD4^pos^ T-cell clusters **(Figure 5K).** This data infer that CCL5^pos^CXCL13^pos^TPH cells can originate from CCL5^pos^TEM cells, providing an explanation for the colocalization of these two CCL5-positive T-cell clusters with iDC3 in the myeloid-rich lining niche and their synchronised increase in co-cultures with ST-iDC3. This analysis also inferred the subsequent progression of CCL5^pos^ CXCL13^pos^ TPH to classical CXCL13^pos^ TPH, reflecting the progression in memory TPH phenotypes observed during the RA trajectory from early RA to chronic, difficult-to-treat disease **(Figure S17E).** Another notable developmental trajectory of CXCL13^pos^ TPH in the synovium is inferred from naïve-T-cells. This accounts for the co-localization of naïve-T and CXCL13^pos^ TPH cells with mReg DC2 in the lymphoid niche.

Together these data indicate that ST-iDC3 are responsible for the activation of CCL5^pos^ TEM and their potential differentiation into CCL5^pos^ TPH in the hyperplastic myeloid-rich lining layer.

### A specific inflammatory signature of blood CD14^pos^ DC3 (iDC3) predicts disease flare in remission RA

To assess the contributions of distinct ST-DC subsets to initiation of joint pathology, we examined the phenotypes of their blood predecessors in a human model of disease flare following treatment withdrawal (the BioRRA cohort^70,71^) **(Figure 6A).** In RA patients (n=12) who achieved sustained clinical and ultrasound remission with cDMARDs, we investigated the frequency and transcriptomic signature of PB DC1, DC2, DC3 and its distinct clusters defined by CD163 and CD14 expression, including iDC3 phenotype (CD163^high^CD14^pos^)^20,21^, as well as monocytes as comparators at baseline (the time of treatment withdrawal) and at a second follow-up time point. This second time point was either at the occurrence of disease flare during the six-month patient follow-up period (flare endpoint) or at the end of six-months if the patient-maintained drug-free remission (drug-free remission endpoint). The blood myeloid cell subsets and their phenotypes were deconvoluted with a 399 immune-gene panel in scRNAseq **(Figure 6B-C and Figure S20A-B).** Among the 12 patients investigated, 8 flared while 4 remained in drug-free remission after treatment withdrawal. The outcomes of flare or remission informed a categorical analysis of myeloid dendritic cell subsets at baseline and follow-up time points. Consistent with our previous studies on macrophage-driven mechanisms of flare^43^, we observed at baseline an increased frequency of activated CD14^pos^CXCL8^pos^ monocytes that give rise to S100A12^pos^ STMs in the synovium of patients in remission who subsequently flare **(Figure S20C).** We did not observe statistically significant differences in the frequencies of different PB DC subsets at baseline or at the time of disease flare **(Figure S20C).** However, comparison of their transcriptomic profiles at baseline revealed a pro-inflammatory gene signature (upregulation of expression of 33 genes) that distinguished patients who subsequently flared from those who would have remained in remission The upregulated expression of 24 of these pre-flare genes persisted in those patients predicted to flare and remained significantly upregulated at onset of flare **(Figure 6D)**.This gene signature was completely absent in patients who achieved drug-free remission, suggesting the stability of this gene panel in predicting flare. This gene panel include increased expression of integrins that facilitate migration into tissue (e.g., *ITGAM, ITGB2*), pattern recognition receptors (e.g., *TLR2 and CLEC4E*), and alarmins (e.g., *S100A12, S100A9 and LGALS3*), suggesting an increased activation **(Figure 6E).** This pre-flare gene module was mostly confined to the PB iDC3 population **(Figure 6F)** and the ST-iDC3 clusters that mature from them in tissue **(Figure 6G)**. This suggests that the activation of blood predecessors of ST-iDC3 clusters precedes pathology in the joint and supports their role in the initiation of synovitis e.g., by driving the activation of CCL5-producing TEM/TPH in tissues.

**Figure 6.**
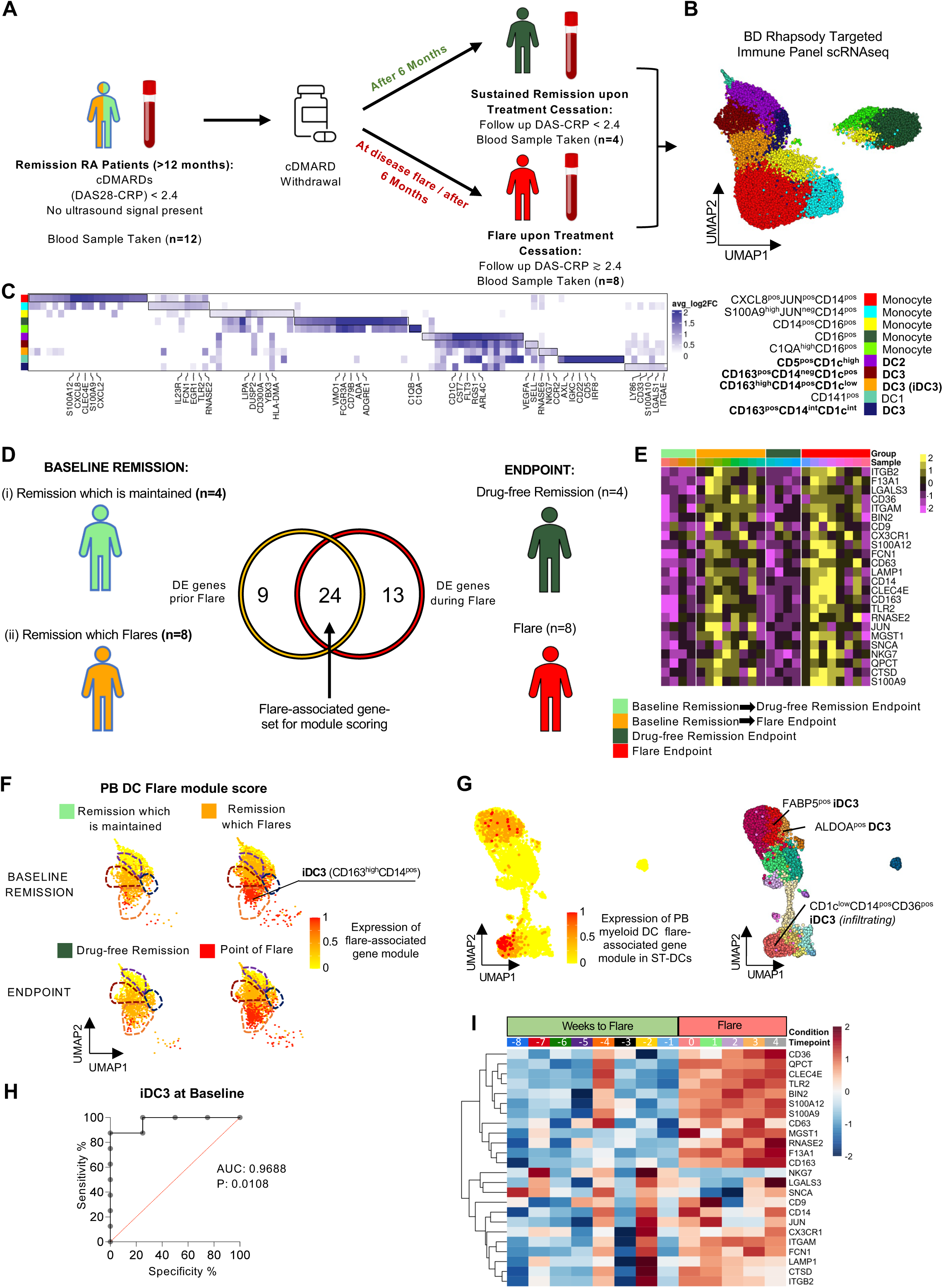
Transcriptomic profiles of peripheral blood CD14^pos^ cluster of DC3 (iDC3) of RA patients in remission are predictive of flare. **(A)** Schematic detailing the layout of the BioRRA Study. cDMARDs, conventional disease modifying anti-rheumatic drugs. Details in Table S8. **(B)** UMAP visualization of ten peripheral blood myeloid cell clusters identified in scRNAseq analysis using an immune panel (399 genes), supported by integration with the whole transcriptome peripheral blood myeloid cell scRNAseq dataset (details in Figure S2). **(C)** Heatmap illustrating the average logFC for genes identified as cluster markers (expressed in >40% of cells/cluster, with log-fold change >0.25, and *p*<0.05 based on MAST with Bonferroni correction for multiple comparison). Differentially expressed marker genes annotated based on greatest average logFC per cluster. **(D)** Schematic detailing flare-associated gene selection in myeloid dendritic cells (DC2, DC3 and its iDC3 phenotype) for preparation of DC flare-associated gene module score. Venn Diagram illustrates differentially expressed genes shared between baseline and end-point conditions (with log-fold change >0.25, *p*<0.05 based on Wilcoxon Rank Sum test with Bonferroni correction for multiple comparison). **(E)** Heatmap visualising scaled expression of 24-genes from the DC flare-associated gene module score by sample and condition timepoints. **(F)** UMAP visualization of the expression of PB myeloid DC flare-associated gene module score by distinct PB DC subsets indicates the highest expression in PB iDC3. **(G)** UMAP visualization of the expression of PB myeloid DC flare-associated gene module by synovial tissue myeloid DCs indicates the highest expression in ST-iDC3 clusters. **(H)** Receiver-operating characteristic curves (ROC-AUC) illustrate the performance of the average flare-associated module score at predicting flare upon treatment withdrawal at baseline in PB iDC3. Statistical information annotated beside plots. **(I)** Heatmap visualising mean scaled batch-corrected expression of genes from BioRRA Flare-associated Gene-Set across weeks relative to flare and during flare in the PRIME-cell data set of disease flare (*Orange et al, NEJM, 2020*). Data is generated from an index patient over 364 time points, both preceding and during eight flares, spanning a period of 4 years.

In addition, AUC-ROC analysis of the pre-flare iDC3 gene module exhibited high sensitivity and specificity in stratifying these two remission outcome groups at baseline **(Figure 6H),** suggesting the potential of the iDC3 pre-flare signature as a biomarker of flare.

To verify this in an independent dataset, we investigated the expression of genes from the DC flare-associated gene module across the weeks leading to flare and during flare in the longitudinal disease flare PRIMEcell study^72^. This dataset encompasses over 364 time points, both preceding and during eight flares, spanning a period of four years. We observed a fluctuating increase in the expression of the genes in our module starting four weeks before flare and persisting throughout the flare **(Figure 6I)**, confirming the biomarker potential of the iDC3 signature in predicting disease flares.

## Discussion

This study delineates the heterogeneity and functions of myeloid dendritic cells (DCs) in the human synovium in health, compared with active inflammation, and remission in rheumatoid arthritis (RA). We identified transcriptionally distinct synovial tissue myeloid DC phenotypic clusters of DC2, DC3 and its iDC3 phenotype, that differ in frequency between healthy individuals and patients with RA with active disease or in remission. Synovial DC subsets exhibit distinct tissue-niche distribution and are associated with different T-cell stimulatory functions, altogether providing a better understanding of joint tissue immunobiology. The deconvolution of the DC landscape in synovium offers insights into mechanisms that maintain tissue tolerance in healthy synovial tissue, drive autoimmunity in RA, or restrain autoimmunity or mediate flares in sustained remission.

Healthy synovial tissue possesses a unique tissue-resident AXL^pos^ DC2 cluster, which is predicted to differentiate from infiltrating KLF4^pos^ DC2 and resides in the superficial sublining layer, just below the protective lining-layer of TREM2^pos^ STMs. They exhibit a tolerogenic phenotype, and as such are likely crucial for maintaining local immune tolerance. We propose that, together with inflammation-resolving TREM2^pos^ STMs^43,44^, this comprises a myeloid innate system that is responsible for maintenance of synovial tissue homeostasis. The relative frequency of AXL^pos^ DC2 cells within the ST myeloid pool decreases in active RA. Instead, infiltrating KLF4^pos^ DC2 cells develop into mReg with a strong immunogenic and migratory program (e.g., high expression of CD40 and CCR7, and production of IL-6 and IL-23) in a process driven by MIR155. A recent study ^73^ indicated that other features of mReg, such as IL-12 production, are also under MIR155 control, supporting a key role of MIR155 in the transcriptomic program of mReg. We found mReg/MIR155-positive DC2 signature in lymph draining from the tissue, supporting their CCR7-driven migratory potential to lymph nodes. However, mReg DC2, and their highly activated CCR7^pos^MIR155^pos^ intermediate stage, are also abundant in ectopic lymphoid structures (niches) in synovium of active RA, suggesting their local retention in inflamed tissue. In this niche they uniquely interact with a small population of naïve-T-cells homing specifically to this niche, and CXCL13^pos^ TPH. Recently, similar close physical interactions of mReg with CXCL13^pos^ TPH were mapped in discrete cellular niches of liver and lung tumours of patients responding to immune therapies^66^ ^74^, inferring that mReg/MIR155^pos^ DC2 could drive CXCL13^pos^ TPH differentiation from naïve-T-cells locally in tissue. Thus, mReg/MIR155^pos^ DC2 might orchestrate local pathogenic antigen-specific effector immune responses of naïve-T-cells, fuelling the germinal centre response in active RA synovial tissue.

Ectopic germinal centre structures resolve in synovium during sustained remission, during which mReg maintain an immunogenic program but also acquire an additional immunoregulatory program. This includes higher expression of immune checkpoints PD-L1 and PD-L2, suggesting an active potential to limit autoimmunity. The functional interactions between mReg cells and regulatory T-cells in healthy and remission tissue remain to be investigated. In active RA, mReg/MIR155^pos^ DC2 cells, in addition to interacting with naïve-T-cells and CXCL13^pos^ TPH cells, showed the highest frequency of localization with regulatory T-cells among all synovial tissue dendritic cell subsets/clusters. This supports their potentially unique role in balancing the priming and limiting of the immune response.

Inflamed RA synovial tissue contains a unique myeloid-rich hyperplastic lining layer that develops in place of the TREM2^pos^ protective barrier present in health. It is populated by inflammatory subsets of MERTK-negative macrophages^5,43^. We show that this niche also hosts a highly activated tissue FABP5^pos^ iDC3 cluster that likely matures from an infiltrating CD14^pos^ phenotype of DC3 (iDC3) and is characterized by NFκB signatures and unique T-cell stimulatory functions. In this niche, iDC3 specifically interact with PD1^pos^CCL5^pos^IFNγ^pos^GZMA^pos^TEM and a highly activated memory PD1^pos^CXCL13^pos^TPH cluster expressing CCL5 (CCL5^pos^CXCL13^pos^). Co-culture of synovial tissue biopsy-sorted DCs with autologous memory T-cells confirmed that ST-iDC3 activate CCL5^pos^ TEM cells and induce their differentiation into CCL5^pos^CXCL13^pos^ TPH cells. The latter cluster of TPH cells characterizes early synovitis and is predicted to differentiate into classical CXCL13^pos^ cells, which dominate in later disease stages. This suggests that iDC3-memory CCL5^pos^ TPH might drive early synovitis and mediate progression from a myeloid to a lymphoid pathotype^7,75^ with ectopic germinal centres homing autoreactive naïve-T-cells for in situ development of TPH. Studies with longitudinal synovial tissue biopsies, spanning from pre-clinical RA to onset of arthritis are needed to validate this hypothesis.

Finally, our human RA disease flare model (BioRRA) demonstrates that a pro-inflammatory gene module characterising tissue iDC3 is present in a PB iDC3 cluster before disease flare onset. This finding, along with a recent report showing an increase in CCL5^pos^ TEM (GZMA^pos^ CD4) at the onset of flare^71^, supports the potential role of the iDC3 - CCL5^pos^ TEM/TPH axis as a critical initiating step in localising immune responses in joints.

Recently, an increase in the infiltration of DC3 has been reported in many tissue pathologies^20,27,21,24,30^. Expansion of the CD14^pos^ DC3 (iDC3) subset was reported in the blood and kidneys of SLE patients^20,27^, where TPH activation and autoantibody production are hallmark pathogenic mechanisms, supporting the role of iDC3 in TPH response. Bourdely et al^21^ proposed that circulating DC3s are immediate precursors of these cells. Our matched blood-tissue DC trajectory analysis is consistent with this finding and indicates that the CD14^pos^ phenotype of PB DC3s is the most likely predecessor of tissue DC3 clusters.

In sustained disease remission, RA synovial tissue exhibits resolution of the hyperplastic myeloid-rich lining and restoration of a protective TREM2^pos^ STM lining-layer^5^. This is accompanied by a decrease in the frequency of iDC3 and restoration of DC2 in the superficial lining layer, resembling the structure of healthy synovium. However, these cells do not fully acquire the AXL^pos^ DC2 phenotypic program characteristic of healthy tissue. Instead, they stall at a predicted intermediate stage, KLF4^pos^ATF3^pos^ DC2, showing the restoration of only some, but not all, of the tolerogenic potential characteristic of healthy tissue DC2. This suggests a latent potential for disease flare in the joints of patients in remission and offers an intriguing therapeutic opportunity to strengthen remission in RA patients once achieved.

Plausible mechanisms preventing the acquisition of the AXL^pos^ tolerogenic DC2 phenotype in remission tissue, as compared to healthy tissue, may include epigenetic changes in their bone marrow precursors and/or the lack of recovery of an appropriate maturation signal in the remission tissue niche, both as result of prior chronic inflammation. The former concept is supported by our observation that changes in the transcriptomic profile of DCs in circulation precede joint disease flare in patients in remission. Causal determinants could include an increase in the epigenetic regulator miR34a that targets the immune-checkpoint AXL for degradation, as we have shown previously^41^. It remains to be investigated how interactions of tissue-infiltrating DC subsets with healthy, inflamed, and resolved stromal synovial tissue niches contribute to the maturation of different phenotypic clusters.

Our strategy of examining tissue DCs from homeostasis through active disease to disease resolution, including matched blood counterparts, enabled the discovery of tissue niche and condition-specific phenotypes (clusters) with diverse functions acquired by infiltrating myeloid DC2 and DC3. It has been previously shown that blood DC1 and DC2 can acquire, in the tissue, the mReg molecular state characterized by the induction of an immunogenic, regulatory, and LN migratory gene program^33,34^. Additionally, we provided here evidence of additional molecular states (clusters) beyond the mReg phenotype developing from DC2 depending on the tissue environment. For example, the tolerogenic AXL^pos^ phenotype acquired by DC2 in a healthy tissue state and the TGFβ-bearing signature KLF4^pos^ ATF3^pos^ acquired by DC2 in a post-immune state.

In summary, our data suggest that therapeutic strategies to block the pathogenic functions of the ST-iDC3s and their blood predecessors and reinstating the tolerogenic functions of the ST AXL^pos^ DC2 cluster might be the necessary step-change for resolution of synovitis and transition from remission into self-maintained tolerance. Additionally, the specific inflammatory signature of PB predecessors of ST-iDC3, which predict the disease flare, could inform the management of RA during the fragile clinical (DAS28-defined) remission.

### Limitations of the study

The primary limitation of this study is the lack of functional insight into the potential of ST-DC subsets to activate RA autoantigen-specific naïve-T-cells in an autoantigen-specific manner. Currently, this type of study is extremely challenging due to the very low frequency of T-cells specific for various citrullinated peptides in the peripheral blood of patients with RA (less than 1%)^1^. CRISPR-Cas9-mediated engineering of patients’ T-cells to express citrullinated peptide-specific TCRs could provide a necessary experimental platform to elucidate this in the future.

## Acknowledgments

This work was supported by the Research into Inflammatory Arthritis Centre - *Versus Arthritis* UK (grant no. 22072) and *Versus Arthritis* UK project grants (no.22253, 22273 and 23229) to M.K-S, by Linea D1 (Università Cattolica del Sacro Cuore, no. R4124500654), and Ricerca Finalizzata Ministero della Salute (no. GR-2018-12366992) to S.A, and by a FOREUM fellowship to A.P. The BioRRA study was funded by research grants from the Wellcome Trust (102595/Z/13/A to K.F.B) and the NIHR Newcastle Biomedical Research Centre (BH136167/PD0045 to K.F.B). K.F.B is supported by a Newcastle Health Innovation Partners Senior Clinical Fellowship, an NIHR Advanced Fellowship (NIHR303620) and the Newcastle Hospitals Charity. JDI is a NIHR Senior Investigator. The views expressed are those of the authors and not necessarily those of the NIHR, the NHS, or the UK Department of Health and Social Care. We would like to thank Claire Williams and colleagues at Nanostring for their support with generation and analysis of spatial transcriptomics data. We acknowledge the support of Scott Arkison in maintaining servers for computational analysis and the staff in the imaging (Dr Leandro Lemgruber Soares) and in the flow-facility labs (Mrs Diane Vaughan and Mrs Alana Hamilton) in the School of Infection & Immunity (University of Glasgow). We would like to thank Dr Dana Orange (the Rockefeller University, New York, USA) for an access to the PRIMEcell data set and Prof Allan Mowat and Dr Megan MacLeod (both at University of Glasgow) for critical reading of this manuscript. We acknowledge the Immunology Research Core Facility of the Gemelli Science and Technology Park (GSTeP) of the Fondazione Policlinico Universitario A. Gemelli IRCCS for all synovial tissue biopsies handling. We are extremely grateful to all the RA patients and the healthy volunteers who contributed to this study and to the nurse team of the SYNGem Biopsy Unit of the Fondazione Policlinico Universitario A. Gemelli IRCCS.

## Author Contributions

**L.M**. contributed to the study concept and manuscript writing, supervised all bioinformatic analysis and performed (i) analysis and interpretation of scRNAseq/CITEseq of ST myeloid cells and DCs, (ii) integration with the blood dataset, (iii) establishment of ST-DC clusters, and (iii) analysis of the mouse DC RNAseq dataset. She established the pipeline for single-cell spatial transcriptomic analysis, integrated it with scRNAseq, identified tissue niches, and identified DC/T interactions. She also performed and interpreted cell trajectory analysis, assisted J.F. with DC/T cell scRNAseq co-culture and BioRRA cohort data analysis, and aided D.S. in deconvolution of surface markers of ST-DCs by CITEseq. **A.E**. contributed to the study concept and manuscript writing, (i) optimized the new CITEseq-guided ST-DC panel and contributed to ST-DC index plate sort, (ii) validated DC scRNAseq data using flow cytometry, performed (iii) sorting of ST-DC subsets for co-cultures and myeloid cells of the BioRRA study for biomarker scRNAseq experiments. She also performed (iv) direct co-culture of ST-DC clusters with autologous memory CD4^pos^ T-cells and their flow cytometry analysis, (v) transfections of PB and ST CD1c^pos^ DCs with miR-155 mimic and validation of their cytokine expression, (vi) all IF staining in this study (vii) and contributed to the CITEseq of ST DCs. **D.S.** performed scRNAseq and CITEseq of ST-DCs, established and interpreted ST-DC index plate sort, analysed CITEseq data, optimized and tested the CITEseq-guided ST-DC panel, integrated ST-DCs with the MNPVerse dataset, contributed to the sorting of myeloid cells of the BioRRA study for biomarker scRNAseq experiments, helped J.F. in integration of BioRRA with PRIMEcell dataset, performed with A.E. direct co-culture of ST-DC clusters with autologous memory CD4^pos^ T-cells and their flow cytometry analysis, and contributed to study concept and manuscript writing. **J.F.** contributed to study concept, manuscript writing, and editing, performed analysis of scRNAseq of ST T-cells, T-cells from DC/T-cell co-culture experiments, and integration and interpretation of these two datasets, assisted L.M. in analysis of ST DC scRNAseq data, was involved in optimisation and acquisition of flow cytometry co-culture system run by A.E. and D.S., assisted L.M. with integration and analysis of scRNAseq T-Cells and DCs with single-cell spatial transcriptomic data, analysed BioRRA dataset, and tested the biomarker potential in PRIMEcell dataset. **A.P.** performed DC experiments and analysis on miR-155 deficient mice**. T.S.** performed scRNAseq of blood myeloid DCs and assisted in establishing ST-DC/T-cells co-cultures. **L.A.C** assisted in establishing ST-DC/T-cells co-cultures and contributed to enrolment of patients to study cohorts. **O.M.H.** performed ligand-receptor interaction analysis of ST-DC and CD4 T-cells and assisted with cell trajectory analysis. **S.A. and D.B.** performed all synovial biopsies included in the study**. S.A., B.T., C.D.M., and D.C.** handled synovial tissue processing and linked clinical information with experimental datasets. **S.A., E.G., M.R.G., S.P., D.B., M.A.D., and L.P.** enrolled the study cohorts. **C.D.M. and B.T.** performed serological assessment for autoantibody positivity for the entire study cohort. **R.B.** handled all synovial tissue processing for histology, and **M.G.** performed semiquantitative assessment of synovitis scores for all synovial biopsies. **R.M.** performed cell segmentation of spatial transcriptomic data and with **K.W. and I.K.** provided pipelines and expertise in analysing spatial transcriptomic data**. C.M.** assisted with data interpretation and manuscript writing. **N.L.M.** provided healthy synovial tissues. **K.F.B.** and **J.D.I.** provided the BioRRA patient cohort and clinical information **A.F., D.M. and J.P.X.M** collected cells from BioRRA patient cohort. **S.M.** provided expertise and RNAseq data on DCs sorted from mouse gut, lymph, and mesenteric lymph nodes. **E.G., K.A., and I.B.M.** assisted with data interpretation. **T.D.O.** assisted with computational analysis. **S.A. and M.K.S.** initiated the study concept, designed and supervised the overall work, and wrote the manuscript.

## Methods

### Rheumatoid Arthritis Patients and Healthy Donors

#### Patient recruitment and management

To study synovial tissue DCs and their interactions with T-cells, 86 patients fulfilling the EULAR classification criteria revised criteria for RA^76^ were enrolled and underwent ultrasound-guided synovial tissue biopsy of the knee or the wrist as a part of ongoing recruitment to the SYNGem cohort^43,77,78^ (standard-of-care protocol, Division of Rheumatology, Fondazione Policlinico Universitario A. Gemelli IRCCS – Università Cattolica del Sacro Cuore, Rome). RA patients were stratified into treatment-naïve, treatment-resistant RA (inadequate responders to conventional or biological Disease Modifying Anti-Rheumatic Drugs, c/b DMARDs), difficult-to-treat RA (inadequate responders to two or more bDMARDs)^79^ and patients in sustained (>9 months) clinical and ultrasound steroid-free remission maintained by stable conventional and/or biologic-DMARDs^78^. Healthy donors (n=12) attending arthroscopy for meniscal tear or cruciate ligament damage, with normal synovium (macroscopically and by MRI) were included as a control group (University of Glasgow). The criteria for RA patients in sustained clinical remission included (i) DAS44 (disease activity score in 44 joints)<1.6 or DAS28<2.6 at 3 sequential assessments (each 3 months apart), and (ii) ultrasound remission (Power Doppler negativity by US assessment at 3 sequential evaluations, each 3 months apart)^43^ ^78,80^. The clinical and laboratory evaluation of each RA patient enrolled included DAS based on the number of tender and swollen joints of 44 or 26 examined, plus the erythrocyte sedimentation rate (ESR) and plasma C-reactive protein (CRP). Peripheral blood samples were tested for IgA-RF and IgM-RF (Orgentec Diagnostika, Bouty-UK), and ACPA (Menarini Diagnostics-Italy) using commercial Enzyme-Linked Immunosorbent Assay (ELISA) and ChemiLuminescence Immunoassay (CLIA) respectively. Peripheral blood from additional 12 healthy donors matched by age and sex to the RA patients was collected. Demographic, clinical, and immunological features of the SYNGem RA cohort and healthy donors enrolled in each experiment included in the manuscript are shown in Tables S1-S7.

To study the peripheral blood predecessors of synovial tissue dendritic cell clusters we used the BioRRA RA disease remission cohort^70,71^ (Table S8). Patients fulfilling the American College of Rheumatology (ACR) / European Alliance of Associations for Rheumatology (EULAR) 2010 or ACR 1987 classification criteria for RA^76,81^ and in remission (DAS28-CRP< 2.4 and no Power Doppler signal on a 7-joint ultrasound examination) on cDMARDs (methotrexate, sulfasalazine and/or hydroxychloroquine therapy) stopped all DMARD therapy without tapering. Other medications, including non-steroidal anti-inflammatory drugs were continued if required. Study reviews were scheduled at months 1, 3 and 6, with additional study visits if requested by the patient. During each review, the maintenance of remission or the emergence of disease flare (defined as DAS28-CRP ≥ 2.4) was recorded.

The study protocols were approved by the Ethics Committee of the Università Cattolica del Sacro Cuore (28485/18, 30973/19 and 14996/20 for the SYNGem cohort), by the Northeast Tyne & Wear South Research Ethics Committee (National Health Service Health Research Authority, reference 14/NE/1042 for the BioRRA cohort) and by the West of Scotland Research Ethics Committee (19/WS/0111 for healthy donors). Use of BioRRA samples was authorised by the Newcastle Biobank Committee under the approval of the Northeast – Newcastle & North Tyneside 1 Research Ethics Committee (17/NE/0361). All subjects provided signed informed consent. The exact number of patients constituting different data set is provided in Tables S1-8 and in legend to each Figure.

### Semiquantitative histological assessment of synovitis degree

Synovial tissue specimens were fixed in 10% neutral-buffered formalin and embedded in paraffin. Briefly, paraffin-embedded synovial tissue specimens were sectioned at 3μm. Sections were stained for Haematoxylin and Eosin as follows: sections were deparaffinized in xylene and rehydrated in a series of graded ethanol, stained in haematoxylin and counterstained in Eosin/Phloxine. Finally, sections were dehydrated, cleared in xylene and mounted with Bio Mount (Bio-Optica). Slides were examined using a light microscope (Leica DM 2000). The severity of synovitis was graded according to the three synovial membrane features (synovial lining cell layer, stromal cell density and inflammatory infiltrate), each ranked on a scale from none (0), slight (1), moderate (2), and strong (3). The values of the parameters were summed and interpreted as follows: 0–1 no synovitis, 2–4 low-grade synovitis, and 5–9 high-grade synovitis^77^.

### Sample preparation for single cell RNA-seq and CITE-seq of peripheral blood and synovial tissue

To investigate synovial tissue myeloid DC and T-cell heterogeneity in RA, we sequenced the transcriptomic profile of 143,851 cells from peripheral blood mononuclear cells (PBMC) (n=8, including 3 matched with synovial tissues) and synovial tissue (ST) biopsies (n=34). The latter includes synovial tissue from healthy donors (n=7), patients with active RA (n=18) and patients in sustained remission (n=9). Synovial biopsies were taken using 14G Precisa Needle (HS Hospital Service, Italy) in ultrasound-guided protocol^43,82^, and digested as described previously^43,54,65^. All live cells or CD45^pos^ immune cells, were FACS sorted, after exclusion of dead cells with Fixable Viability Dye eFluor® 780 (eBiosciences). Maximum 20,000 immune cells from blood or tissue cells were sorted into Protein LoBind 1.5 ml Eppendorf tube containing 300 μl of RPMI media with 10% of FSC. Cells were loaded onto a Chromium Controller (10X Genomics) for single-cell partitioning, followed by library preparation using Single-Cell 3’ Reagent Kits v3.1. For 3 healthy ST and 4 ST from active RA **(Figure S1),** Totalseq Hashtag were used (Biolegend #394601, #394603, #394605, #394607, #394609) to combine 2 samples per run, and TotalSeq™-A Human Universal Cocktail (V1.0) was used to collect protein expression (CITE-seq) data together with transcriptome data. Single-cell libraries were sequenced on the Illumina HiSeq 4000 system to a minimum depth of 50k reads/cell.

### ST-DC index SORT-seq, flow cytometry evaluation of ST-DC subsets’ frequency and sorting for the co-cultures with autologous T-cells

CITE-seq and our previous data on synovial tissue myeloid cells^43^ provided surface markers for the gating strategy to evaluate ST-DC subset frequency and sort them for co-cultures with T-cells, which was validated in ST-DC index SORT-seq **(Figure S12 and S13).** ST cell suspensions were stained with Fixable Viability Dye eFluor® 780 (eBiosciences) and a CITE-seq-guided antibody panel (Biolegend) to identify distinct DC subsets. The gating strategy was based on fluorescence minus one (FMO) and/or unstained controls. Catalogue numbers and fluorochromes of antibodies are provided in **Table S9-10.** Briefly, live cells and then CD45^pos^ cells were gated. In the next step, lineage-positive cells expressing CD3 (T-cells), CD19/20 (B-cells), CD15 (neutrophils), CD117 (mast cells), and CD56 (NK/NKT cells) were excluded. Subsequently, cells expressing high levels of HLA-DR or HLA-DR and CD11c were gated. This was followed by gating cells negative for the synovial tissue macrophage markers, FOLR2&MerTK. Cells expressing FOLR2 and CLEC10A are CLEC10A^pos^ STM. Gating on FOLR2&MERTK-negative, CLEC10A and CD39&32-positive cells captures all clusters of DC2, DC3 and its iDC3 phenotype, and exclude CD14^pos^CD16^pos^ tissue monocytes and TNF^pos^ICAM1^pos^ STMs because they lack or show low expression of CLEC10A and CD32&39. Both DC3 and its iDC3 phenotypes as well as SPP1^pos^ STM clusters express CD9 but SPP1^pos^ STM cluster can be excluded from DCs by the lack of CLEC10A. CLEC10A-positive CD32&39-positive DCs were gated into different subtypes based on the combination of CD1c and CD14 expression: DC2 by high CD1c, iDC3 by low CD1c and high CD14 expression, DC3 by low CD1 and lack of CD14 **(Figure S12 and Figure S13).**

To validate this sorting strategy, ST DC2, DC3 and its DC3 phenotypes, as well as SPP1^pos^ and CLEC10A^pos^ STMs to serve as negative controls, were sorted using Sony MA900 directly into two 384-well plates to perform ST-DC index SORT-seq (a modified version of CEL-Seq2) (Figure S13B-C). Library generation and sequencing were provided by Single Cell Discoveries (Utrecht, Netherlands). Read alignment and generation of count matrices from raw data were performed using STAR (v 2.7.11a) pipeline against the Human Genome (GRCh38-3.0.0) and performed UMI counting. The ST-DC index SORT-seq confirmed the accuracy of the ST-DC subset gating strategy (Figure S13C). This gating strategy was used for the co-cultures of ST-DC subsets with PB memory CD4^pos^ T-cells, in which T-cell phenotypes were evaluated by intracellular cytokine production, and expression of surface costimulatory molecules *(Co-culture-2 below).* The same gating strategy was also used to evaluate the frequency of ST-DC subsets in synovial biopsies from patients with active RA and RA in disease remission (Figure 1H-I). For the initial ST-DC subset/T-cell co-culture experiment *(Co-culture-1),* we used a smaller panel that included markers for the exclusion of lineage-positive cells as above (dump-channel, Table S9). The STMs were excluded by high surface expression of CD14 that similarly to studies by Cytlak et al^23^ showed at least 1 log higher expression of CD14 compared to ST-iDC3. The accuracy of this initial panel was further confirmed through back validation using ST-DC index SORT-seq (Figure S13D), which shows that inclusion of low and intermediate CD14 expression capture majority of DC3/iDC3 and excludes majority of STMs. To ensure the specific capture of all ST-DCs, CLEC10A positive cells were gated and ST-DC2 were identified by high CD1c expression while DC3/iDC3 were identified by high CLEC10a and low/neg CD1c expression (Figure S19A-B).

#### ST-DC/T-cells Co-cultures. Co-culture-1: scRNAseq of T-cells

Autologous memory CD4^pos^ T-cells were FACS-sorted from PBMCs based on their co-expression of CD3, CD4 and CD45RO. Details of the antibodies used are provided in Table S9. We used CD3 activating antibody to mimic TCR engagement (BioLegend, #300438 clone UCHT1). ST-DC subsets and T-cells were sorted into FACS tubes containing complete RPMI1640 medium (10% FCS, penicillin/streptomycin 100U/mL, and 2mM Glutamax). Co-culture was set up when synovial biopsy yielded enough cells at least in one ST-DC subset (minimal 200 cells). Cells were co-cultured at a 1:5 ratio for each DC subset (200-1000 cells) plus memory T-cell (1000-4000 cells) in 200μl of X-VIVO 15 Serum-free Cell Medium (BE02-060F, Lonza) supplemented with 2% human serum (H3667, Sigma) or in complete RPMI1640 in a 96-well round-bottom cell-culture plate (3799 – SLS, Corning). After 5 days in culture, the changes in T-cell phenotype were investigated using scRNAseq (BD Rhapsody Immune Response Panel, described in the section below). Data from all co-cultures were used to build a Seurat object of T-cells. Subsequently, only those that yielded results from both matched ST-DC subsets (DC2 and DC3/iDC3) were used in the statistical comparison. *Co-culture-2: Intracellular cytokine staining of T-cells.* Guided by the data from the first co-cultures, we modified culture conditions to better support T-cell responses. Briefly, autologous memory CD4^pos^ T-cells were enriched via negative selection from PB of RA patients with active disease using the EasySep™ Human Memory CD4+ T Cell Enrichment Kit (STEMCELL, #19157), which minimises T-cell stress as compared to FACS sorting. Cells were co-cultured with synovial ST-DC subsets sorted according to optimised CITEseq-guided strategy (Figure S12 and 13) in 96-well round-bottom cell-culture plates (3799 – SLS, Corning) in the presence of anti-CD3 at 0.25 μg/ml (to mimic antigen stimulation) (BioLegend, #300438, clone UCHT1) and IL-15 at 20 ng/ml (PEPROTECH, #200-15) to provide a survival signal, at a ratio of 1:5 for 5 days in complete RPMI1640 medium. After 5 days in culture, changes in T-cell phenotype were investigated by evaluating the expression of a set of extracellular and intracellular receptors/cytokines and transcription factors by flow cytometry. Details of antibodies are provided in Table S10.

### Mapping DC subsets in synovial tissues using immunofluorescent staining

Formalin-fixed paraffin-embedded 5μm-thick synovial tissue sections were stained with antibodies directed against markers LAMP3 or AXL or CD1c or CD68 or CLEC10A or in combination, or with appropriate isotope control antibodies following previously published protocol^43^. Details of the primary and secondary antibodies used are provided **Table S11.** The sections were visualised with a Zeiss LSM 880 confocal microscope, using either a water immersion LD C-Apochromat ×40/aperture1aperture1.3, or an oil immersion Plan-Apochromat ×63/1.4 objectives, and images acquired using Zen Black software (Zeiss). All images were processed (brightness/contrast adjustment and background subtraction) using the same software.

### The *in vivo* model of disease flare in remission RA

Patient recruitment criteria and the study design of the BioRRA study^70,71^ are described briefly in the “Patient recruitment and management” Materials section. PBMCs isolated from anticoagulated peripheral blood from n=12 RA patients in sustained disease remission at baseline (at which treatment was withdrawn without tapering) and at the follow-up time point (disease flare or drug-free remission) were isolated by density centrifugation and collected into foetal calf serum (FCS) with 10% DMSO and stored at -150°C. On the day of myeloid cell isolation, PBMCs were carefully defrosted and all live PB myeloid cells were FACS-sorted based on their expression of HLADR and the absence of markers of T-, B- and NK-cells. Details of the antibodies are provided in **Table S12**. Cells from individual patients were tagged and processed into single-cell libraries using BD Rhapsody system as described in the BD_Rhapsody scRNASeq section.

### Sample Preparation for BD Rhapsody scRNAseq of DC-T Co-Culture_1 and BioRRA Cohort and PB DCs

Cells were labelled with unique sample identifier tags (Sample Tag 1-12) using the BD Human Single-Cell Sample Multiplexing Kit (633781/BD Bioscience) according to the manufacturer’s protocol. Cells were then loaded onto the scRNA-seq BD Rhapsody Cartridge using the BD Rhapsody Cartridge Reagent Kit (633731) according to the manufacturer’s protocol. Single-cell cDNA was prepared using the BD Rhapsody cDNA Kit (633773). This was followed by single-cell Tag library preparation kit (633774) and mRNA library preparation either for the BD Rhapsody Immune Response Panel (633750) (ST-DC/ T-Cell co-culture / BioRRA Cohort Experiments) or for the BD Rhapsody WTA (633801) (PB DCs). Libraries were sequenced using Illumina NextSeq 500 (Glasgow Polyomics).

### Analysis of all Single Cell RNA Sequencing Data

#### Raw data analysis

Read alignment and generation of count matrices from raw scRNAseq data of 10x Genomics platform was performed using the Cell Ranger (v7.0.0, with parameter “include-introns=false”) pipeline. The “cellranger count” tool was used to map the reads against the Human Genome (GRCh38-3.0.0) and performed UMI counting. For analysis of data from BDRhapsody platform, the sequencing reads were processed with BD Genomics Rhapsody Analysis Pipeline CWL (BioRRA cohort and synovial organoid data processed with v.1.0 and DC-T co-culture cohort with v.1.9.1). In some runs of the co-culture cohort where the read2 was too short for the pipeline; two random base-pairs were added. Reads were either mapped against the BD Rhapsody Immune Response Panel reference (BioRRA and Co-culture-1, or against GRCh38.p12 human genome reference (Organoid cohort). For the co-culture cohort, the expected cell number was defined with the “cellNum” parameter as defined in **Supplementary xls, *QC & Filtering.*** The Seurat package (4.0.3) in R was used to create Seurat objects for each dataset (CreateSeuratObject) either from CellRanger output containing the matrix.mtx, genes.tsv (or features.tsv), and barcodes.tsv files from 10XGenomics data or from the RSEC_MolsPerCell.csv file for BDRhapsody data. Ambient RNA was removed using SoupX^83^ (1.6.2) from 10XGenomics data. Cells of all datasets were filtered for number genes and UMIs, and those with whole transcriptome for % mitochondrial genes within three median absolute deviation (MAD) around the median population^84,85^. The data was normalized (NormalizeData) and the top 2000 variable genes were identified for all samples (FindVariableFeatures). Cell doublets were marked using DoubletFinder^86^ (2.0.3) on single objects, and clusters >25% doublets were removed after integration. Protein level information derived from antibody-derived tags (ADT) were added and normalized using centred log ratio transformation (CLR). Deconvolution of the hashtag/sample information was performed using cellhashR (1.0.3) and Souporcell (2.5) was used to improve the deconvolution using SNPs information. ***General data integration and clustering.*** Prior to integration all relevant samples were merged based on common features and re-processed as one Seurat object. The data was renormalized, adjusting the scale factor to the median number of counts, as provided in **Table S13**. Principle component analysis (PCA) was performed on identified variable features across all samples (FindVariableFeatures). Cell embeddings from the selected (as given in **Table S13**), principal components (PCs) were used in UMAP generation (RunUMAP) to allow for visual inspection of batch separation prior to integration. Integration was then performed using the Seurat wrapper function (RunHarmony, SeuratWrappers, 0.3.0) for Harmony^87^ integration (specific versions of harmony used for each integration given in **Table S13)**. Batch variables to be removed by integration and theta values were specified (group.by.vars, theta parameters), as given in **Table S13.** The resulting harmony-corrected PCA embeddings were then used for UMAP generation, and the selected principal components (harmony-corrected PCs) were visualized (RunUMAP). The same PCs were used to determine the k-nearest neighbours for each cell during SNN graph construction before clustering at the chosen resolution of (FindNeighbors, FindClusters) as in **Table S13.** Clusters were identified their expression of canonical marker genes (FeaturePlot) and identification of cluster-markers (FindAllMarkers, test.use=MAST). Such cluster markers were identified as genes with significant adjusted p-value of <0.05 (Bonferroni and multiple test correction) and expressed by greater than 40% of cells in the cluster (‘min.pct’ parameter 0.4). ***Isolation and Identification of PB and ST myeloid DC.*** The raw scRNAseq data from PB and ST with 10XGenomics platform were integrated with our previously published myeloid cell data set^43^ and a scRNAseq dataset of 10K healthy PBMCs from 10x Genomics. All previously published data were reprocessed using the same methods described above. Coarse cell types were identified as shown in **Figure S1** before all myeloid cell populations (CD14^pos^ monocyte, CD16^pos^ monocyte, broadly annotated synovial tissue macrophage and DC) were isolated from integrated PB and ST dataset. Selected cells (n= 70,471) were re-processed with pipeline described above and values supplied in **Table S13.** Myeloid cell clusters were identified based on previously described nomenclature^88^ and clusters found in analyses of PB alone (**Figure S2**). Next, we investigated which cells cluster with PB DCs and expressed classical myeloid DC markers, including the proteins CD11c and MHC-II as well as genes CLEC10A and CD1c **(Figure S3).** Clusters with high expression included CD1c^pos^ DCs, CCR7^pos^ DCs, and SDS^pos^NR4A3^pos^CXCR4^pos^ tissue DC cluster as well as a population of FOLR2^high^CLEC10A^pos^ STM. These populations and their potential PB predecessors were highlighted and isolated for further analysis. Selected cells (n= 37,725) were re-processed with same pipeline and re-visualization and clustering of these data identified additional intermediate clusters. We excluded macrophages by high expression of FOLR2 and C1QA RNA. The remaining cells were then isolated (n= 7869) and reanalysed with same methods described above and visualized in Figure 1. Cells were annotated based on trajectory analysis and well annotated DC2, DC3 and iDC3 markers. The exact flow of analysis is described in result section and illustrated in **Figures S2-11**. ***Isolation and Identification of ST CD4^pos^ T-Cells.*** CD4 T-Cell, CD8 T-Cell, and NK populations were isolated from integrated PB and ST dataset. Isolated cells were re-processed with pipeline described above. Contaminant cells were removed based on identified differentially expressed marker genes and samples with fewer than 42 remaining cells were excluded. Preprocessing, integration and clustering was repeated after removal of contaminants. CD4, CD8, and NK clusters were annotated based on differentially expressed marker genes and those annotated as NK or CD8^pos^ T-Cells were excluded, CD4^pos^ T-Cells were re-processed and clustered using given parameters (**Table S13).** Clusters were annotated based on differentially expressed marker genes, as described above, and guided by published ST T-cells data sets^54,65^. ***Reference annotation.*** scRNAseq data of CD4^pos^ T cell from ST-DC co-culture were integrated with the appropriate reference data (synovial tissue CD4^pos^ T cells) for annotation of clusters. The appropriate datasets were merged based on common features and integrated using method described above, adjusting the scale factor for normalization (**Table S13**) to account for read depth differences between platforms/experiments. Integrated clusters were annotated based on reference (synovial tissue) dataset clustering, by generating heatmaps of gene expression correlation matrix as well as a confusion matrix, illustrating the proportion of cells from original synovial tissue reference clusters within each of the new integrated clusters. ***Pathway Analysis.*** Pathway activity was inferred across selected ST-DC clusters using PROGENy (v1.24.0) as recommended in package scRNAseq vignette. Briefly, a progeny assay was created in the Seurat object using the progeny function, specifying the top 500 footprint genes per pathway, alongside num_perm=1, and scale=FALSE. The progeny assay scores were scaled using the Seurat ScaleData function. Mean scaled pathway activity was calculated by ST-DC cluster and visualised using the pheatmap package (v1.0.12). Cytokine and Cytokine receptor interaction were investigated in selected DC and CD4^pos^ T-Cell clusters by looking at genes from the KEGG_Cytokines and cytokine receptors pathway (hsa 04060). Genes from this pathway were specified in the ‘features’ parameter within the FindAllMarkers function of Seurat (5.0.1). For both DC and CD4^pos^ T-Cell cytokine pathway analysis, minimum logFC was specified as 0.5, DE genes were to be expressed in minimum 25% of cells in the cluster alongside being positively upregulated in the cluster, and p<0.05 based on MAST with Bonferroni correction for multiple comparison. The same parameters were applied where an individual clusters profile was assessed by synovial disease state. ***Cell Trajectory Analysis.*** Single-cell trajectory analysis (RNA velocity^89^) of ST myeloid DC and active RA CD4 T-cells clusters, and their potential peripheral blood precursors, was performed by estimation of spliced and unspliced counts using the velocyto command line interface (velocyto run10x). Generated .loom files for each sample, containing transcript splicing information, were incorporated into our analysed Seurat object by splitting the object by sample, loading each .loom file using the (ReadVelocity, SeuratWrappers (0.3.0)) and creating a new assay for spliced, unspliced and ambiguous counts before merging our samples back together again, recreating our integrated Seurat object. This Seurat object was then converted for application in python (SaveH5Seurat, Convert) using SeuratDisk (0.0.0.9019) package. The converted .h5ad file can then be read into python using the scanpy^90^ (1.9.3) package, which creates an AnnData (0.9.1) object (sc.read). The spliced and unspliced count data was normalized and pre-processed as recommended by scvelo^89^ (0.3.1) before running RNA velocity analysis. A PAGA^91^ graph was constructed (scv.tl.paga, scv.pl.paga) to illustrate cluster connectivities and RNA velocity is used to infer direction of identified PAGA cluster transitions. Velocity pseudotime was estimated and genes were ranked by velocity to identify top differentially expressed unspliced genes for each cluster (rank_velocity_genes). ***Ligand-receptor interaction analysis.*** Inference of cellular communication was computed using the CellphoneDB^92^ package (v.5.0) using the *cpdb_statistical_analysis_method* ran in Python (v.3.8). The analysis was run using the following constraints; returned ligand/receptor genes must be expressed in at least 10% of all cells in a given cluster, with significant interactions being defined at a p-value of <0.05 after mean expression values of interacting clusters are subject to permutation. All other parameters were run as recommended. Predicted interactions were generated for all cell type clusters in the scRNA-seq dataset followed by a refined analysis limited to co-localised cell types of interest as reported in the spatial analysis. Significant interactions were visualised using the R (v.4.2.2) implementation of ktplots (v2.3.0) and additional custom visualisation scripts. ***Predictive Module Score Analysis (BioRRA Cohort).*** Annotation of clusters in BioRRA cohort (targeted immune gene panel) was guided by integration with whole transcriptome PB data. Differentially expressed genes upregulated in DC2, DC3, and iDC3 clusters at baseline of patients who go on to flare upon treatment withdrawal were identified using the FindMarkers function (only.pos = TRUE, logfc.threshold = 0.25). This process was repeated for genes upregulated at endpoint flare versus endpoint sustained remission. Shared genes between these two tests were identified for generation of our DC flare-associated module score. To generate this score, the AddModuleScore function from the Seurat R package was utilised with parameters (features=list(DC Flare Associated Genes), ctrl=10, name=’DC Flare Module Score’, pool=TRUE). This module score was calculated for DC2, DC3, and iDC3 clusters, and mean expression of the module scores by patient and timepoint were utilised in AUC-ROC analyses.

### MNP-Verse analysis

We utilized the AddModuleScore function from the Seurat R package to compute module scores for feature expression programs in single cells. The genes belonging to each cluster were selected using the FindAllMarkers function (with parameters “min.pct = 0.4, logfc.threshold = 0.7, test.use = “MAST”, only.pos = T”). Subsequently, the score for each cluster was computed and integrated into the MNPverse Seurat object, and scores for the ST DC2 LAMP3^pos^CCR7^pos^ were visualised. Harmony (1.2.0) integration of our myeloid cell dataset from RA PB and ST with MNPverse dataset was performed. Common features between the two datasets were selected, and we used the median number of counts for scaling factor for normalization. We corrected for batch variables including sample donor, experimental differences, and tissue variations within the dataset (group.by.vars=c(“Unique_ID”,“Experiment”,“Tissue”), theta=c(0,10,0)).

### Mapping of ST myeloid DC in situ with CosMx single cell spatial transcriptomics

We used the Nanostring CosMx Spatial Molecular Imaging platform to measure expression of 960 genes discriminating transcriptional profiles and spatial localization of 127,199 cells (69 fields-of-view (FOV)) in paraffin-embedded synovial biopsies from 3 active naive to treatment and 3 RA patients in sustained clinical and imaging remission (∼11 FOV per donor). Demographic, clinical and immunological characteristics of enrolled patients as well as synovitis degree of corresponding synovial tissues are described in **Table S4 and Supplementary Material_xls file**. ***Cell Segmentation.*** Initial image segmentation was performed with Mesmer^61^ with the following parameters: mesmer_mode = “both”, scale = pixel size of the images. We used the cell boundaries estimated by Mesmer as a prior for refinement of the segmentation with Baysor^62^ based on transcript densities, using the R wrapper (https://github.com/korsunskylab/baysorrr). Following successful cell assignment, we generated a gene-cell expression matrix and performed quality control, removing any cells with less than 30 counts and/or expression of less than 20 genes. Additionally, cells with radius less than 2 µm were also removed. Cells which passed QC filtering were then annotated using pipeline for cell type labelling described in Chen et al. (2024)^93^. ***Coarse cell type annotation.*** Briefly, read counts were normalized and log-transformed to median total counts of all cells remaining after filtering. PCA was performed and embeddings were corrected by integration with Harmony^87^ (0.1.0), specifying sigma value of 0.25 and theta values of 0 for both, sample run and FOV batch variables. Harmony corrected PC-embeddings were used to generate two-dimensional UMAP^94^ (uwot 0.1.16) and cell clusters were identified by shared nearest neighbour (SNN) modularity clustering. Clusters of coarse cell types were annotated based on marker genes identified by differential expression analysis performed using presto wrapper (1.0.0, https://github.com/immunogenomics/presto/tree/glmm/) for Generalised Linear Mixed Model (GLMM) estimation with lme4 (1.1-34) as described in Chen et al. (2024)^93^. Genes were considered significant when adjusted p value was less than 0.01 and an average logFC more than 0.5. ***Preparation of scRNAseq for reference annotation of spatial data*.** Following coarse cell type annotation, CosMx data were integrated with our synovial tissue scRNAseq dataset for reference annotation of subclusters. In preparation for this, our synovial tissue scRNAseq dataset was refined by removing genes that are not present in the CosMx SMI gene (n=922) panel. The data were then re-filtered to remove cells that now have low number of counts/features due to reduced gene panel. Variable features were identified, and the data was renormalized, adjusting the scale factor to account for reduced number of counts (median = 1255). We then followed standard pipeline for Seurat pre-processing and clustering of scRNAseq data, as described above, for coarse cell type annotation. Coarse cell type populations of interest were isolated, and as before, data was re-integrated, a new UMAP generated, and re-clustered with reduced gene panel. Any clusters that were indistinguishable with CosMx gene panel were removed. Relevant genes for population of interest arranged by z-score (presto, 1.0.0) and we ran sensitivity analysis by running the pipeline for integration and clustering using from 50-900 genes top variable genes and selecting the minimum number of genes necessary to distinguish our described DC and T-cell subsets. We then harmonized and clustered with minimum relevant genes selected from sensitivity analysis and annotated clusters based on correlation of gene expression with original annotations. Cells with clashing labels were removed and the number of cells per cluster was down sampled to median number cells per cluster. ***Reference annotation of CosMx spatial data with refined scRNAseq data.*** Each population of interest (Myeloid, Stromal, Endothelial and T Lymphocyte) from the spatial data was isolated based on coarse cell type annotation for integration with the appropriate scRNAseq reference. CosMx data was reduced to genes selected from sensitivity analysis in preparation of scRNAseq reference for that cell type. This allowed us to minimize noise and focus only on minimum genes necessary to define clusters. The data was then merged and renormalized adjusting the scale factor to account for reduced number of counts between both CosMx and scRNAseq dataset before following standard harmony pipeline for integration across modalities, accounting for source of the data (spatial/scRNAseq, theta=2) and sample ID (donor/sequencing run, theta=0) as batch variables. The integrated dataset was then re-clustered and new integrated clusters were identified. To do so, a heatmap of correlation matrix comparing the marker genes of new clusters with marker genes of the original single cell clusters was visualized. We also generated a confusion matrix – a heatmap illustrating the frequency of cells from original single cell reference clusters within each of the new integrated clusters. In the case that the new integrated cross-modality clusters contained multiple of scRNAseq reference clusters we performed subclustering and revisualization of gene correlation matrix and confusion matrix. Fractions of cells of each cluster from different sources was also visualized as stacked bar plot to identify any populations unique to CosMx spatial technology. The new integrated clusters were automatically reference annotated using the gene correlation matrix, annotating new clusters with the name scRNAseq reference cluster with the highest correlation of gene expression. This reference annotation was also performed manually, and results compared to finalize annotations before transferring new cell labels. Once all coarse cell populations of interest in spatial transcriptomic were isolated, integrated with scRNAseq reference, re-clustered and annotated, the new fine type cell annotations were transferred to the original CosMx spatial dataset containing all cell types. Spatial localization of coarse and fine type cell annotations were plotted using ggplot2 (geom_sf) allowing for visualization of cell geometries identified from segmentation (described above) manipulated using sf package (1.0.16).

### Niche and colocalization analysis of CosMx spatial transcriptomic data

To do spatial segmentation we first identify low-quality regions within the tissue, performing the following steps: (1) FOV region annotation and gridding, (2) spatial smoothing, and (3) dimensional reduction and clustering. ***FOV region annotation and gridding.*** We gridded the cellular region of each FOV by performing Voronoi tessellation on the cell centroids with the FOV boundary as the bounding box. Voronoi tessellation divides the space such that:

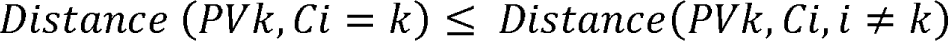

where PVk is any point P(x,y) in the Voronoi region Vk, and Ci is the centroid of the Voronoi region Vi. Because Voronoi tessellation grids the whole FOV irrespective of empty spaces within the tissue, we chose to perform Voronoi tessellation only between cells that are less than 50 µm apart from at least one other cell. Cells that are over 50µm apart from other cells are included in the analysis but with their original cell polygons instead of Voronoi regions. For most of the FOVs, we observed a gap between the last layer of cells and the FOV boundary. This led to edge effects where the cells closer to the edges had elongated shapes. To correct this, we changed the shapes of Voronoi regions of the edge cells to an intersection between a circular buffer of 15µm from the cell centroid of the boundary cells and the corresponding Voronoi region. This marked the end of gridding of the cellular region of the tissue. We merged all the Voronoi regions in each FOV and annotated it as “tissue”. We determined “glass” regions in each FOV by finding the non-intersecting region between the bounding box of the FOV and a 30µm buffered tissue region of the FOV. We buffered the tissue region to ensure we didn’t capture probes in the boundary regions between glass and tissue. Our rationale behind ignoring boundary transcripts is that these probes could belong to cells but were not assigned to cells due to segmentation errors. Grouping these into “glass” regions could skew our background identification. We then tiled the glass region of the FOV into 4-sided polygons that contain the same number of transcripts as the mean number of transcripts per Voronoi region in that FOV. ***Spatial smoothing.*** To construct the gene expression matrix of the tissue region, we mapped only the transcripts (both positive and negative probes) assigned to cells during segmentation to Voronoi regions. Because negative probes are excluded during cell segmentation, we assigned negative probes to cells by assigning a cell ID to a negative probe if it was within a cell boundary and 0 otherwise. To construct the glass region’s gene expression matrix, we used the “st_intersect” function to map transcripts to the glass tiles. We then combined both expression matrices to build a gene-polygon matrix for each FOV. From this point on, we will refer to both the Voronoi regions and the glass tiles as “polygons” and original cell shapes as “cell polygons”. To perform spatial smoothing, we ensured each cell captures a fraction of its neighbors (in addition to all transcripts from itself) in a diffusion-based method controlling for how aggressively we borrow transcripts (l) from our neighbors and how many degrees of neighbors we want to borrow transcripts from (k). The first step of spatial smoothing is to construct an adjacency matrix. We did that by constructing an unweighted Delaunay graph on the polygon centroids and pruning the edges between tissue and glass polygons. Pruning is important because our goal was to identify regions in the tissue that have similar gene expression profiles as glass, and borrowing transcripts from glass would make some tissue regions look like glass because of smoothing and not because they are low quality. After calculating the adjacency matrix, we smoothed it by diffusion process where the smoothed matrix M is calculated as:

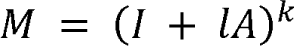

Where I is the Identity matrix, l is the rate of diffusion, A is the adjacency matrix, and k is the number of steps of diffusion. We row-normalized the smoothed matrix and built the smoothed gene expression matrix (G) as:

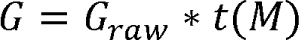

#### Dimension reduction and clustering

We then performed log-normalization, scaling, weighted-PCA, Harmony to correct for batch effects (sigma = 0.2, batch variables = SampleID, SampleFOV, nPCs = 20), UMAP, and clustering as described in the cell type labeling section to identify the tissue regions clustering with glass regions. These regions were labeled “low-quality” regions and removed from the analysis. ***Region annotation*.** To identify regions, we perform spatial smoothing, dimension reduction and clustering as described above on high-quality tissue regions. Clusters are annotated based on their cell composition. Furthermore, we performed colocalization analysis to define organization of cell subsets within the described tissue niches. Applying a permutation approach, as in Chen et al.^93^, we identified nearest neighbours and then randomized the positions of cells surrounding the defined cell type of interest and determined whether or not the colocalization of two subsets was not expected by chance. Significant colocalizations (adjusted p value < 0.05) were plotted as Z-score.

### PRIME cells data set analysis

The PRIME cell data set^72^ encompasses over 364 time points, both preceding and during eight flares, spanning a period of 4 years. Samples from this dataset which had conflicting metadata information or did not have information denoting weeks to flare were excluded. Raw readcounts were loaded and analysed using DESeq2 (1.40.2), transformed with VST, batch effects were evaluated using PCAtools (2.12.0) and readcounts were adjusted to remove batch of sequencing and occurrence of flare timepoint batch effect using Combat/SVA (3.48.0). Average expression per timepoint of batch-adjusted read counts for each DC signature gene was calculated using the avearrays function (limma 3.56.2). Average gene expression was scaled by row, and data was visualised using pheatmap (1.0.12). Module scores for DC signature genes were calculated on batch-adjusted read counts per timepoint using GSVA (1.48.3). Average GSVA module scores per timepoint were calculated using the avearrays function (limma 3.56.2) and visualised using ggplot2 (2_3.4.4).

### MIR155 expression and experimental overexpression of MIR155 in ex-vivo peripheral blood (PB) DC2 cells

#### MIR155 expression

DC2 from healthy and active RA PB, and synovial tissue from active RA were sorted based on negative expression of cell lineage markers (CD3, CD19/20, CD56, CD15 and CD117) and high expression of CD1c into tubes with microRNA preservation buffer from miRNeasy micro-Kit (217084, Qiagen). Clinical information for these patients is in **Table S5.** To evaluate MIR155 and housekeeping control, RNU6 expression, RNA was transcribed into cDNA and amplified with miScript Reverse Transcription Kit II (218161, Qiagen) and miScript PreAMP PCR Kit (331451, Qiagen), respectively. The miScript primer assays (Qiagen) were used for semi-quantitative determination of expression of U6B snRNA (MS00033740) and MIR155 (MS00031486) in combination with miRScript SybR Green PCR kit (1046470, Qiagen). The expression of genes of interest was presented as a relative value 2^-ΔCT^, where ΔCt is the Ct (Cycle threshold) for RNU6 (housekeeping genes) minus the Ct for the gene of interest.

#### Ex-vivo MIR155 overexpression

DC2 (CD1c^high^) from PB of active RA patients (n=12) were FACS-sorted into tubes with complete RPMI1640 media and seeded overnight in flat bottom 96-wells plates at a density of 10×10^3^ cells/well. The next day, cells were transfected using the Dharmafect 3 transfection system (T-2003-02, lot 00662107, Dharmacon) with either hsa-miR-155 mimic (C300647-05-305, lot 180510, Dharmacon), negative control miRNA mimic (CN-001000-01-05, lot 2145003, Dharmacon), or negative control labelled with 20nM Dy547 fluorochrome (CP-004500-01-05, lot 2054853, Dharmacon). After 4h, cells were either left unstimulated as controls, or were stimulated with LPS (100ng/mL, L6529, Sigma) for 48h. Culture supernatants were collected for soluble mediator analysis. Transfection efficiency was estimated based on the proportion of cells that were successfully transfected with the Dy547 mimic, and experiments where the transfection efficiency was below 60% were discarded. Cytokine concentrations in the culture supernatants of the RA DC2 were quantified using a predesigned high-sensitivity Luminex 100^TM^ Multiplex Kit (Millipore UK) on a Bio-Plex system (Bio-Rad).

### Phenotyping DCs in mesenteric lymph nodes (mLN) in wild type and miR-155 gene deficient mice

The mesenteric lymph nodes (mLN) from 8–12-week-old C57BL/6J (WT) and congenic Cg-Mir155tm1Rsky/J (miR-155 deficient) were harvested and digested with 1mg/mL of Collagenase D for 40 min in a 37°C shaking incubator at 150 RPM speed. After neutralisation of collagenase with complete media, cells were incubated with antibodies indicated in **Table S14)** for 30min at 4°C and acquired by FACS AriaIII. Data was analysed using FlowJo (Version 10.7.1).

### Isolation and analysis of mouse DC2 from gut, lymph and draining lymph nodes

To obtain thoracic duct lymph, mesenteric lymphadenectomy was performed on 6-week-old male C57/Bl6 mice by blunt dissection at laparotomy. The thoracic lymph duct was cannulated with a polyurethane cannula (2Fr). Lymph was collected in PBS / 20 U/mL of heparin sodium on ice, overnight as we described previously^95,96^. Matched small intestine was isolated and digested as described previously^95,96^. Lymph DC2s were identified as MHC II^hi^ CD11c^+^ B220^-^ CD11b^+^. In the mLN and intestine, cDC2s were identified as F4/80^lo^ MHC II^hi^ CD11c^+^ CD11b^+^. DC2 from different compartments were sorted (>100,000 cells). Information on antibodies used is provided in **Table S14**. RNA was isolated using RNAeasy kit (217084, Qiagen). cDNA samples were then run using the Agilent-066423 design and the 048306On1M (4×180K) array. Information on data deposition is provided in^96^.

### Data Availability

All scRNAseq and spatial data raw files are available from ArrayExpress under accession numbers E-MTAB-14191 (Co-Culture scRNAseq), E-MTAB-14169 (BioRRA scRNAseq), E-MTAB-14192 (Paired PB/ST scRNAseq), E-MTAB-14213 (healthy, active RA and remission ST scRNA/CITEseq), E-MTAB-14198 (Sort-seq), S-BSST1483 (CosMx Spatial Transcriptomics). Scripts for analysis of scRNAseq and published bulk RNA seq datasets are available at https://github.com/domenico-somma/MacDonald_et_al_Immunity_2024. Scripts for analysis of spatial transcriptomics data are available at https://github.com/DrLucyMac/spatial_synoDC.

### Statistical analysis

Detailed statistical methods are provided in each Figure legend and in the scRNAseq method sections above.

## Supplemental information index

**Document S1**: Supplementary Figures 1-20 and Supplementary Table 1-14.

---

**Supplementary Figure 1.**
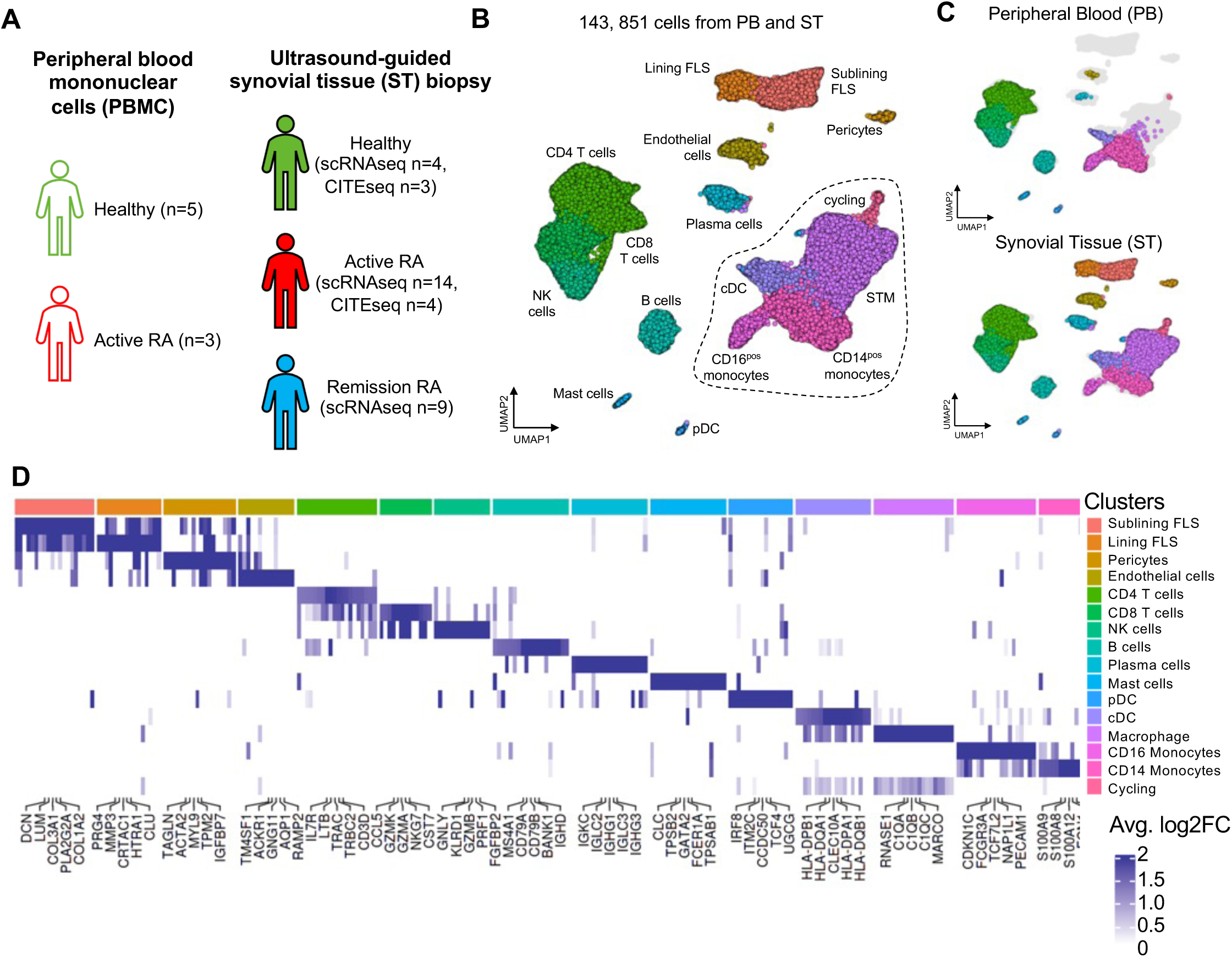
Integration of scRNAseq/CITE-seq data from peripheral blood (PB) and synovial tissue (ST) samples from healthy donors and patients with active RA and RA in sustained remission. **(A)** Illustration of the samples used in this study. **(B)** UMAP visualization of integration of 10x Genomics and BD Rhapsody whole transcriptome scRNAseq of peripheral blood mononuclear cells (PBMC, n= 53080 cells) and cells from synovial tissue (n= 90771 cells). Each point represents an individual cell (n=143,851) and points are coloured by coarse cell type annotation. **(C)** Split UMAP visualization illustrating the coarse cell types present in either PB or ST. **(D)** Heatmap visualization of the average log fold change (logFC) of the top 5 marker genes of annotated cell types. Differentially expressed (DE) genes were determined using MAST with Bonferroni correction for multiple comparisons and were considered significant if adjusted p value was less than 0.05 and they were expressed in more than 40% of cells in the appropriate cluster.

**Supplementary Figure 2.**
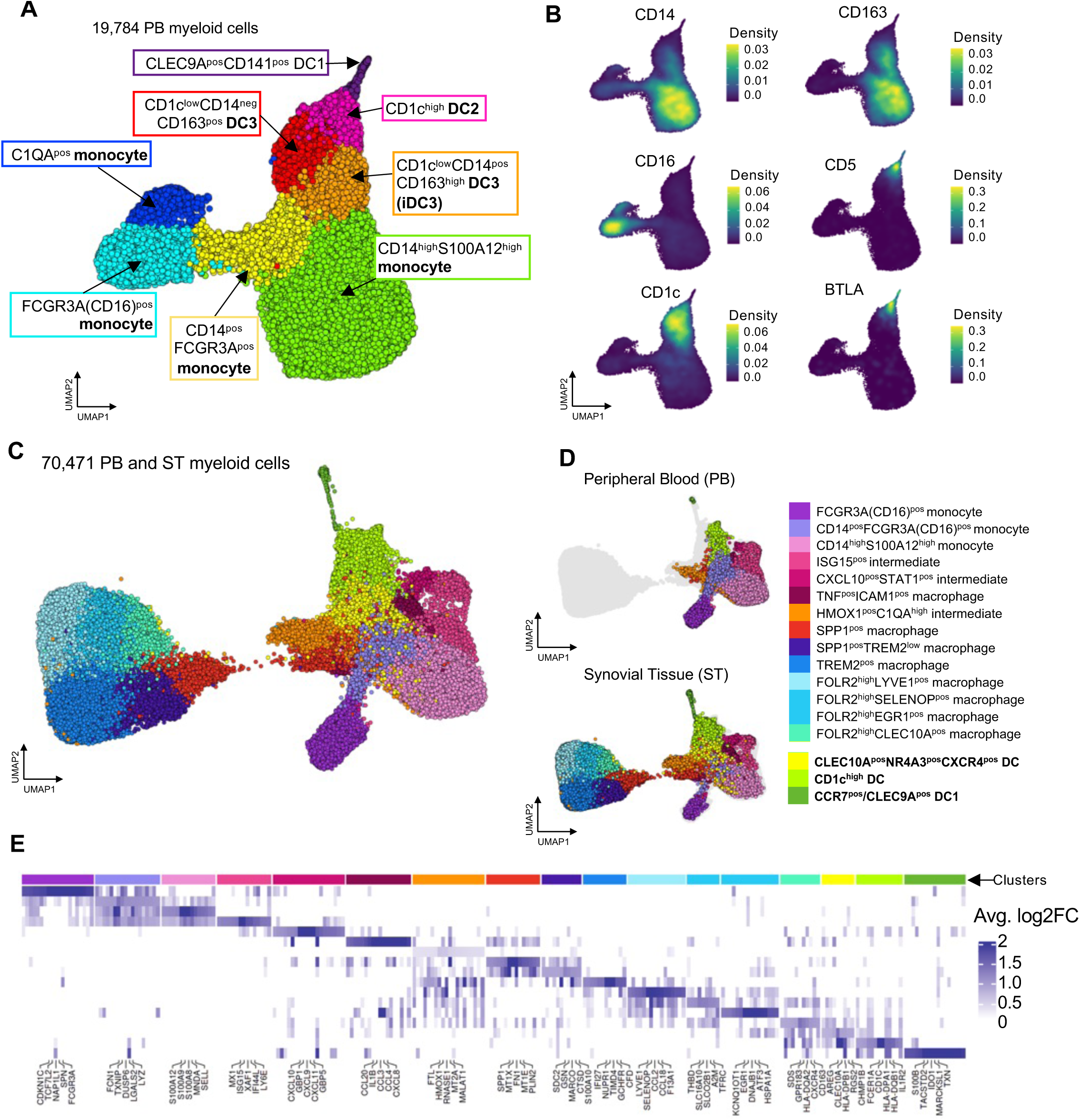
Integration of data from various scRNAseq platforms and sub-setting conventional PB/ST cells from total blood and synovial tissue data set. **(A)** UMAP visualization of Harmony integrated 10x Genomics and BD Rhapsody whole transcriptome and targeted immune panel scRNAseq of 19,784 PB myeloid cells (n=5 Healthy, n=3 Active RA and n=12 longitudinal remission to RA flare). Each point represents an individual cell and points are coloured by cluster annotation. **(B)** UMAP visualization of gene-weighted density estimation for expression of markers of PB DC and monocyte subsets. **(C)** UMAP visualization of integration of 10x Genomics and BD Rhapsody whole transcriptome scRNAseq of 70,471 myeloid cells from synovial tissue (ST, n=34) and peripheral blood (PB, n=8). Each point represents an individual cell and points are coloured by cluster annotation. **(D)** Split UMAP visualization illustrating the clusters present in either PB or ST. **(E)** Heatmap visualization of the average log fold change (logFC) of the top 5 marker genes. Differentially expressed (DE) genes were determined using MAST Bonferroni correction for multiple comparisons and were considered significant if adjusted p value was less than 0.05 and they were expressed in more than 40% of cells in the appropriate cluster.

**Supplementary Figure 3.**
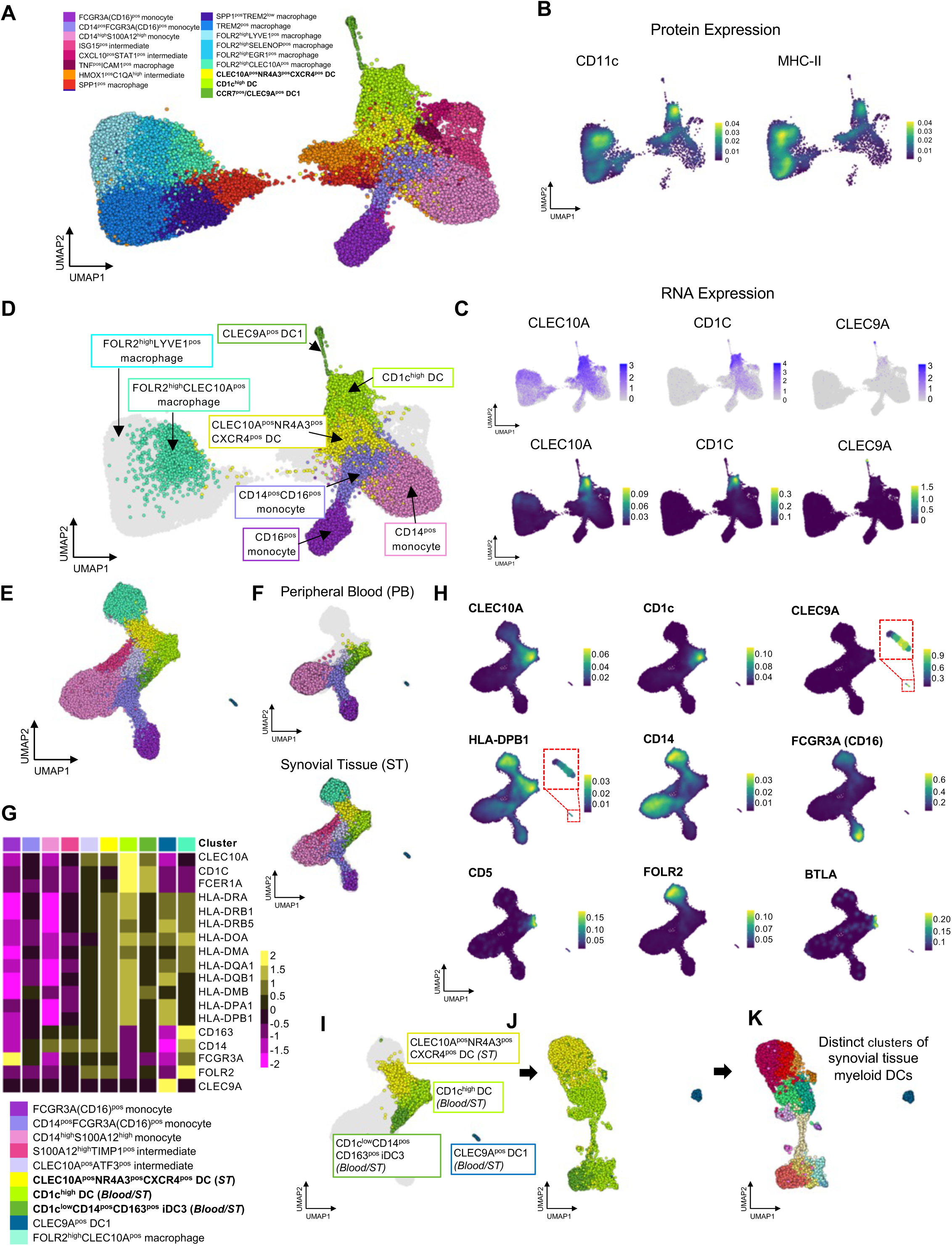
Identification of myeloid DC from Integrated scRNAseq/CITEseq PB and ST myeloid cells dataset of healthy donors and patients with active RA and RA in sustained remission. **(A)** UMAP visualization of the integration of 10x Genomics and BD Rhapsody whole-transcriptome scRNAseq of 70,471 myeloid cells from synovial tissue (ST) and peripheral blood (PB) as in Figure S1. Each point represents an individual cell, and points are coloured by broad cluster annotation. **(B)** UMAP visualization of embedding density of CD11c and MHC-II protein expression across all myeloid cells in CITE-seq dataset. **(C)** UMAP visualization of RNA expression level (top panel) and embedding density (bottom panel) of CLEC10A, CD1C and CLEC9A across all myeloid cells. **(D)** UMAP visualization of 70,471 PB and ST myeloid cells coloured by clusters with high expression of CLEC10A (RNA), CD11c (Protein) and MHCII (Protein) highlighted and selected for further analysis. **(E-F)** Re-clustering and re-annotation of myeloid cell clusters (n= 37,725) identified in D (E). Split UMAP visualization illustrating the clusters present in either PB or ST (F). **(G)** Pseudobulk heatmap illustrating average gene expression value of cluster markers and MHC-II associated genes of each cluster selected for further analysis in (D) based on high expression of CLEC10A (RNA), CD11c (Protein), and MHCII (Protein) and re-annotated in (E). **(H)** Embedding density visualization of RNA markers of STM (FOLR2), monocytes (CD14, FCGR3A, C5AR1), DC1 (CLEC9A) and myeloid DCs (CD5, BTLA, CD1c, CLEC10A) on UMAP from E enable to exclude macrophages from future clustering of myeloid DCs. **(I-J-K)** Selection of cell clusters (I) that cluster with PB DC2, DC3, and iDC3 and are different from monocytes/macrophages followed by unbiased re-clustering and illustration of where cells from (I) are located on the new UMAP (n= 7869) (J). Novel annotation of identified ST myeloid DC clusters on new UMAP visualization containing only myeloid DC and a small DC1 cluster (K).

**Supplementary Figure 4.**
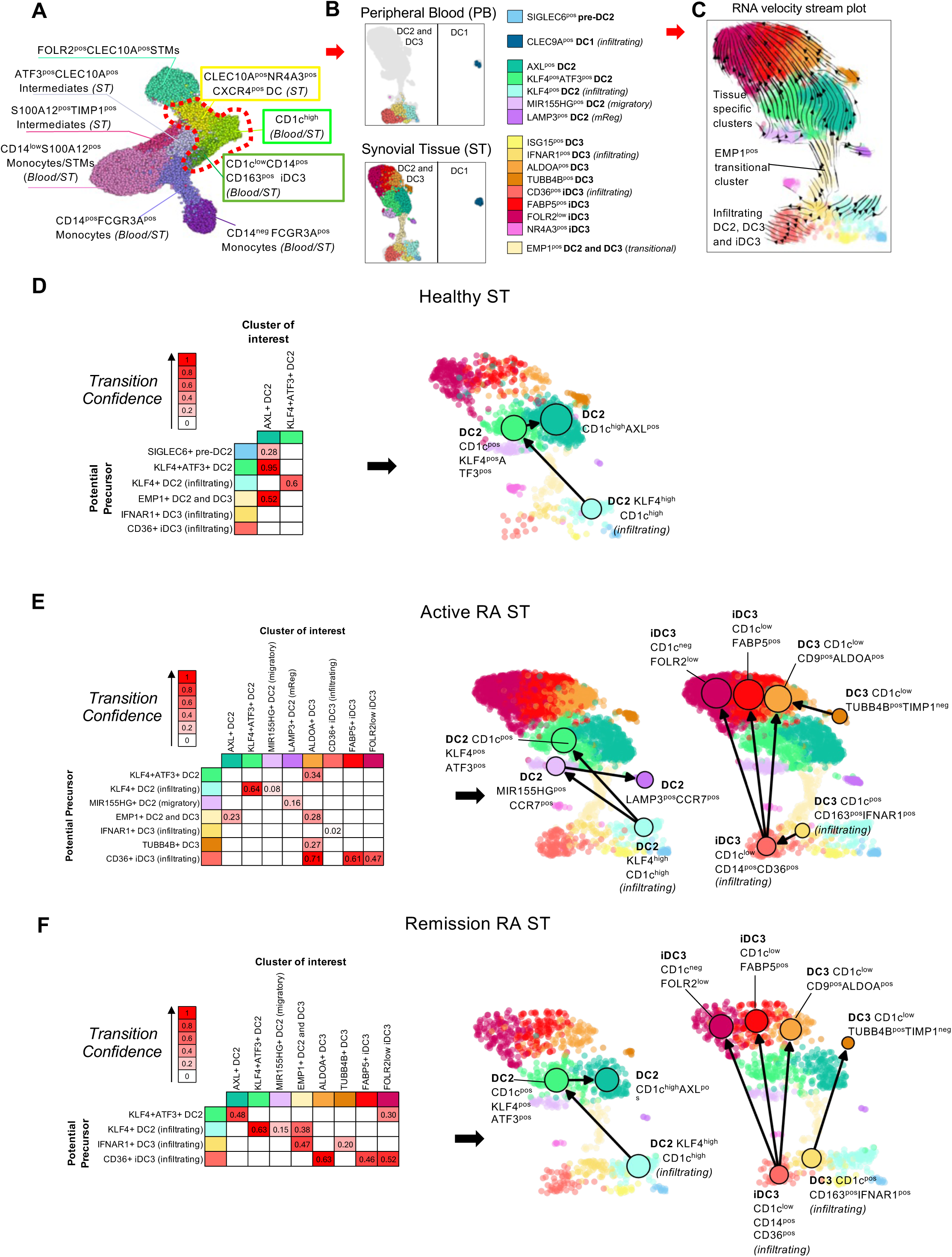
Differentiation trajectory of synovial tissue myeloid dendritic clusters from PB DC predecessors. **(A)** UMPA of an integrated dataset comprising PB myeloid DCs (DC2, DC3 and its iDC3 phenotype), monocytes, and ST myeloid cells expressing high levels of CLEC10A (RNA), CD11c (Protein), and MHCII (Protein) as in Figure S3E. **(B)** Split blood versus tissue UMAP illustrating 14 distinct myeloid DCs emerged from unbiased re-clustering of cell clusters from (A) (dotted red line) and a small cluster of DC1. **(C)** Single-cell trajectory analysis (RNA velocity stream plot) of synovial tissue (ST) myeloid dendritic cell (DC) clusters from peripheral blood (PB) DC2, DC3, and iDC3 to determine which ST DC cluster differentiates from which PB DC subset. The direction of arrows infers the path of cell trajectory based on spliced versus unspliced RNA counts. Pseudotime infers the ordering of cells along a lineage based on the cells’ changing gene expression profiles. Nomenclature provided is based on trajectories presented in panels D-F. **(D-F)** Single-cell trajectory analysis was performed by generation of RNA velocity directed PAGA connectivity graphs to illustrate cell state transitions in healthy (D), active RA (E), and remission RA (F). Confidence values between identified cell state transitions are given in table on the left with selected transitions visualized in UMAP on the right. We selected cell state transitions from the predecessor with the highest transition confidence for each cluster of interest. Trajectories leading to different myeloid ST-DC subsets are highlighted in separate UMAPs. Full transition confidence values are given in Supplementary Material in xls file.

**Supplementary Figure 5.**
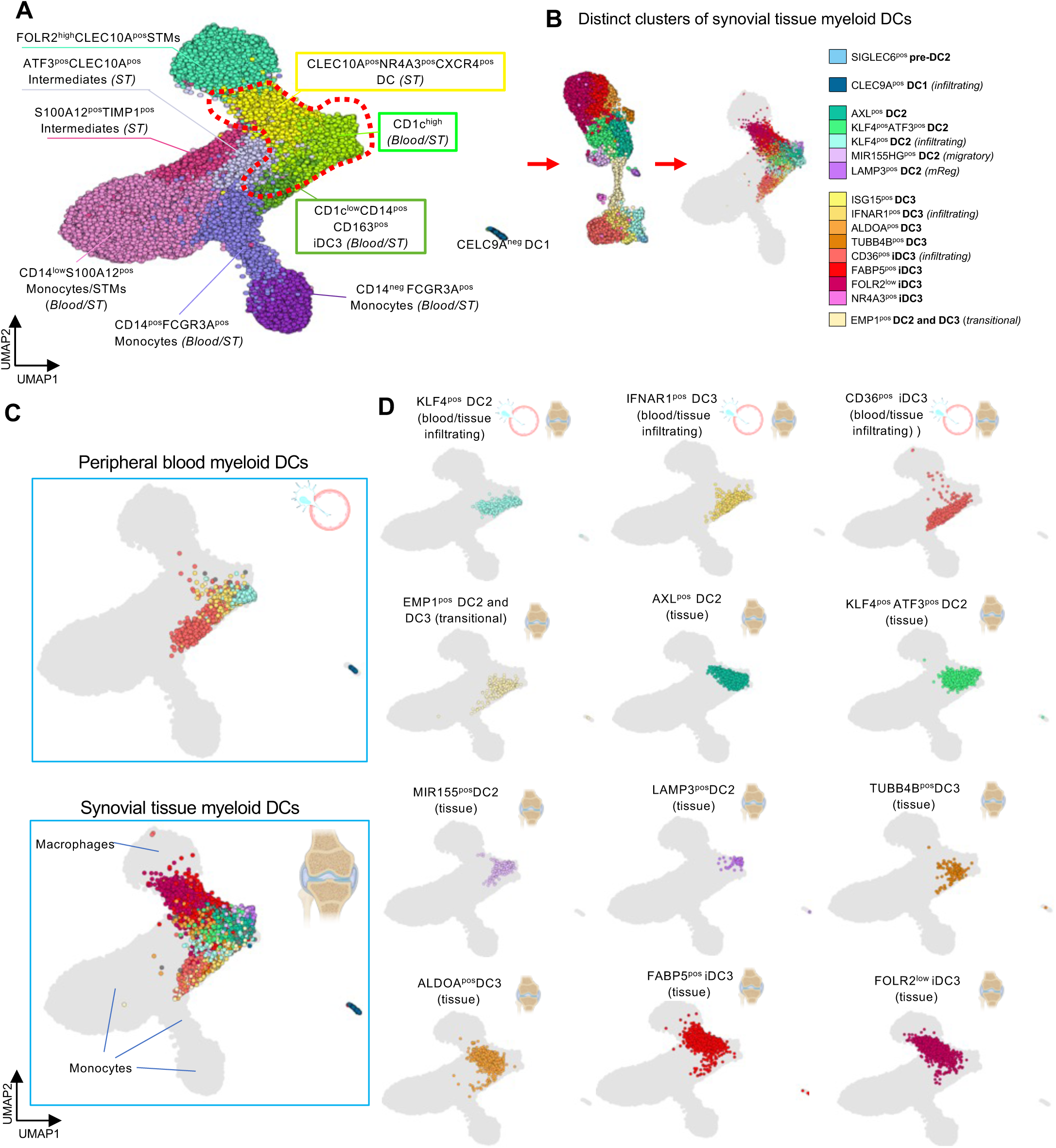
Illustration of myeloid ST-DC clusters in integrated PB/tissue myeloid cell data set. **(A)** UMPA of an integrated dataset comprising PB myeloid DCs (DC2, DC3 and its iDC3 phenotype) and monocytes, and ST myeloid cells expressing high levels of CLEC10A (RNA), CD11c (Protein), and MHCII (Protein) as in Figure S3E. **(B)** Unbiased re-clustering of cell clusters from (A) (dotted red line) showing 14 distinct phenotypic DC clusters of DC2, DC3, and iDC3 as in Figure S4B. These are then mapped back on the UMAP of PB/ST myeloid cells. **(C)** Split blood versus tissue UMAP of PB/ST myeloid cells visualizing 14 myeloid ST-DC clusters and showing that they do not cluster/overlap with macrophages and PB monocytes. **(D)** Visualization of each myeloid DC cluster on the UMAP as in C, indicating whether they are present in blood, tissue, or both, and showing no overlap with monocyte/macrophage clusters.

**Supplementary Figure 6.**
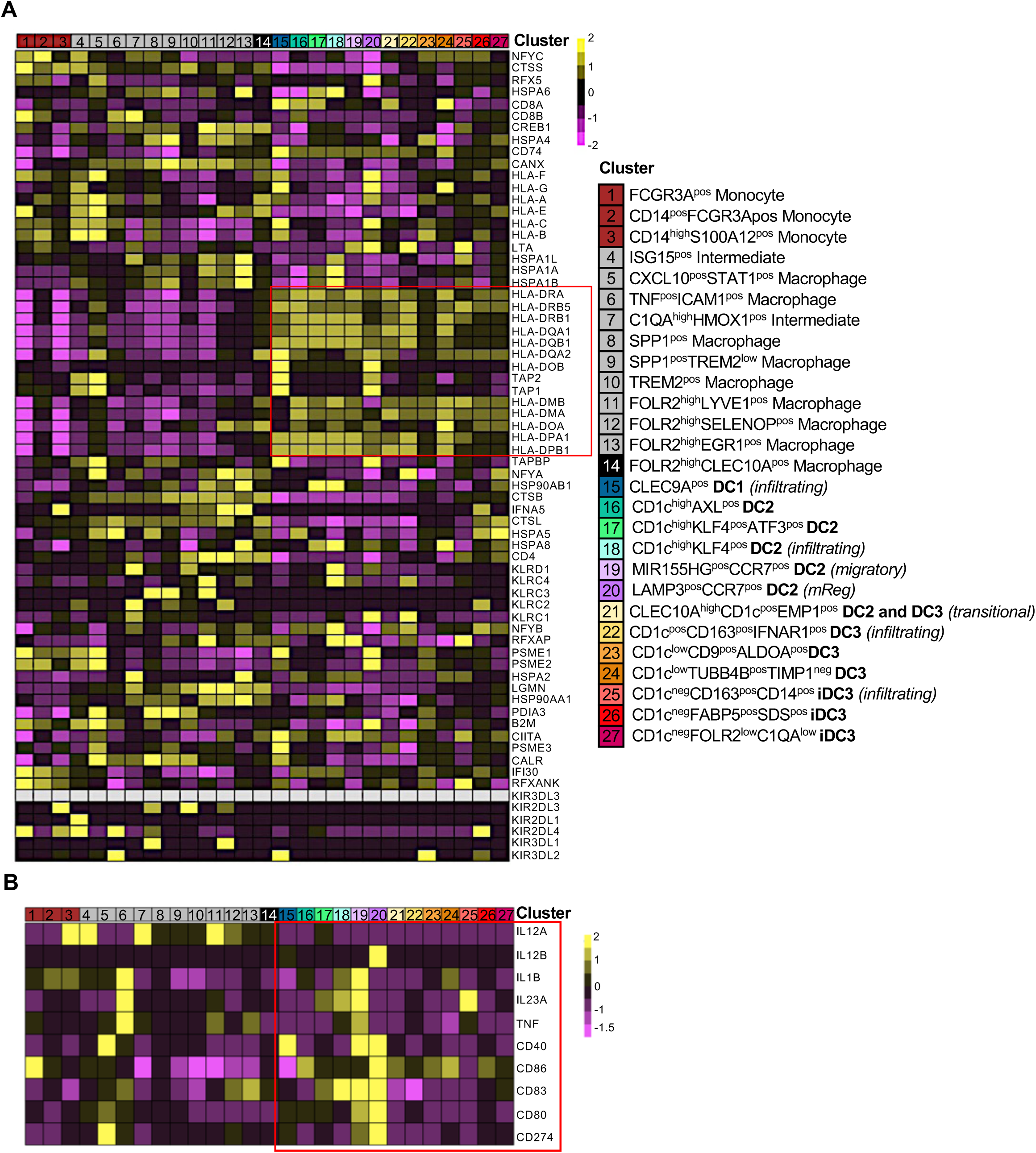
Antigen Processing and Presentation Gene Expression of ST Myeloid Cells. **(A)** Heatmap visualising scaled antigen presentation and processing genes from the KEGG antigen processing and presentation pathway across ST myeloid cells from Healthy (n=7), active RA (n=18), and remission RA (n=9) synovial tissue. **(B)** Heatmap visualising scaled expression of PB DC2 and DC3 -associated cytokines and co-stimulatory molecules across ST myeloid cells from Healthy (n=7), active RA (n=18), and remission RA (n=9) synovial tissue. Red boxes indicate all ST-DC clusters.

**Supplementary Figure 7.**
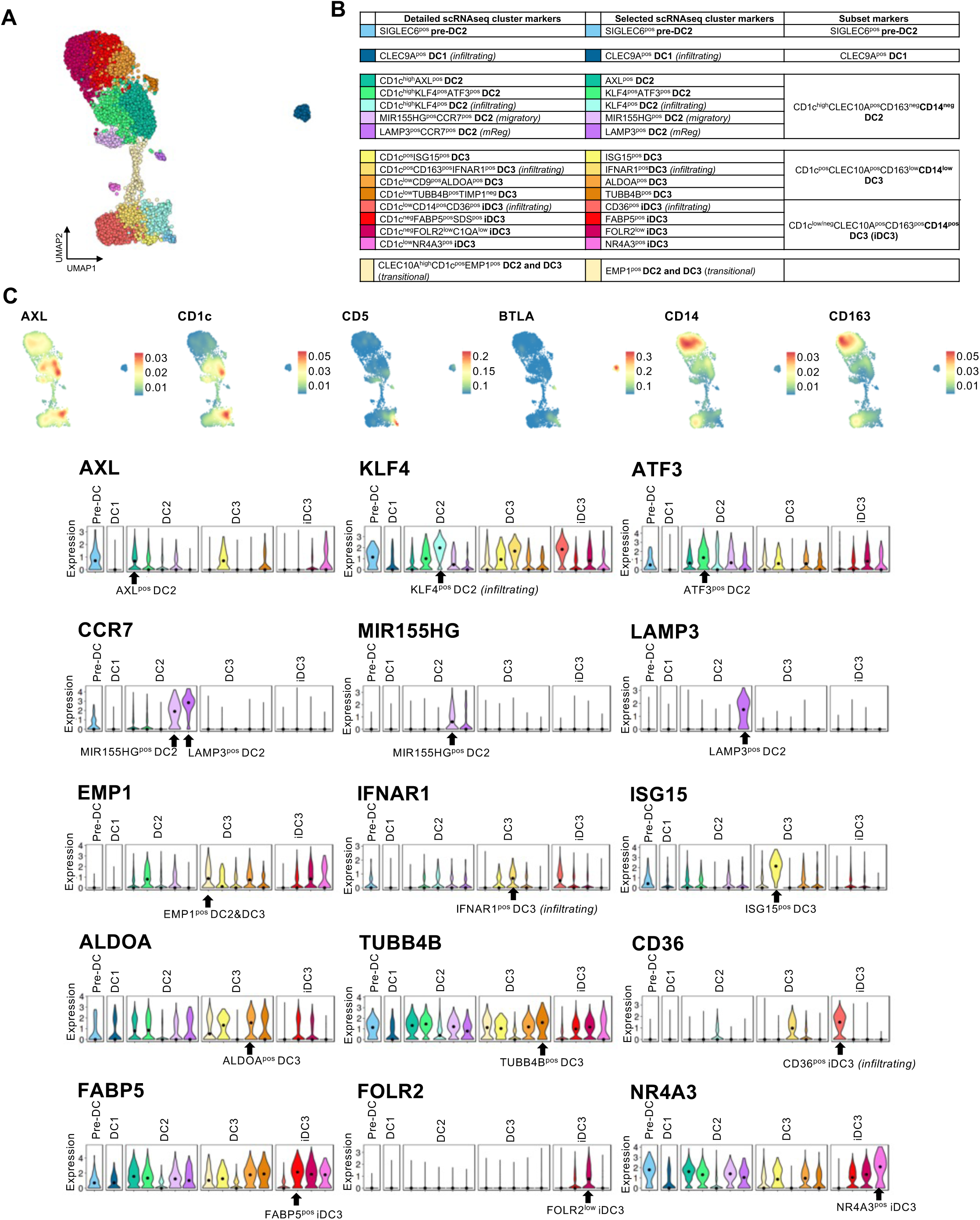
Single-cell omics identified phenotypically distinct clusters of synovial tissue myeloid DCs subsets. **(A)** UMAP visualization of integrated CITE-seq (n=7) and scRNAseq (n=35) data of myeloid DCs (n=7869) from synovial tissues (6510 cells) and blood (1359 cells). Each cell is represented by an individual point and is coloured by cluster identity based on trajectory analysis from blood precursors and its validation in synovial organoids. **(B)** Table summarizing scRNAseq markers of ST-DC subsets and their phenotypic clusters. **(C)** Density plots illustrate the expression of ST-DC2, ST-DC3, and ST-iDC3 subset markers. **(D)** Violin plots illustrate the expression of markers for different phenotypic clusters of ST-DC2, ST-DC3, and ST-iDC3.

**Supplementary Figure 8.**
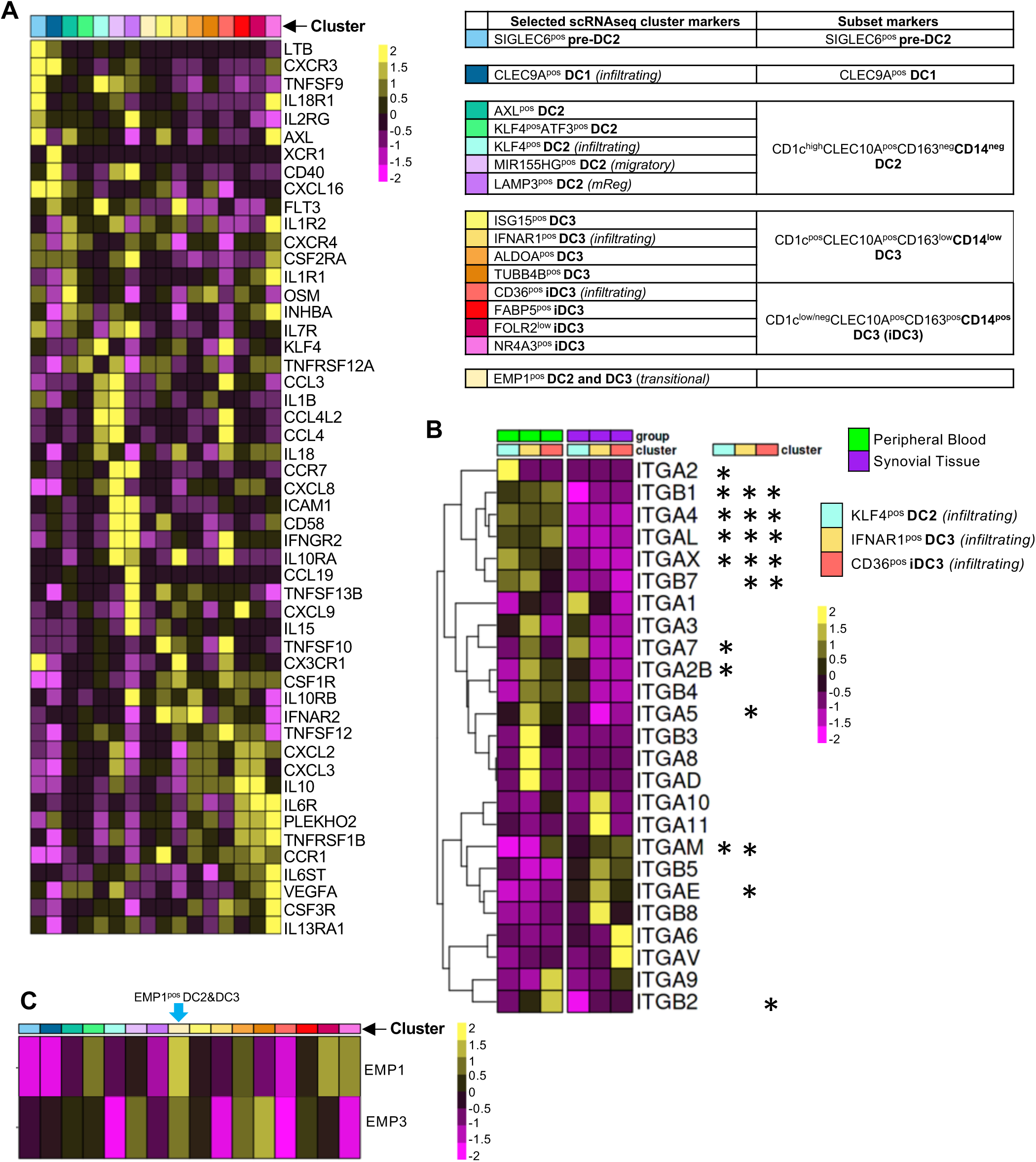
Cytokine/cytokine receptor and integrin profiles of synovial tissue dendritic cells. **(A)** Heatmap visualising scaled differentially expressed genes from the KEGG cytokines and cytokine receptors pathway between different synovial tissue (ST) dendritic cell clusters (Criteria: expressed in >25% of cells per DC cluster, with log-fold change >0.5, and p<0.05 based on MAST with Bonferroni correction for multiple comparison). Data from healthy (n=7), active RA (n=18), and RA in remission (n=9) ST. **(B)** Heatmap visualising scaled differentially expressed genes of integrin family between PB DC2, PB DC3, its iDC3 phenotype and tissue infiltrating counterparts: infiltrating ST KLF4^pos^ DC2, ST IFNAR1^pos^ DC3 and ST CD36^pos^ iDC3. Those integrins that are statistically significant (p<0.05 based on MAST with Bonferroni correction for multiple comparison) are indicated with asterisks. **(C)** Heatmap visualising scaled expression of EMP1 and EMP3 between all ST-DC clusters. EMP1 and EMP3 are marker genes of blood to tissue DC2 and DC3 transitional cluster (EMP1^pos^ DC2&3).

**Supplementary Figure 9.**
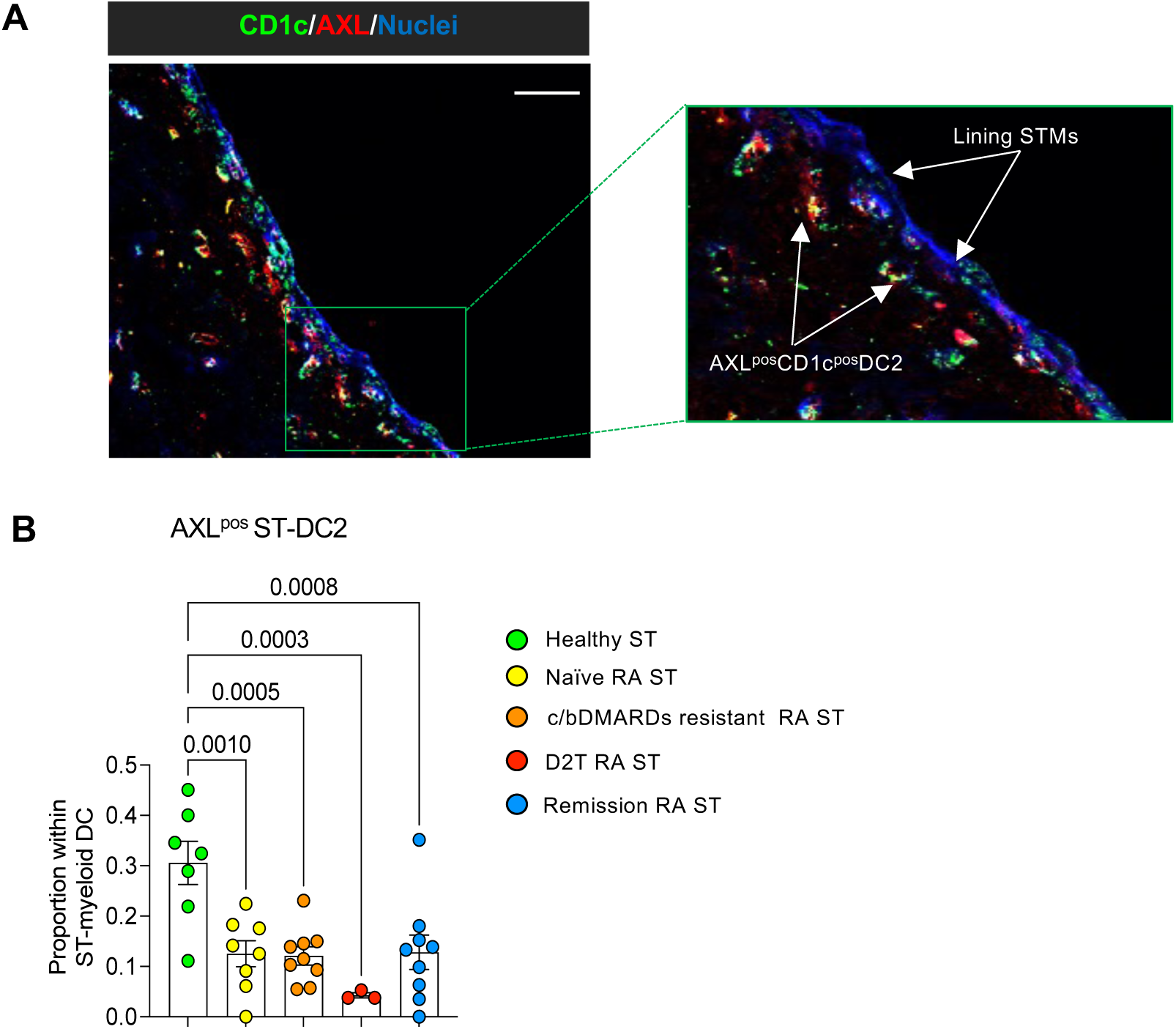
Difficult-to-treat RA shows the most profound decrease in tolerogenic AXL^pos^ DC2. **(A)** Representative confocal microscopy image showing immunofluorescence staining for CD1c (green), AXL (red), and nuclei (DAPI, blue) in healthy synovial tissues (ST) at 40x magnification. The insert on the right shows an enlarged view of the lining layer area. Images are representative of synovial tissue from healthy donors (n=5), active RA (n=6), and remission RA (n=5). Images are representative of synovial tissue from healthy donors (n=4). Scale bar = 50μm. **(B)** Difference in the proportion of ST AXL^pos^ DC2 cluster within the ST-DC pool between healthy controls and different phenotypes of active RA and remission RA based on ST scRNA-seq analysis. Data is presented as a boxplot with median and interquartile range. One-way ANOVA with Tukey corrections for multiple comparisons was used. The exact p-values are provided on the graphs. c/bDMARDs: Conventional or biological disease-modifying anti-inflammatory drugs, D2T: Difficult-to-treat (failed at least two biological therapies).

**Supplementary Figure 10.**
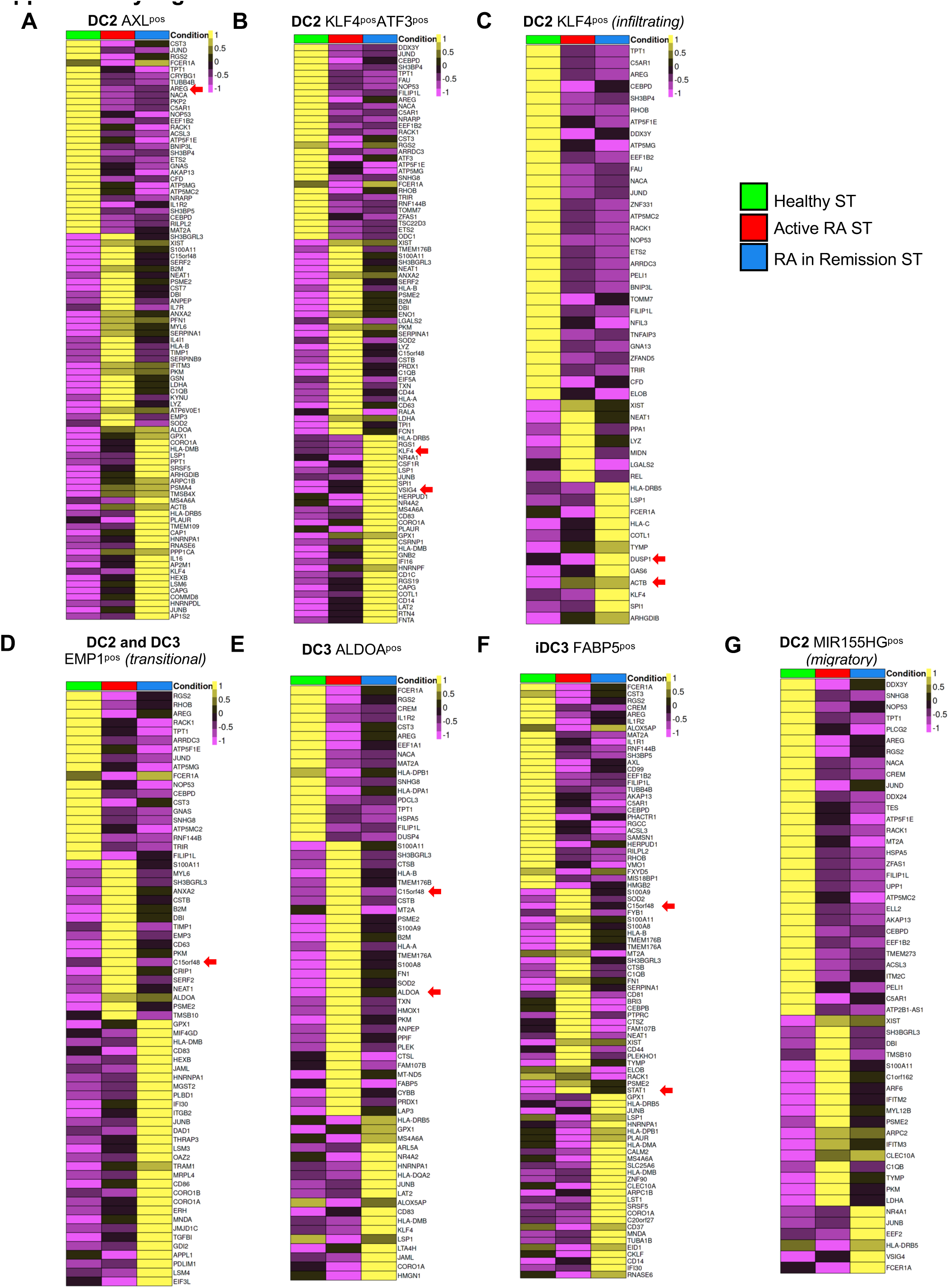
Joint immuno-state specific phenotypes of ST-DC clusters that differ in frequency between healthy, active RA and RA in remission. **(A-G)** Heatmaps visualising the top 30 scaled differentially expressed marker genes in healthy and RA synovium (expressed in >40% of cells per condition, with log-fold change >0.5, and p<0.05 based on MAST with Bonferroni correction for multiple comparison). Comparison based on specific cluster (annotated by colour above each heatmap) for Healthy (n=7), Active RA (n=18), and remission RA (n=9). Red arrows indicate gene of interest as discussed in the Results section.

**Supplementary Figure 11.**
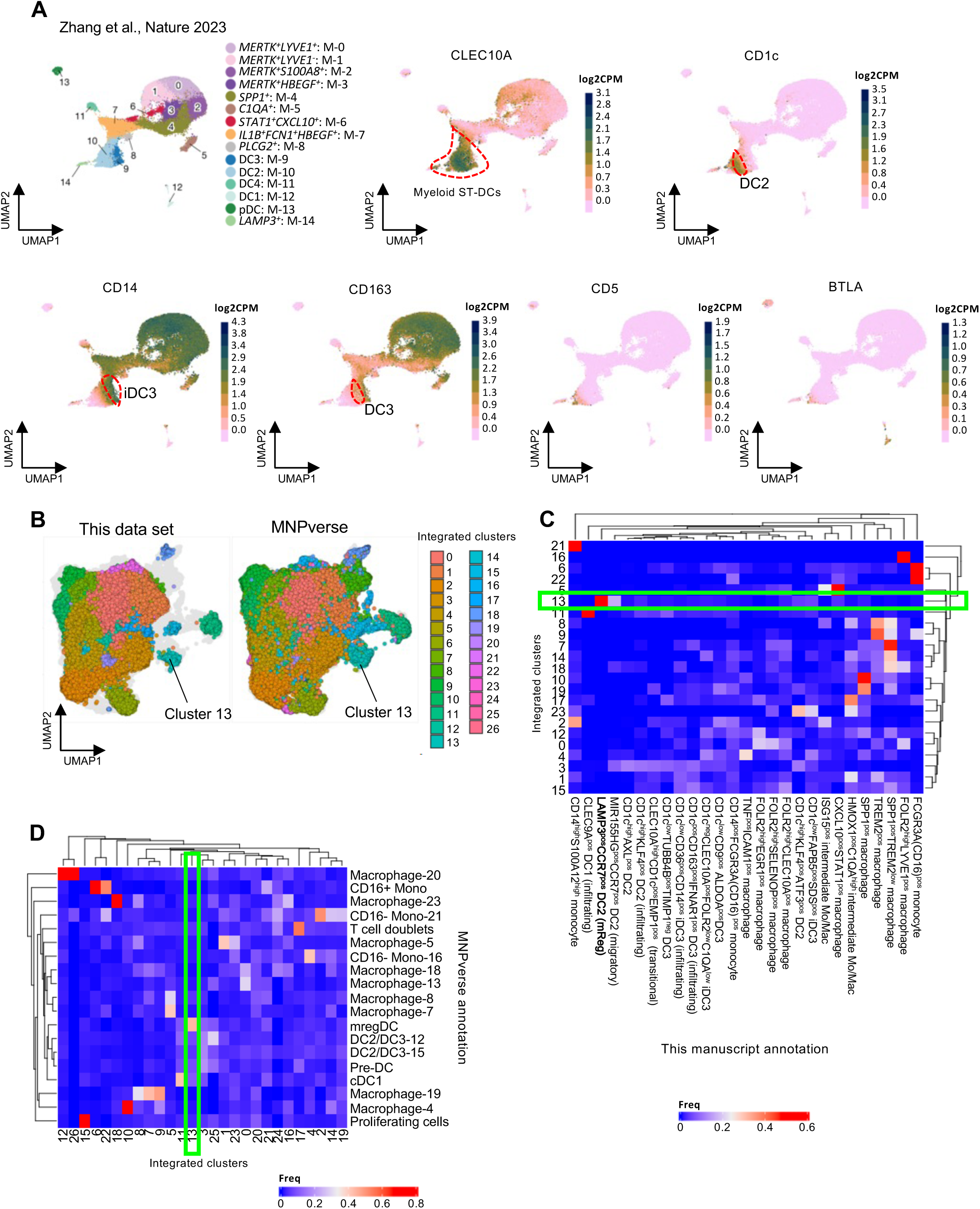
Validation of ST myeloid DC clusters with CTAP and MNPverse datasets. **(A)** Zhang et al (CTAP, Nature 2023) synovial myeloid data set of active RA shows presence of CLEC10A positive synovial myeloid DC subsets that include DC2 (CD1c^high^CD14^neg^CD163^neg^), and two subsets of DC3: CD1c^low^CD14^low^CD163^low^ and CD14^pos^CD163^pos^ inflammatory DC3 (iDC3) that clearly cluster separately from monocytes/macrophages, confirming data in the current manuscript. **(B)** Split UMAP visualisation of integrated scRNAseq/CITEseq data of myeloid cells (healthy (n=7), Active RA (n=18), and RA in Remission (n=9) from the current manuscript integrated with the MNPverse dataset. Cluster 13 correspond to mReg DC from MNPVerse and ST LAMP3^pos^CCR7^pos^ DC2. **(C-D)** Confusion-matrix showing the percentage of the original cell-annotation (y-axis) either in this manuscript (B) or in MNPverse (C) compared with the new integrated clustering. Cluster 13 is highlighted and corresponds to mReg DC/ ST LAMP3^pos^CCR7^pos^ DC2 in both datasets.

**Supplementary Figure 12.**
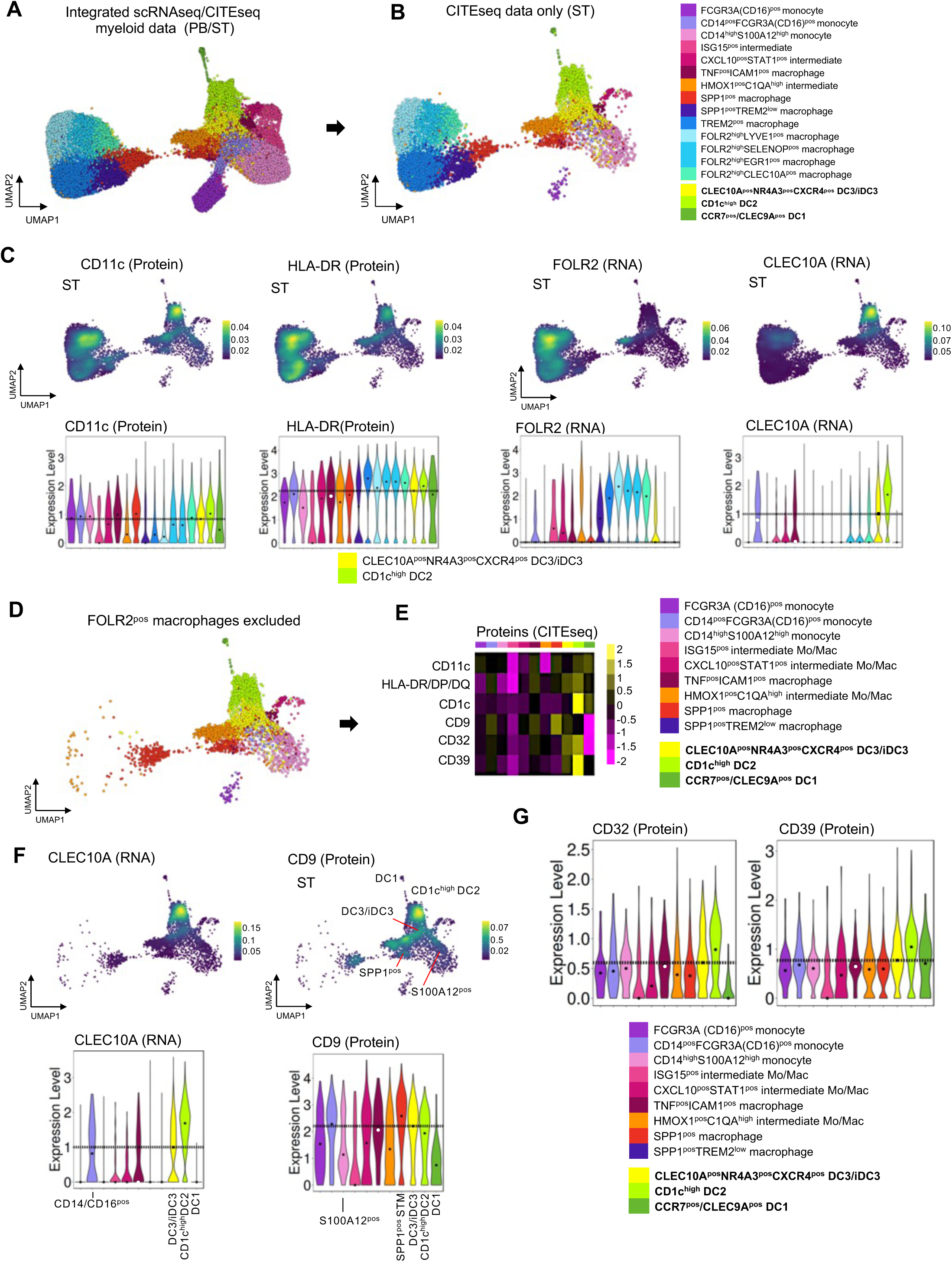
Establishing flow cytometry gating strategy for sorting ST-DC2 and ST-DC3/iDC3 based on CITEseq data. **(A-B)** UMAP visualization of integrated scRNAseq/CITE-seq PB/ST myeloid dataset (n=42 samples) (A), and UMAP visualizing subsampled CITEseq dataset (n=7) in B. **(C)** Expression density plots and violin plots visualizing protein/RNA expression of CD11c, HLA-DR, FOLR2, and CLEC10A suggest that gating cells with high expression of HLA-DR and CD11c captures all myeloid DC clusters while excluding tissue-infiltrating CD14^pos^S100A12^pos^ and CD16^pos^ monocytes and some synovial tissue macrophage (STM) populations. All tissue-resident STM can then be excluded by high expression of FOLR2. ST-DC2 and ST-DC3/iDC3 can subsequently be gated out of the remaining contamination of infiltrating monocytes and monocyte-macrophage intermediates based on their high CLEC10A expression. The dashed line on the violin plots represents the median expression values for DC3/iDC3 for convenience of comparison with other myeloid cells. **(D)** UMAP illustrating CITE-seq data of myeloid cell clusters that remain after exclusion of FOLR2^pos^ STMs. **(E)** Heatmap illustrating differentially expressed (DE) surface markers in ST-DC3/iDC3 and ST-DC2 compared to other myeloid cell clusters (CITE-seq data). After exclusion of FOLR2^high^ STMs, CD32&CD39 are pinpointed as specific markers of ST myeloid DCs, in addition to CLEC10A. DE criteria: expressed in >40% of cells per cluster, with log-fold change >0.5, and p<0.05 based on MAST with Bonferroni correction for multiple comparison. **(F)** Gene/Protein expression density and violin plots on the UMAP from (D) illustrating that CLEC10A expression distinguishes ST-DC2 and ST-DC3/iDC3 from remaining myeloid cell clusters. Any remaining contamination with SPP1^pos^ macrophage cluster can be excluded by the high expression of CD9 protein and the lack of CLEC10A. **(G)** Violin plots illustrating that the expression of CD32&39, similarly to CLEC10A, distinguishes ST-DC3/iDC3 and ST-DC2 from any remaining contamination with other myeloid cells.

**Supplementary Figure 13.**
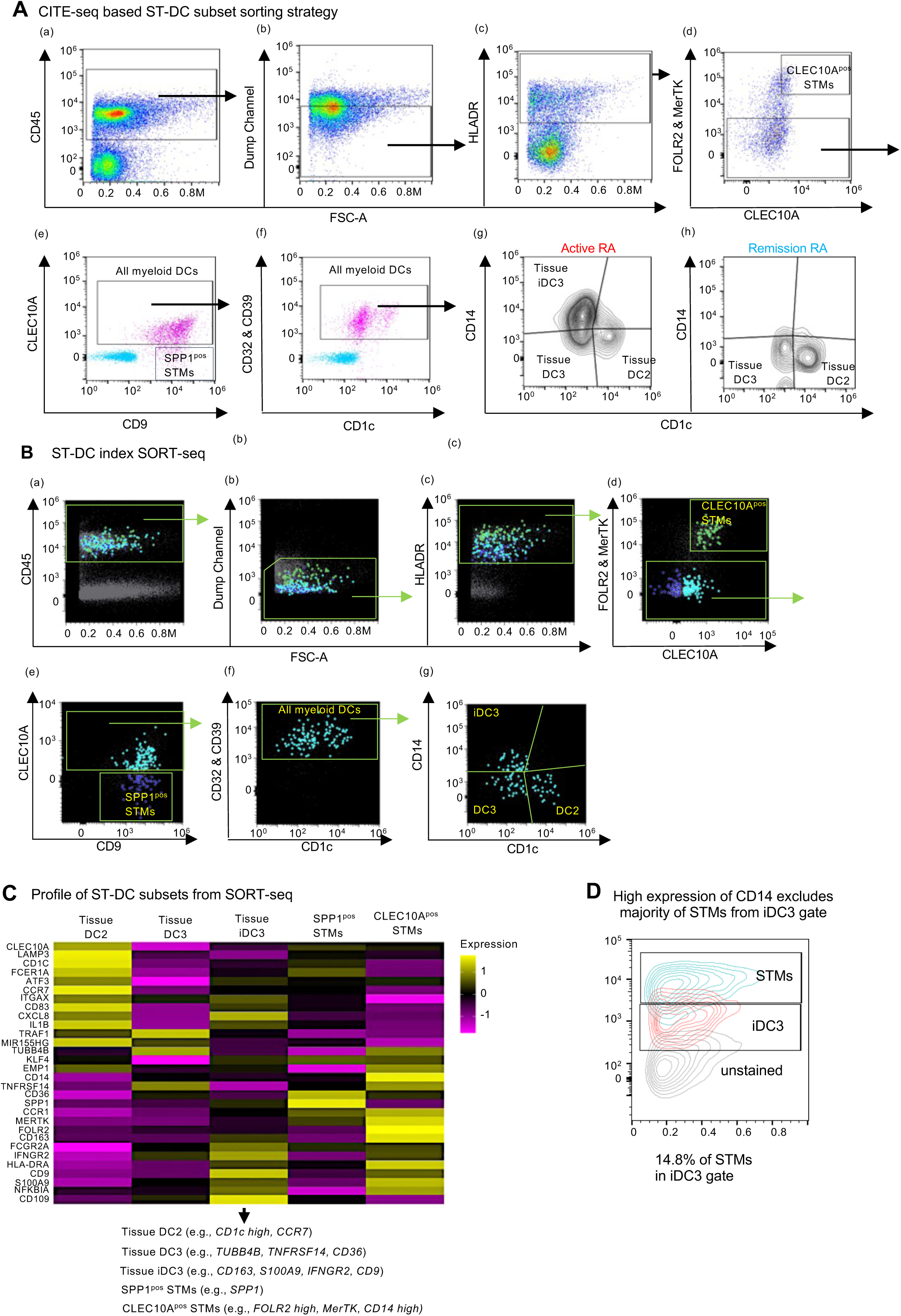
Validation of ST-DC gating strategy with ST-DC Index Single-Cell SORT-seq. **(A)** CITE-seq-based myeloid ST-DC subsets sorting strategy: Synovial tissue (ST) biopsies were digested as described in the Methods section. First, live cells were gated, followed by CD45-positive cells (a). In the next step, lineage-positive cells expressing CD3 (T cells), CD19 & CD20 (B cells), CD15 (neutrophils), CD117 (mast cells), and CD56 (NK & NKT cells) were excluded (Dump Channel) (b). Subsequently, cells expressing high levels of HLA-DR were gated (c). This was followed by gating cells negative for the synovial tissue macrophage markers FOLR2 & MerTK. Cells expressing FOLR2 and CLEC10A are CLEC10A^pos^ STMs (d). In the FOLR2 & MerTK negative gate, myeloid DCs were gated based on the expression of CLEC10A. The SPP1^pos^ STMs were excluded by the expression of CD9 and lack of expression of CLEC10A (e). To further exclude any remaining SPP1^pos^ STMs and CD14^pos^CD16^pos^ tissue monocytes from myeloid DCs, ST DC2/DC3/iDC3 were confirmed by high expression of CD32 & CD39 (f). The subsequent combination of CD1c and CD14 expression distinguished ST DC2, DC3, and iDC3 in active and remission (g-h). Unstained controls for CLEC10A, CD9, CD32 & CD39, and CD1c in (e) and (f) are shown in blue. **(B)** Confirmation of CITE-seq guided ST-DC subsets sorting strategy with ST-DC Index Single-Cell SORT-seq. Individual cells from the DC2, DC3, and iDC3 gates as in A, as well as from the SPP1^pos^ STM and CLEC10A^pos^FOLR^high^ STM gates (comparators), were sorted into a 364-well plate for subsequent single-cell sequencing. **(C)** Transcriptomic profiles of DC and macrophage subsets as defined by their sort-gate in B. Heatmap showing pseudobulk expression of DC subsets and macrophage subset markers based on this manuscript’s findings and published literature. **(D)** ST-DC index SORT-seq shows that the intermediate CD14 expression gate captures the majority of ST-iDC3, while high expression of CD14 excludes most STMs from the ST-iDC3 gate.

**Supplementary Figure 14.**
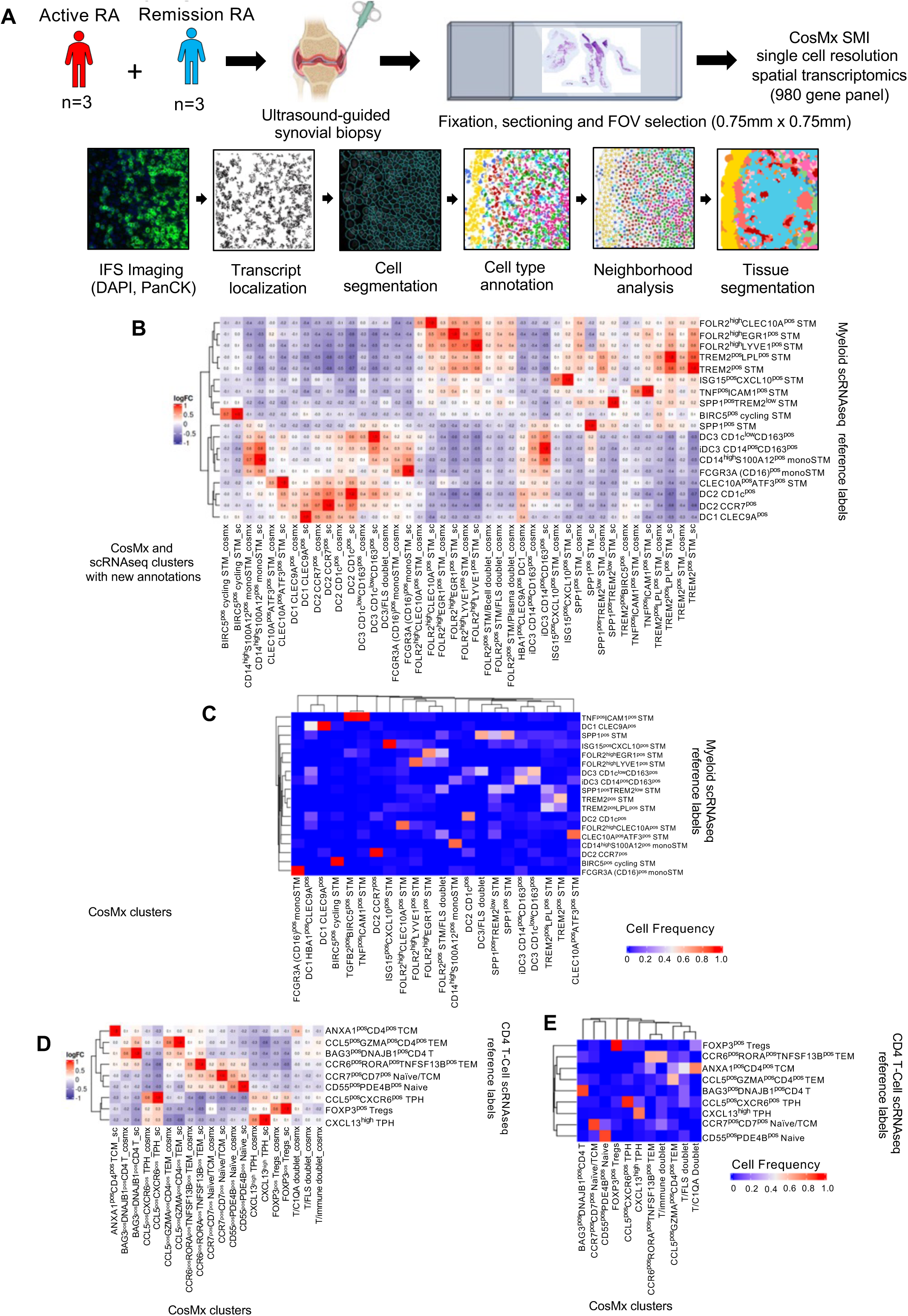
Annotation of myeloid cells and T-cells from CosMx spatial transcriptomic data with scRNAseq reference data of human synovium. **(A)** Workflow of single cell spatial transcriptomic experiment (CosMx) on synovial tissues. **(B)** Heatmap visualization of myeloid cells marker gene correlation between reference synovial tissue scRNAseq clusters (rows) and clusters identified from harmony integration of CosMx spatial transcriptomic and scRNAseq data (columns). **(C)** Confusion matrix showing the percentage of scRNAseq myeloid cell annotation (y-axis) compared with the new integrated clustering of scRNAseq and CosMx data. **(D)** Heatmap visualization of T-cells marker gene correlation between reference synovial tissue scRNAseq clusters (rows) and clusters identified from harmony integration of CosMx spatial transcriptomic and scRNAseq data (columns). **(E)** Confusion matrix showing the percentage of scRNAseq T-cell annotation (y-axis) compared with the new integrated clustering of scRNAseq and CosMx data.

**Supplementary Figure 15.**
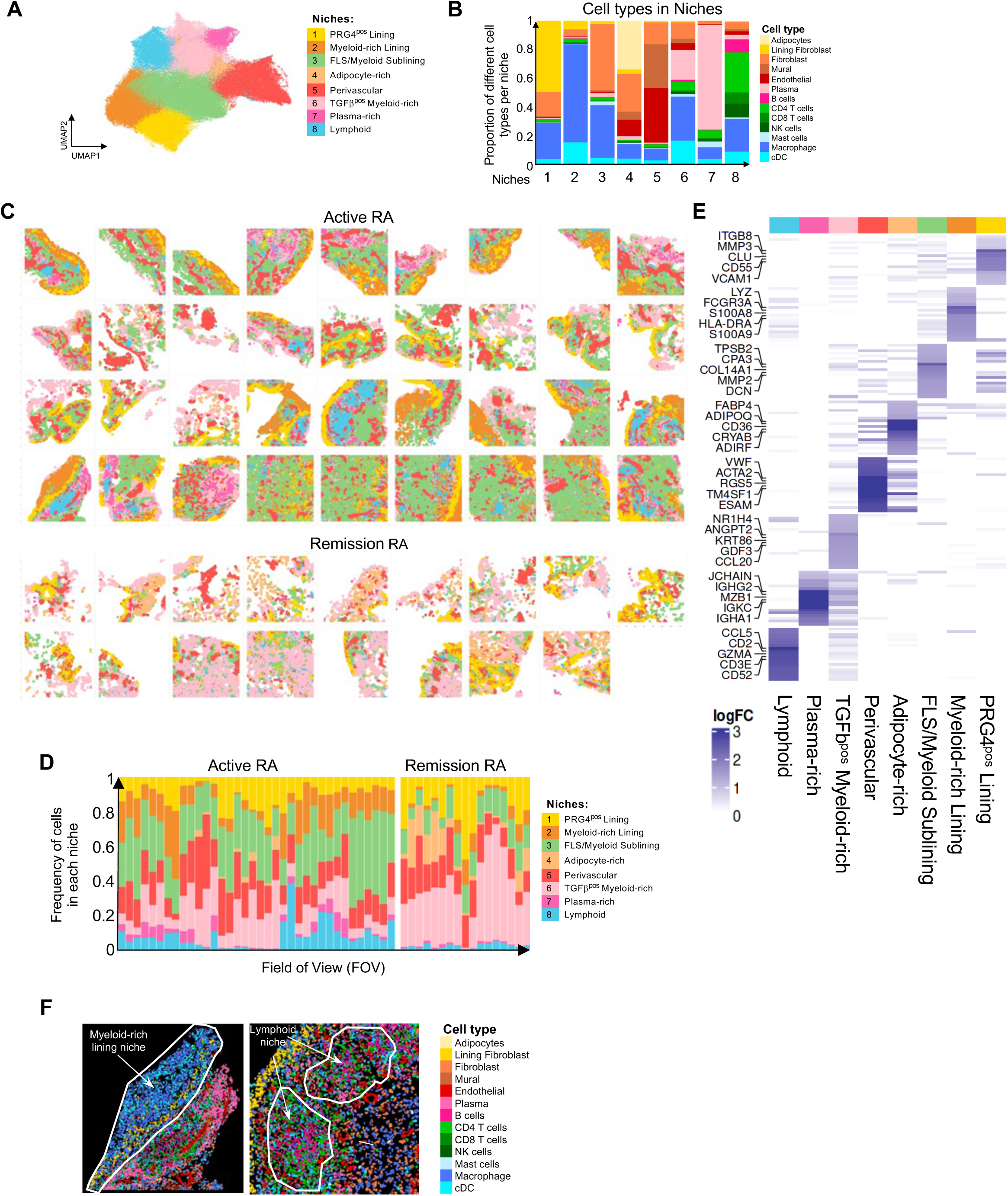
Identification of synovial tissue niches in active and remission RA CosMx spatial transcriptomic data. **(A)** Louvain clustering of spatial tiles (voronoi polygons) identified 8 unique tissue niches, as shown by UMAP visualization. **(B)** Stacked bar plots illustrating the relative distribution of different cell types in niches. **(C)** Overview of niche annotation in n=36 FOVs from active RA and n=17 FOVs from remission RA tissues. **(D)** Stack plot illustrating niche distribution in all FOVs analysed. **(E)** Heatmap visualization of the average log fold change (logFC) of the marker genes per niche, with top 5 genes annotated. Differential expression analysis performed using presto (1.0.0) for Generalised Linear Mixed Model (GLMM) estimation with lme4 (1.1-34). Genes were considered significant when adjusted p value was less than 0.01 and an average logFC more than 0.5. **(F)** Example of mapping of coarse cell type annotations in representative FOVs for niches of interest (Myeloid-rich lining (Left) and Lymphoid (right)).

**Supplementary Figure 16.**
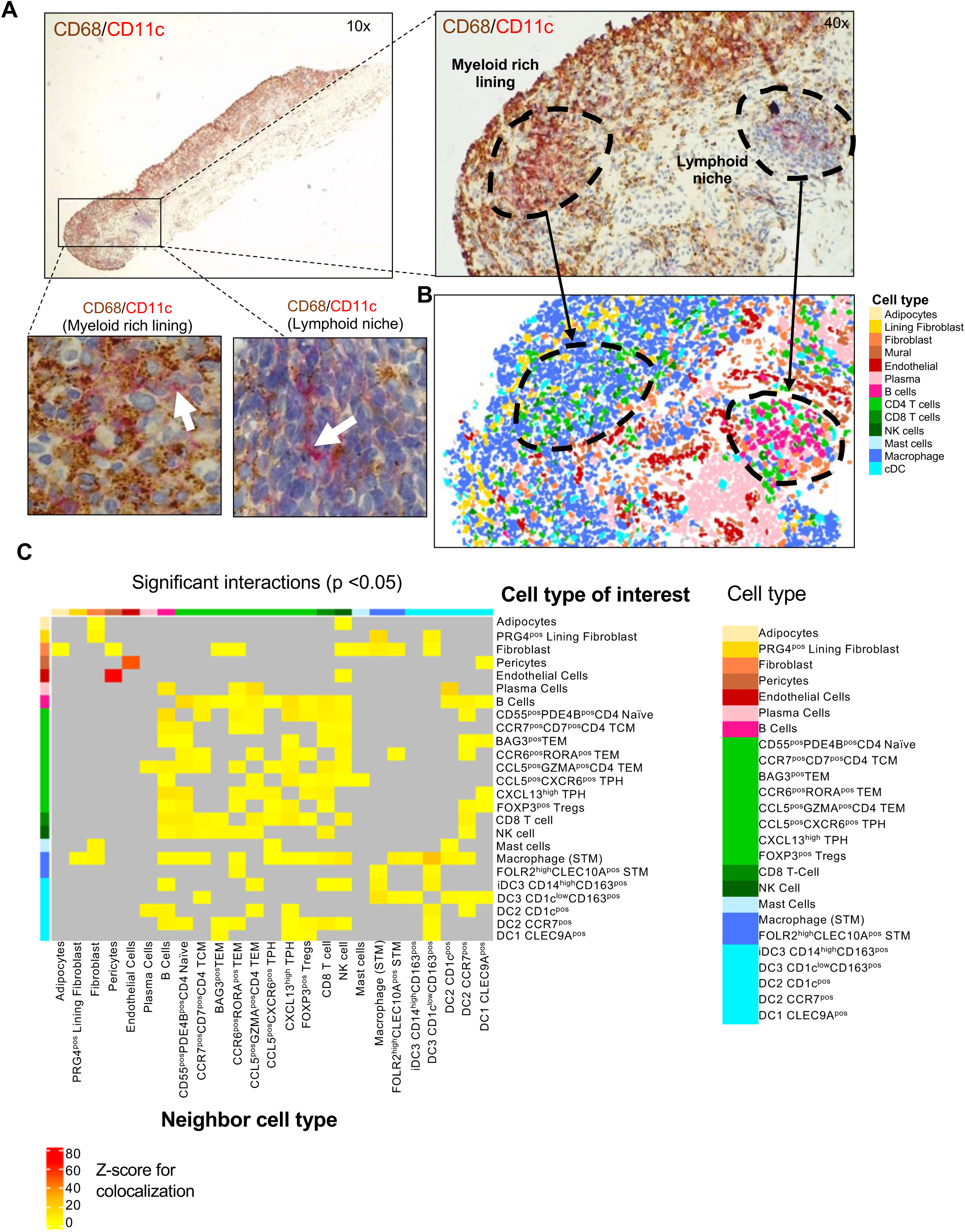
Illustration of the presence of myeloid DCs in synovial biopsies used in spatial transcriptomics and tissue cell colocalization analysis. **(A)** Representative images of CD11c^pos^ (pink) myeloid DCs in a synovial biopsy used in spatial transcriptomics. Synovial tissue stained with anti-CD68 (macrophage marker) and anti-CD11c (myeloid DC and some macrophage marker) antibodies show CD11c^pos^CD68^neg^ myeloid DCs (white arrows) in the lymphoid niche and hyperplastic myeloid-rich lining niche (Pink, CD11c staining and brown, CD68 staining). **(B)** Spatial transcriptomic deconvolution of section in A. **(C)** Neighbourhood analysis of direct interactions (40mm) between cells in the tissue. Colocalization with significant Z-scores/p-value is labelled with yellow-red colours, while insignificant ones are marked with grey (n=36 FOVs in active RA). DCs and CD4^pos^ T-cells are annotated to the level of specific clusters, while other cells are annotated by type.

**Supplementary Figure 17.**
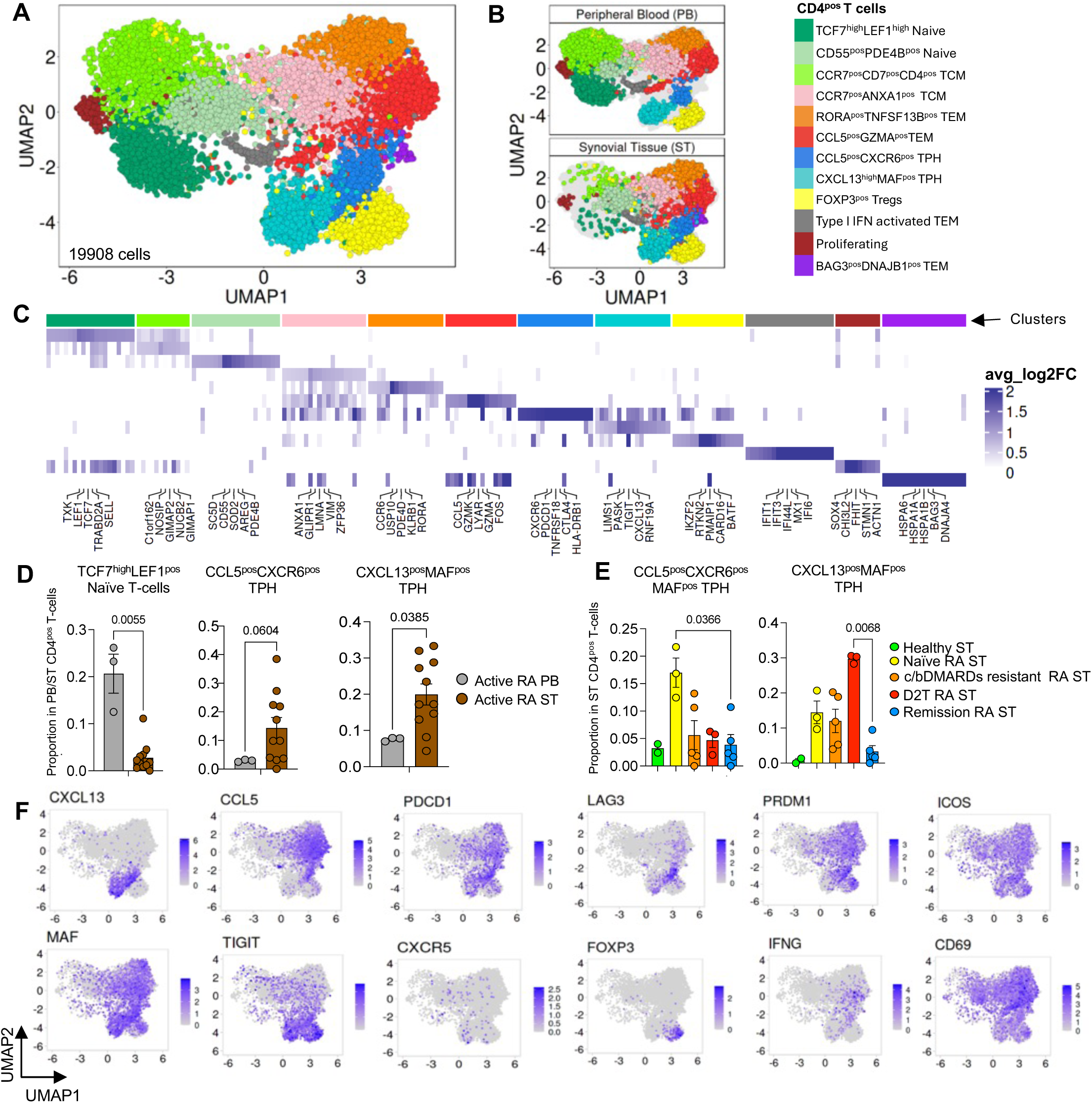
Distinct CD4^pos^ T-cell populations in the peripheral blood and synovial tissue of healthy, and patients with active RA and RA in sustained remission. **(A)** UMAP visualisation of scRNAseq data from healthy (n=3), active RA (n=12) and RA in remission (n=5) synovial tissue and from matched blood of active RA (n=3), identifying twelve CD4^pos^ T-cell clusters based on top marker genes. (B) Split UMAP visualising data from PB versus synovial tissue. **(B)** Heatmap illustrating cluster markers (expressed in >40% of cells/cluster, with log-fold change >0.25, and *p*<0.05 based on MAST with Bonferroni correction for multiple comparison). Differentially expressed marker genes annotated based on greatest average logFC per cluster. **(C)** Bar plots (mean ± SEM) showing relative proportion of naïve T-cells and two TPH clusters in peripheral blood and synovial tissue CD4^pos^ T-cell pool in active RA. Two-sided Mann-Whitney test, exact p-values are provided on the graphs. c/bDMARDs: Conventional or biological disease-modifying anti-inflammatory drugs, D2T: Difficult-to-treat (failed at least two biological therapies). **(D)** Bar plots (mean ± SEM) showing relative proportion of two TPH clusters in ST CD4^pos^ T-cell pool across different joint conditions. Each dot represents healthy donor/patient. One-way ANOVA with Dunn’s correction for multiple comparison, exact p-values are provided on the graphs. **(E)** Expression of TPH and CCL5^pos^ TEM markers in ST CD4^pos^ T-cell clusters.

**Supplementary Figure 18.**
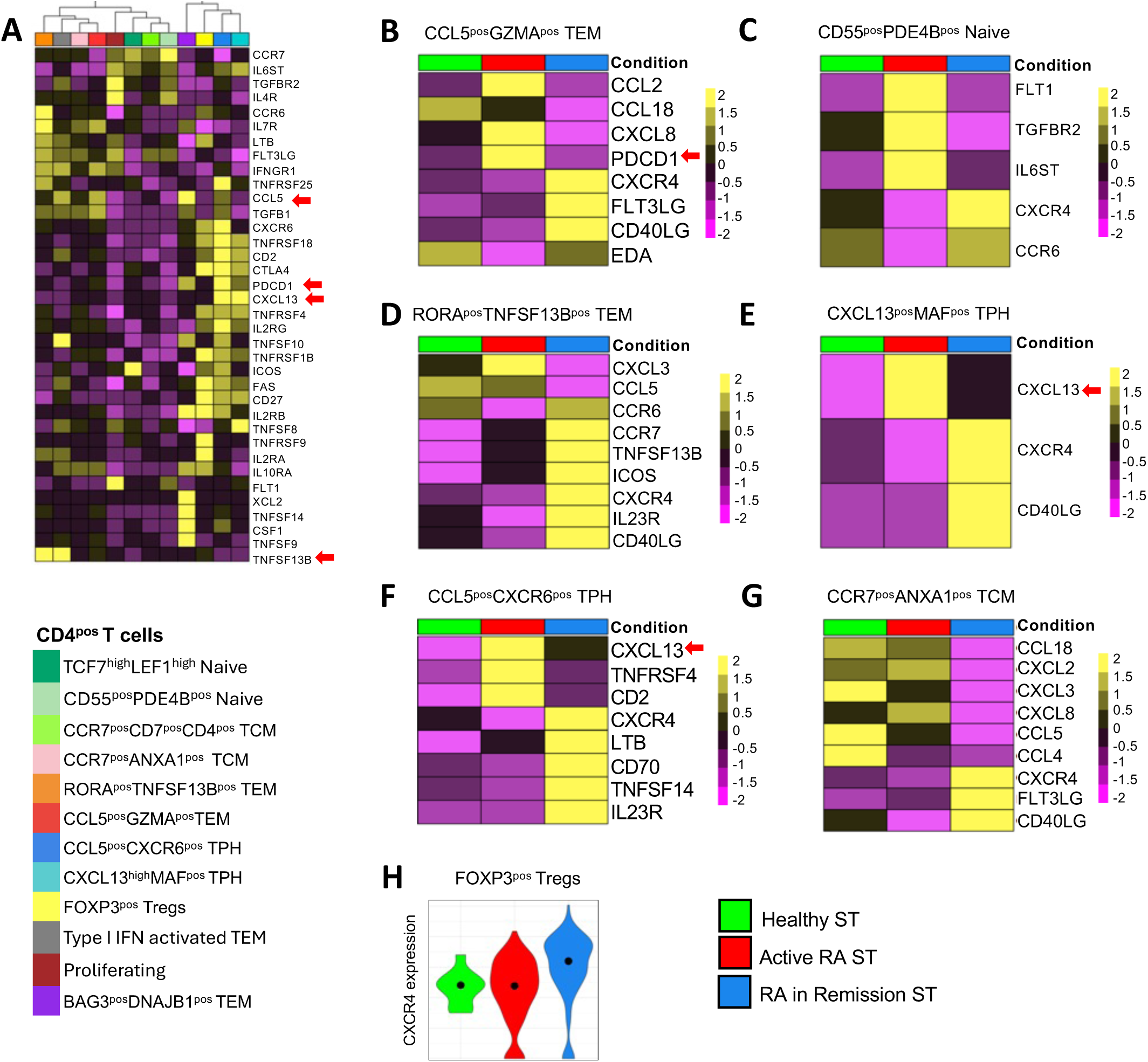
Cytokine and cytokine receptor profiles of synovial tissue CD4^pos^ T cells. **(A)** Heatmap visualising scaled differentially expressed genes from the KEGG cytokines and cytokine receptors pathway between different synovial tissue (ST) CD4^pos^ T-cell clusters (Criteria: expressed in >25% of cells per T-cell cluster, with log-fold change >0.5, and p<0.05 based on MAST with Bonferroni correction for multiple comparison). Data from healthy (n=3), active RA (n=12), and RA in remission (n=5) synovial tissue. **(B-H)** Heatmaps or Violin plot visualising scaled differentially expressed genes from the pathway as in A in different ST CD4^pos^ T-cell clusters between active RA (n=12) and remission RA (n=5). Criteria: expressed in >25% of cells per condition, with log-fold change >0.5, and p<0.05 based on MAST with Bonferroni correction for multiple comparison.

**Supplementary Figure 19.**
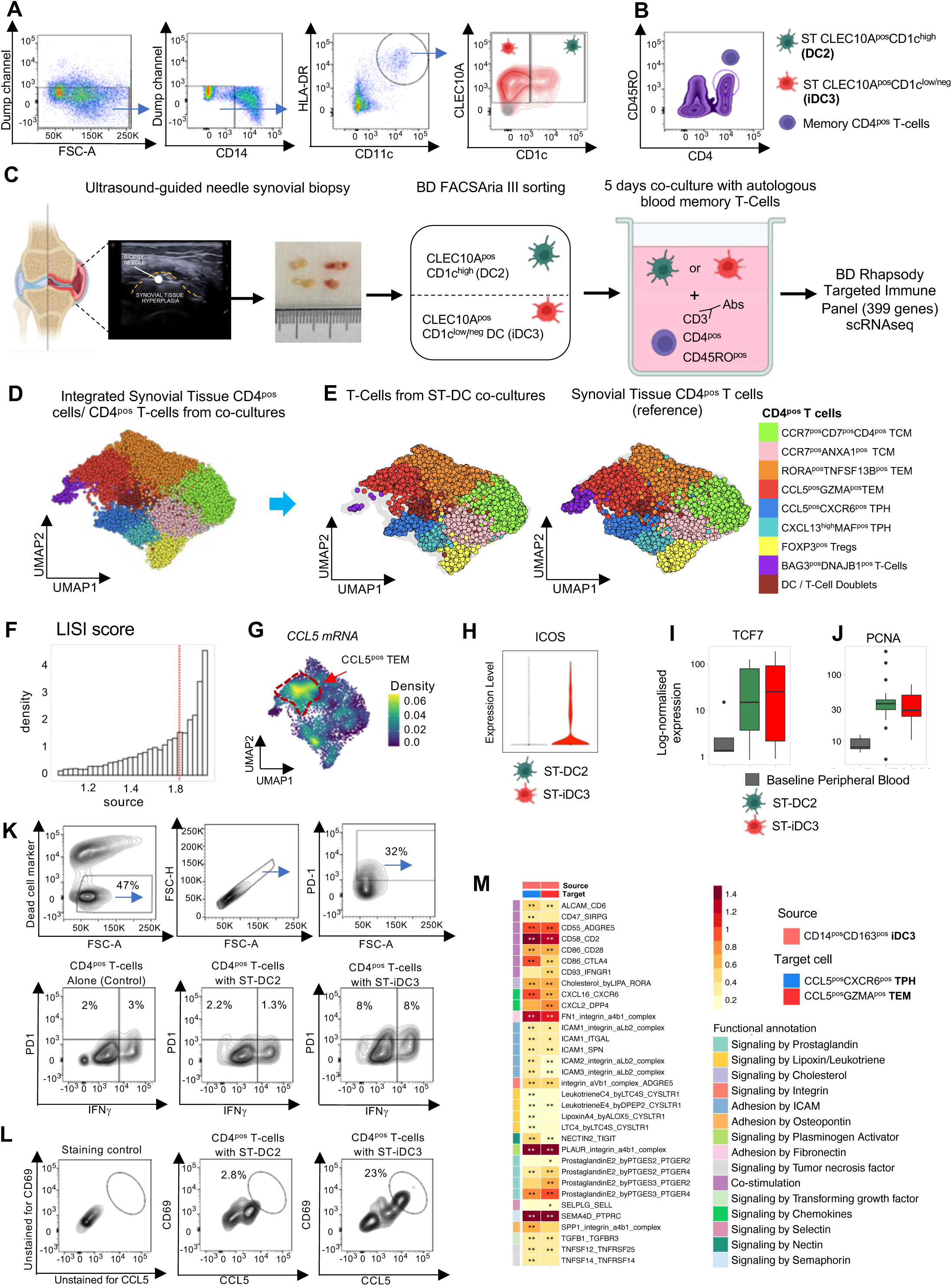
Synovial tissue iDC3 support CCL5^pos^TEM in ex-vivo co-culture with autologous CD4^pos^ memory T-cells. **(A)** Gating Strategy for Sorting ST Myeloid DC subsets. After exclusion of lineage-positive cells (CD3, CD19/20, CD15, CD117, and CD56 positive cells), synovial tissue macrophages were excluded based on high levels of CD14 expression (validated by ST-DC index cell sort, Figure S13C). Cells negative or expressing low/intermediate levels of CD14 were then taken forward, and cells expressing high levels of CD11c, and HLA-DR were gated. ST-DC2 were gated based on CLEC10A expression and high CD1c expression, while DC3/iDC3 were gated based on high CLEC10A expression and low/negative CD1c expression. Grey dots represent isotype controls. **(B)** Autologous memory T-cells were FACS-sorted from peripheral blood mononuclear cells based on the expression of CD3, CD4, and CD45RO. **(C)** Cells were co-cultured at a 1:5 ratio of ST-DC subset: memory T-cells in a 96-well, round-bottom cell culture plate. After 5 days in culture, cells were subjected to scRNAseq (BD Rhapsody Targeted Immune Panel). **(D)** UMAP visualization of CD4^pos^ T cells isolated after 5-day co-culture with ST-DCs (n=8 active RA) integrated with reference scRNAseq data from ST CD4^pos^ T-cells (n=12 active RA and n=5 from RA in remission). **(E)** Split UMAP visualization of CD4^pos^ T-cells from ST and co-culture annotated according to the reference ST dataset. **(F)** Histogram of the Local Inverse Simpson’s Index (LISI) score for integration of CD4^pos^ T-cells from ST and co-culture. LISI measures the degree of mixing amongst CD4 T-cells from coculture with those from reference synovial tissue dataset. Values reflect the number of different categories represented in the local neighbourhood of each cell and range from 1 (unmixed) to 2 (optimal mixing of in vitro or in vivo T cells). The median value for integration of CD4^pos^ T-cells from ST and co-culture is 1.82. (G) UMAP of ST T-cells (excluding doublets) illustrating that CCL5^pos^TEM express *CCL5*. (H) Violin plots visualising log-normalised expression of activation marker *(ICOS*) in T-cells from matched co-cultures with ST-DC2 or ST-DC3/iDC3 as in A-G. **(I, J)** Boxplots (median, IQR) illustrating the log-normalised expression per sample of TCF7 (I) and PCNA (J) between blood memory T-cells (baseline before co-culture with DCs) and matched T-cells from the co-cultures with ST-DC subsets. p< 0.01, Wald-test with Benjamini-Hochberg correction for multiple comparison. **(K)** An example of T-cell gating from the co-culture with ST-DCs sorted from synovial biopsies, according to the CITE-seq gating strategy presented in Figure S13A. Live cells were gated, followed by single cells, then PD1^pos^ T-cells. Intracellular IFN-γ protein staining in T-cells co-cultured with autologous ST-DC2 or ST-iDC3 in the presence of anti-CD3 abs (0.25mg/ml) and IL-15 (20ng/ml) or in T-cells cultured with anti-CD3 abs (0.25mg/ml) and IL-15 (20ng/ml) alone are shown. **(L)** Expression of CD69 activation marker on CCL5-producing T-cells from co-cultures with ST-DCs as in K is shown. **(M)** Heatmap showing the mean expression values of predicted statistically significant cellular interactions initiated by ST-iDC3 (CD14^high^CD163^pos^) (pink cluster) to receiver CCL5^pos^CXCR6^pos^MAF^pos^ TPH (red) and CCL5^pos^GZMA^pos^CD4^pos^ TEM (blue) cells in the synovial niche based on single cell data. Functional annotations of curated interactions are annotated where available. Significant interactions are denoted by an asterisk (** < 0.01, * for p <0.05).

**Supplementary Figure 20.**
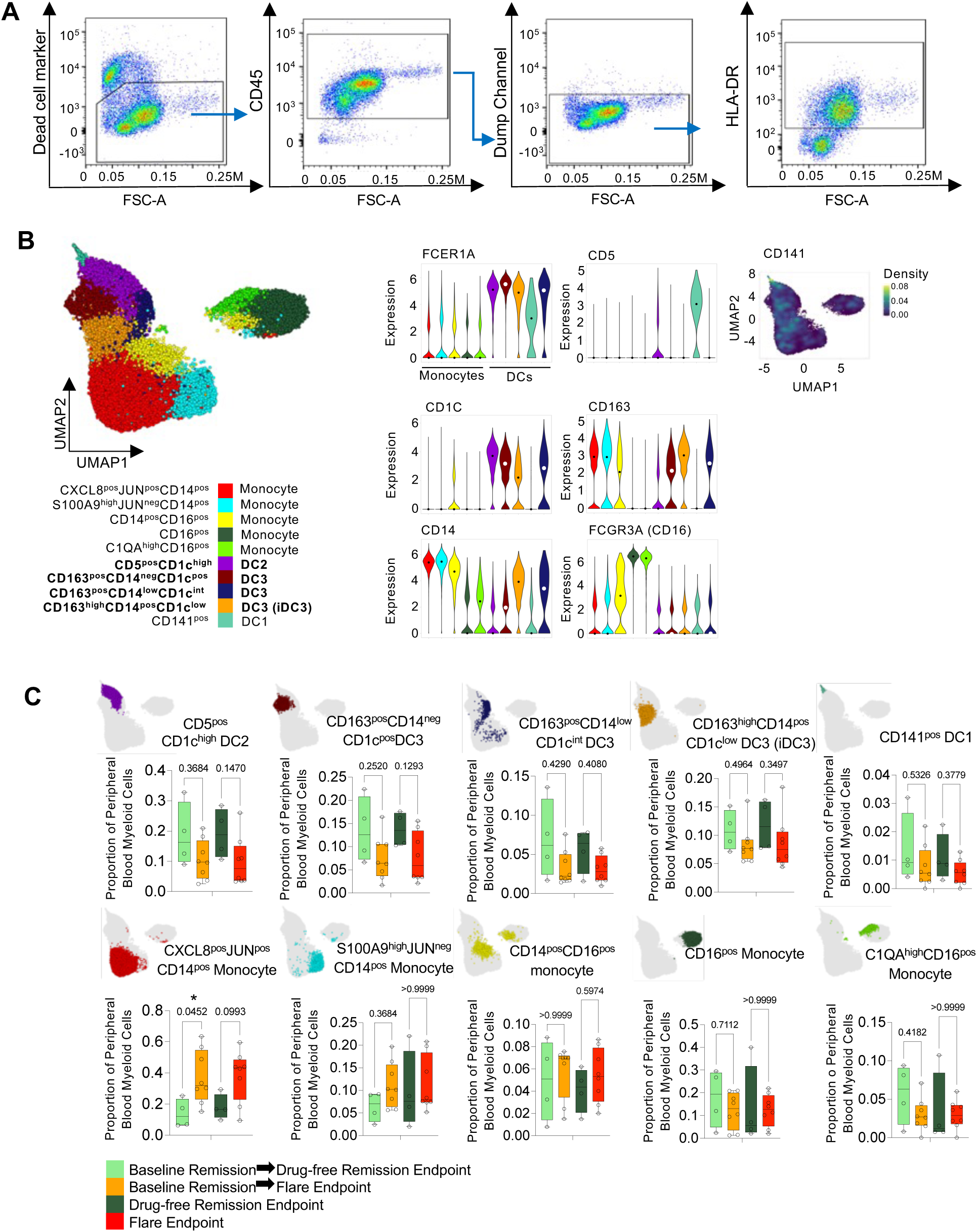
Frequency of blood monocyte and DC clusters in RA in disease remission before treatment withdrawal (baseline) and at subsequent clinical outcome (drug-free remission or flare). **(A)** Representative gating strategy for sorting peripheral blood myeloid cells from RA patients in disease remission for scRNAseq (Figure 6). Dead cells were excluded, then cells positive for CD45 were gated. CD3, CD19, CD15, CD117, and CD56 lineage positive cells were excluded (Dump channel). Remaining HLA-DR positive cells were sorted, and subject to scRNAseq using BD Rhapsody targeted immune panel. **(B)** Visualization of the expression of markers of PB monocytes and conventional DC subsets: DC1, DC2, DC3 and its three distinct clusters, defined by CD14 and CD163 expression. **(C)** Boxplots (median and IQR) illustrate relative proportions of different myeloid cell clusters and DC1 at baseline remission and at subsequent clinical outcome: maintained remission (n=4) or flare (n=8). Each dot represents an individual patient. Colour represents clinical outcome as in legend. Kruksal-Wallis adjusted for multiple comparisons with Dunn’s test. The exact p-values are on the graphs.

**Table S1.** Demographic, clinical and immunological characteristics of patients included in scRNAseq/CITEseq data sets.

|  | Healthy<br>n=7 | Active RA<br>n=18 | Remission<br>RA<br>n=9 | *p* |
| --- | --- | --- | --- | --- |
| **Female sex, n(%)** | 4 (57.1) | 17 (94.4) | 8 (88.9) | *0.60* |
| **Age, years (mean ± SEM)** | 34.7 ± 3.8 | 55.9 ± 3.10 | 56.8 ± 3.7 | *0.91* |
| **Disease duration, years (mean ± SEM)** | - | 3.0 ± 0.9 | 5.6 ± 0.8 | *0.0091* |
| **Disease Activity Score 28, (mean ± SEM)** | - | 5.7 ± 0.2 | 2.0 ± 0.1 | *<0.0001* |
| **ACPA positivity, n(%)** | - | 15 (83.3) | 6 (66.7) | *0.32* |
| **RF positivity, n(%)** | - | 11 (61.1) | 5 (55.5) | *0.78* |
| **Concomitant treatments:** | - |  | - | - |
| Naive, n(%) |  | 8 (44.4) |  |  |
| c/b-DMARDs resistant, n(%) |  | 10 (55.6) |  |  |
| **Total KSS, (mean ±SEM)** | - | 5.9 ± 0.5 | 2.4 ± 0.4 | *<0.0001* |
| **KSS sub-items:** |  |  |  |  |
| Synovial hyperplasia, (mean ±SEM) |  | 2.2 ± 0.2 | 0.8 ± 0.1 | *<0.0001* |
| Stromal density, (mean ±SEM) |  | 2.0 ± 0.1 | 0.9 ± 0.2 | *0.0002* |
| Inflammatory infiltrate, (mean ±SEM) |  | 2.1 ± 0.1 | 0.7 ± 0.3 | *<0.0001* |

**ACPA:** Anti-Citrullinated Peptide Antibodies; **RF:** Rheumatoid Factor; **c/b-DMARDs:** conventional/biologic-Disease Modifying Anti-Rheumatic Drugs; **KSS:** Krenn Synovitis Score; **SEM:** Standard Error of Mean.

**Table S2.** Demographic, clinical and immunological characteristics of Rheumatoid arthritis patients included in flow cytometry validation of synovial tissue DC subsets and ST-DC index SORT-plate.

|  | **Active RA**<br>n=15 | **Remission RA**<br>n=8 | *p* |
| --- | --- | --- | --- |
| **Female sex, n(%)** | 11 (84.6) | 6 (75.0) | *0.585* |
| **Age, years (mean ± SEM)** | 60.9 ± 3.9 | 50.1 ± 4.4 | *0.060* |
| **Disease duration, years (mean ± SEM)** | 3.7 ± 0.8 | 5.8 ± 0.9 | *0.099* |
| **Disease Activity Score, (mean ± SEM)** | 6.0 ± 0.3 | 1.2 ± 0.3 | *<0.0001* |
| **ACPA positivity, n(%)** | 8 (61.5) | 5 (62.5) | *0.964* |
| **RF positivity, n(%)** | 10 (76.9) | 4 (50.0) | *0.203* |
| **Concomitant treatments:** |  |  |  |
| Naive, n(%) | 4 (30.7) |  | - |
| c/b-DMARDs, n(%) | 9 (69.3) | 8 (100.0) | - |
| **Total KSS, (mean ±SEM)** | 6.2 ± 0.6 | 2.1 ± 0.3 | *0.0003* |
| ***KSS sub-items:*** |  |  |  |
| Synovial hyperplasia, (mean ±SEM) | 2.4 ± 0.2 | 0.6 ± 0.2 | *0.0004* |
| Stromal density, (mean ±SEM) | 1.6 ± 0.2 | 1.0 ± 0.0 | *0.0159* |
| Inflammatory infiltrate, (mean ±SEM) | 2.2 ± 0.2 | 0.5 ± 0.2 | *<0.0001* |

**ACPA:** Anti-Citrullinated Peptide Antibodies; **RF:** Rheumatoid Factor; **c/b-DMARDs:** conventional/biologic-Disease Modifying Anti-Rheumatic Drugs; **KSS:** Krenn Synovitis Score; **SEM:** Standard Error of Mean.

**Table S3:** Demographic, clinical and immunological characteristics of Rheumatoid arthritis patients included in mapping DCs in synovial tissue by immunofluorescent staining.

|  | Healthy<br>n=5 | Active RA<br>n=6 | Remission<br>RA<br>n=6 | *p* |
| --- | --- | --- | --- | --- |
| **Female sex, n(%)** | 2 (40.0) | 4(66.7) | 4(66.7) | *1.00* |
| **Age, years (mean ± SEM)** | 51.4 ± 4.5 | 60.3 ± 4.1 | 60.1 ± 5.1 | *0.89* |
| **Disease duration, years (mean ± SEM)** | - | 3.3 ± 1.1 | 5.5 ± 1.3 | *0.76* |
| **Disease Activity Score, (mean ± SEM)** | - | 5.8 ± 0.4 | 1.2 ± 0.2 | *<0.0001* |
| **ACPA positivity, n(%)** | - | 5 (83.3) | 4 (66.7) | *0.50* |
| **RF positivity, n(%)** | - | 5 (83.3) | 4 (66.7) | *0.50* |
| **Concomitant treatments:** | - |  |  |  |
| Naive, n(%) |  | 6(100.0) | - | - |
| c/b-DMARDs, n(%) |  |  | 6(100.0) | - |

**ACPA:** Anti-Citrullinated Peptide Antibodies; **RF:** Rheumatoid Factor; **c/b-DMARDs:** conventional/biologic-Disease Modifying Anti-Rheumatic Drugs; **KSS:** Krenn Synovitis Score; **SEM:** Standard Error of Mean.

**Table S4.** Demographic, clinical and immunological characteristics of RA patients whose synovial biopsies were used for spatial transcriptomics (CosMx).

|  | Active RA<br>n=3 | Remission RA<br>n=3 | *p* |
| --- | --- | --- | --- |
| **Female sex, n(%)** | 3(100.0) | 3(100.0) | 1.00 |
| **Age, years (mean ± SEM)** | 48.8 ± 3.9 | 52.2 ± 4.8 | 0.68 |
| **Disease duration, years (mean ± SEM)** | 1.3 ± 1.0 | 4.3 ± 1.2 | 0.005 |
| **Disease Activity Score, (mean ± SEM)** | 6.0 ± 0.4 | 1.3 ± 0.1 | <0.0001 |
| **ACPA positivity, n(%)** | 3 (100.0) | 3 (100.0) | 1.00 |
| **RF positivity, n(%)** | 3 (100.0) | 3 (100.0) | 1.00 |
| **Concomitant treatments:** |  |  |  |
| Naive, n(%) | 3(100.0) | - | - |
| c/b-DMARDs, n(%) |  | 3(100.0) | - |

**ACPA:** Anti-Citrullinated Peptide Antibodies; **RF:** Rheumatoid Factor; **c/b-DMARDs:** conventional/biologic-Disease Modifying Anti-Rheumatic Drugs; **KSS:** Krenn Synovitis Score; **SEM:** Standard Error of Mean.

**Table S5.** Demographic, clinical and immunological characteristics of healthy and RA patients included in evaluation of MIR155 expression.

|  | Healthy PB<br>n=12 | Active RA PB<br>n=16 | Active RA ST<br>n=5 |
| --- | --- | --- | --- |
| **Age, years (mean ± SEM)** | 51.5 ± 8.8 | 58.1±14.4 | 57.1 ± 14.4 |
| **Disease Activity Score, (mean ± SEM)** | - | 4.6 ± 0.2 | 5.37 ± 0.65 |

**SEM:** Standard Error of Mean.

**Table S6.**
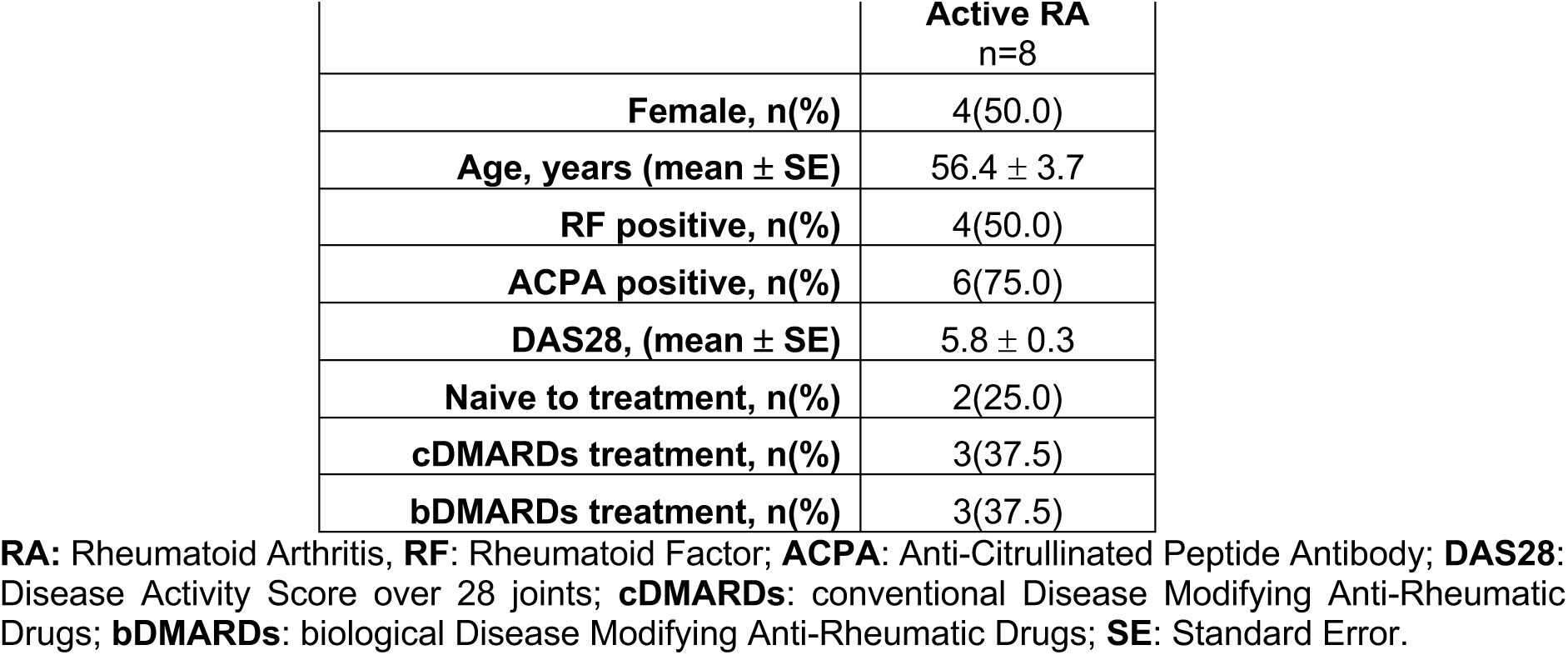
Demographic, clinical, and immunological characteristics of Rheumatoid Arthritis patients whose ST-DCs and PB T-cells were used in co-cultures and analysed with T-cell scRNA-seq readouts.

|  | **Active RA**<br>**n=8** |
| --- | --- |
| **Female, n(%)** | 4(50.0) |
| **Age, years (mean ± SE)** | 56.4 ± 3.7 |
| **RF positive, n(%)** | 4(50.0) |
| **ACPA positive, n(%)** | 6(75.0) |
| **DAS28, (mean ± SE)** | 5.8 ± 0.3 |
| **Naive to treatment, n(%)** | 2(25.0) |
| **cDMARDs treatment, n(%)** | 3(37.5) |
| **bDMARDs treatment, n(%)** | 3(37.5) |

**RA:** Rheumatoid Arthritis, **RF:** Rheumatoid Factor; **ACPA:** Anti-Citrullinated Peptide Antibody; **DAS28:** Disease Activity Score over 28 joints; **cDMARDs:** conventional Disease Modifying Anti-Rheumatic Drugs; **bDMARDs:** biological Disease Modifying Anti-Rheumatic Drugs; **SE:** Standard Error.

**Table S7.** Demographic, clinical and immunological characteristics of Rheumatoid Arthritis patients whose matched ST-DC subsets and PB T-cells were used in co-cultures, and T-cells analysed with intracellular cytokine readout.

|  | **Active RA**<br>**N=4** |
| --- | --- |
| **Female, n(%)** | 3 (75.0) |
| **Age, years (mean ± SE)** | 49.5 ± 8.2 |
| **RF positive, n(%)** | 2 (50.0) |
| **ACPA positive, n(%)** | 2 (50.0) |
| **DAS28, (mean ± SE)** | 5.4 ± 0.3 |
| **Concomitant treatments:** |  |
| **Naive, n(%)** | 1 (25.0) |
| **cDMARDs treatment n(%)** | 2 (50.0) |
| **bDMARDs treatment n(%)** | 1 (25.0) |
| **Total Krenn score (0-9) (mean ± SE)** | 4.5 ± 1.2 |
| **Synovial Hyperplasia (0-3) (mean ± SE)** | 1.8 ± 0.5 |
| **Stromal density (0-3) (mean ± SE)** | 1.5 ± 0.5 |
| **Inflammatory Infiltrate (0-3) (mean ± SE)** | 1.3 ± 0.5 |

**RA:** Rheumatoid Arthritis, **RF:** Rheumatoid Factor; **ACPA:** Anti-Citrullinated Peptide Antibody; **DAS:** Disease Activity Score; **cDMARDs:** conventional Disease Modifying Anti-Rheumatic Drugs; **bDMARDs:** biological Disease Modifying Anti-Rheumatic Drugs; **SE:** Standard Error.

**Table S8.** Demographic, clinical and immunological characteristics of patients with Rheumatoid Arthritis in Remission (BioRRA).

| Characteristic<br>(Baseline) | Clinical Outcome within 6 months<br>after treatment withdrawal |  |
| --- | --- | --- |
|  | Flare | Maintained<br>Remission |
| Number of patients | 8 | 4 |
| Female n (%) | 4 (50) | 2 (50) |
| Age, years (median, IQR) | 64 (50–71) | 62 (44–78) |
| Either ACPA or RhF positive n(%) | 8 (100) | 3 (75) |
| Years since diagnosis (median, IQR) | 6 (4–10) | 2 (2–3) |
| Months since last change in DMARD therapy (median, IQR) | 28 (16–36) | 15 (9–26) |
| Months since last glucocorticoid use (median, IQR) | 34 (24–39) | 22 (12–34) |
| DAS28-CRP at enrolment (median, IQR) | 1.05 (0.98–1.69) | 1.48 (1.04–2.00) |
| ACR/EULAR Boolean remission at enrolment n(%) | 6 (75) | 4 (100) |
| Methotrexate use at enrolment n(%) | 8 (100) | 3 (75) |
| Sulfasalazine use at enrolment n(%) | 3 (38) | 2 (50) |
| Hydroxychloroquine use at enrolment n(%) | 2 (25) | 0 (0) |
| DAS28-CRP at flare (median, IQR) | 3.94 (2.86–4.38) | n/a |

**Table S9.**
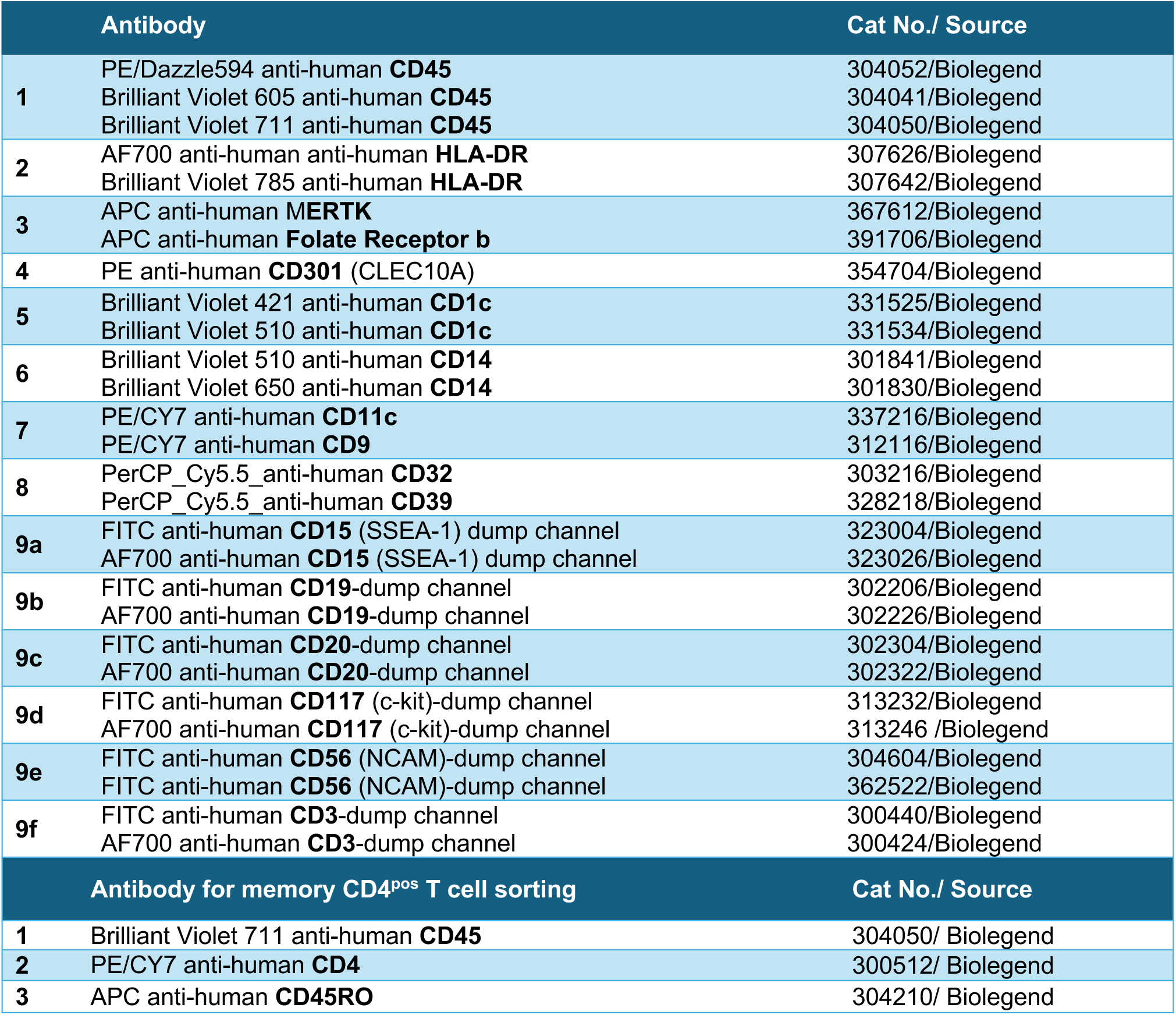
Details of antibodies used for flow cytometry phenotyping, Plate SORT-seq and ST-DC and PB memory CD4^pos^ T cells sorting for co-cultures.

**Table S10.**
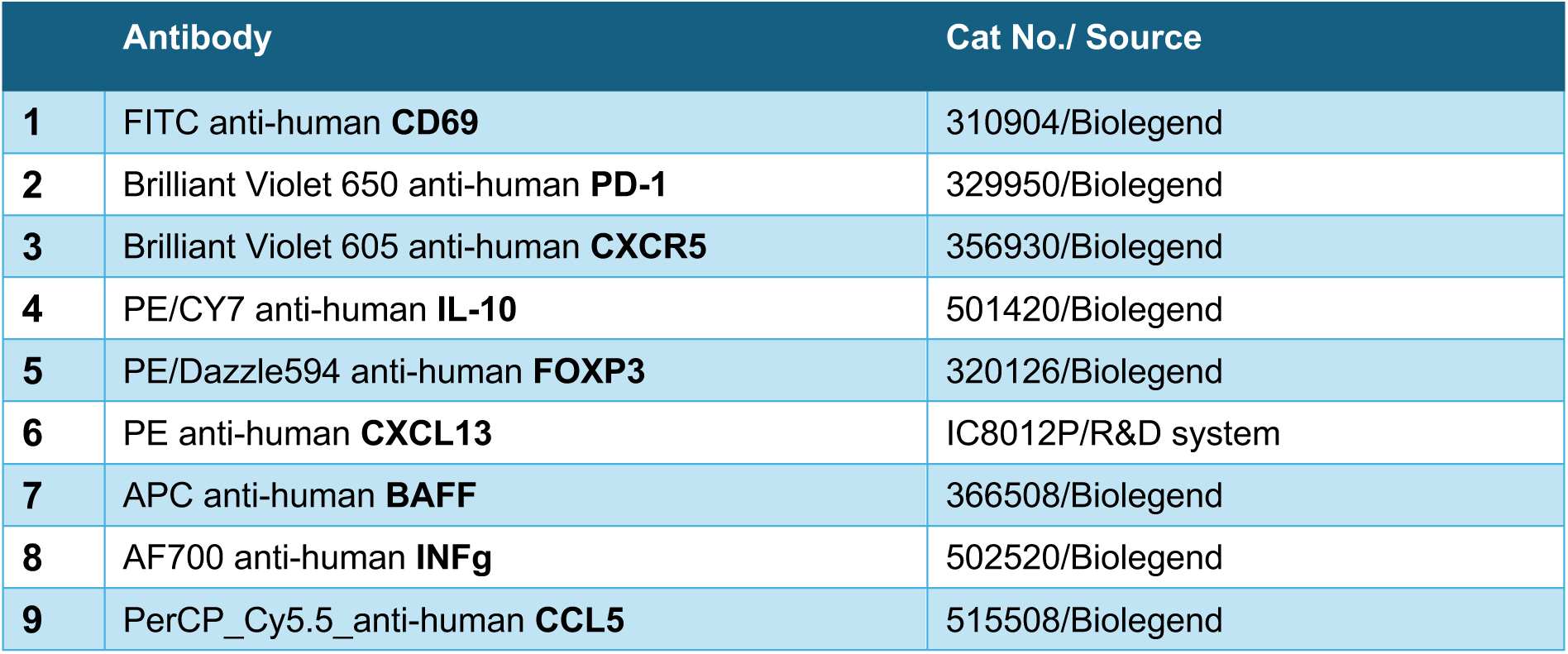
Details of antibodies used to evaluate PB memory CD4^pos^ T cell phenotype after co-culture with ST-DC subsets using flow cytometry.

**Table S11.** Primary and secondary antibodies used to map distinct DCs in human synovial tissues by immunofluorescent staining.

| Primary Antibodies | Working Dilution | Cat No./ Source | Secondary antibodies |
| --- | --- | --- | --- |
| Rabbit anti-human **CLEC10A** | 1:100 | Ab197346/Abcam plc | A-11008/ Goat anti-Rabbit IgG Alexa Fluor 488 (1:100) |
| Rabbit anti-human **LAMP3** | 1:100 | AF154/R&D system | A-21074/ Goat anti-Rabbit IgG Alexa Fluor 660 (1:100) |
| Rabbit anti-human **AXL** | 1:200 | Ab219651/Abcam plc | A-21074/ Goat anti-Rabbit IgG Alexa Fluor 660 (1:100) |
| Mouse anti-human **CD68** | 1:40 | M087629-2/ Dako, Cambridgeshire, UK | A-21055/ Goat anti-Mouse IgG Alexa Fluor 660 (1:100) |
| Mouse anti-human **CD1c** | 1:100 | Ab156708/ Abcam plc | A-21202/ Goat anti-Donkey IgG Alexa Fluor 448 (1:100) |

**Table S12.** Details of antibodies used for sorting myeloid cells from BioRRA PBMC samples.

|  | **Antibody** | **Titration** | **Cat No./ Source** |
| --- | --- | --- | --- |
| **1** | BUV395 anti-human **CD8** | 1:20 | 563795/BD |
| **2** | BUV737 anti-human **CD19** | 1:100 | 741829/BD |
| **3** | Brilliant Violet 421 anti-human **PD-1** | 1:10 | 564323/BD |
| **4** | Brilliant Violet 510 anti-human **CD3** | 1:20 | 300448/Biolegend |
| **5** | PE/CY7 anti-human **CD45RO** | 1:20 | 304230/Biolegend |
| **6** | AF700 anti-human **CD4** | 1:100 | 344622/Biolegend |
| **7** | PerCP_Cy5.5_anti-human **CD15** | 1:10 | 323020/Biolegend |
| **8** | PerCP_Cy5.5_anti-human **CD66b** | 1:10 | 305108/Biolegend |
| **9** | PerCP_Cy5.5_anti-human **CD203c** | 1:10 | 324608/Biolegend |

**Table S13.** Details of parameters used in integration and clustering of all single-cell and spatial transcriptomic data.

| *Dataset* | *Platform* | *Total cells* | *Norm. Scale Factor* | *Harmony Integration Variables* | *Harmony Theta* | *PCs* | *Clustering Resolution* | *Figure* | *Harmony Version* |
| --- | --- | --- | --- | --- | --- | --- | --- | --- | --- |
| *All cells from PB and ST* | 10X Genomics, BDRhapsody | 143,851 | 10,000 | sample donor, experiment, origin and sequencing platform | 0, 0, 1, 0 | 40 | 0.5 | Supp. Figure 1 | 0.1.0 |
| *Myeloid cells from PB and ST* | 10X Genomics, BDRhapsody | 70,471 | 10,000 | sample donor, experiment, origin and sequencing platform | 0, 0, 1, 0 | 40 | 1.5 | Supp. Figure 2 | 0.1.0 |
| *All myeloid DCs, CLEC10A STM and PB monocytes* | 10X Genomics, BDRhapsody | 37,725 | 10,000 | sample donor, experiment, origin and sequencing platform | 0, 1, 1, 2 | 20 | 2 | Supp. Figure 3 | 0.1.0 |
| *All myeloid DCs* | 10X Genomics, BDRhapsody | 7,869 | 10,000 | sample donor, experiment, origin and sequencing platform | 0, 2, 0, 2 | 20 | 1.5 | Figure 1 | 0.1.0 |
| *All CD4 T cells from PB and ST* | 10X Genomics | 19,908 | 10,000 | sample donor, experiment, tissue | 0, 10, 10 | 40 | 0.6 | Figure 4 | 1.1.0 |
| *CD4 T cells from ST with in vitro coculture* | 10X Genomics, BDRhapsody Targeted Immune Panel | 13,561 | 270.5 | cell source, sample donor, co-culture media | 5, 0, 5 | 20 | 1 | Supp. Figure 17 | 1.2.0 |
| *spatial transcriptomics data* | CosMx 1000 plex | 127,199 | NA | sample run, sample FOV | 0, 0 | 40 | 1.5 | Figure 3 | 0.1.0 |
| *ST myeloid cells from scRNAseq and spatial transcriptomics* | 10X Genomics, CosMx | 57,336 | 267 | source, sample donor | 2, 0 | 20 | 3.5 | Supp. Figure 13 | 0.1.0 |
| *ST stromal and endothelial cells from scRNAseq and spatial transcriptomics* | 10X Genomics, CosMx | 59,777 | 307 | source, sample donor | 2, 0 | 20 | 2.5 | Supp. Figure 13 | 0.1.0 |
| *ST lymphoid cells from scRNAseq and spatial transcriptomics* | 10X Genomics, CosMx | 8,787 | 219 | source, sample donor | 2, 0 | 20 | 2 | Supp. Figure 13 | 0.1.0 |
| *Integration of PB from Biorra Cohort* | BDRhapsody Targeted Immune Panel | 24, 711 | 10,000 | sample donor_timepoint | 2 | 30 | 1 | Figure 6 | 0.1.0 |
| *Integration with MNPverse* | 10X Genomics, BDRhapsody, SmartSeq2 | 186,466 | 4,680 | Unique_ID,Experiment,Tissue | 0,10,0 | 40 | 0.5 | Supp. Figure 10 | 0.1.0 |
| *Integration of CD4 T-Cells, CD8 T-Cells and NK cells from PB and ST* | 10X Genomics | 39,375 | 10,000 | sample donor, experiment and origin | 0, 2, 1 | 40 | 1.3 | N/A: Included as CD4 T-Cells were subsetted from this integration | 0.1.0 |

**Table S14.**
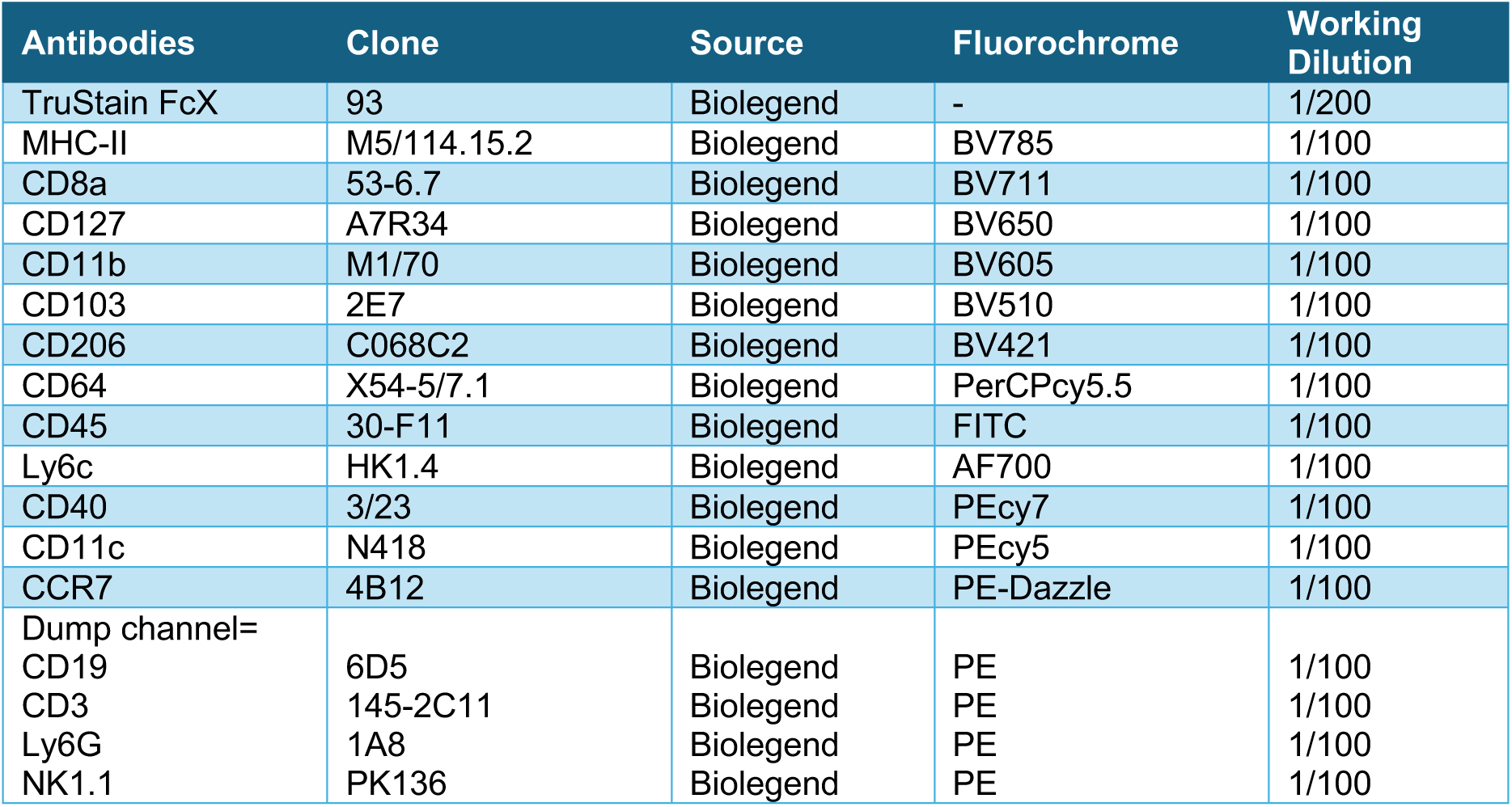
Details of antibodies used for evaluation of mouse CCR7^pos^ DC2 migrating from tissue to DLN.

| Antibodies | Clone | Source | Fluorochrome | Working Dilution |
| --- | --- | --- | --- | --- |
| TruStain FcX | 93 | Biolegend | - | 1/200 |
| MHC-II | M5/114.15.2 | Biolegend | BV785 | 1/100 |
| CD8a | 53-6.7 | Biolegend | BV711 | 1/100 |
| CD127 | A7R34 | Biolegend | BV650 | 1/100 |
| CD11b | M1/70 | Biolegend | BV605 | 1/100 |
| CD103 | 2E7 | Biolegend | BV510 | 1/100 |
| CD206 | C068C2 | Biolegend | BV421 | 1/100 |
| CD64 | X54-5/7.1 | Biolegend | PerCPcy5.5 | 1/100 |
| CD45 | 30-F11 | Biolegend | FITC | 1/100 |
| Ly6c | HK1.4 | Biolegend | AF700 | 1/100 |
| CD40 | 3/23 | Biolegend | PEcy7 | 1/100 |
| CD11c | N418 | Biolegend | PEcy5 | 1/100 |
| CCR7 | 4B12 | Biolegend | PE-Dazzle | 1/100 |
| Dump channel= |  |  |  |  |
| CD19 | 6D5 | Biolegend | PE | 1/100 |
| CD3 | 145-2C11 | Biolegend | PE | 1/100 |
| Ly6G | 1A8 | Biolegend | PE | 1/100 |
| NK1.1 | PK136 | Biolegend | PE | 1/100 |

